# Reconstructing Waddington’s Landscape from Data

**DOI:** 10.1101/2025.08.11.669575

**Authors:** Dillon J. Cislo, M. Joaquina Delás, James Briscoe, Eric D. Siggia

**Affiliations:** Center for Studies in Physics and Biology, Rockefeller University, New York, NY 10065, USA; The Francis Crick Institute, Midland Road, London, NW1 1AT, UK; Laboratory for Molecular Cell Biology, University College London, Gower Street, London WC1E 6BT, UK

## Abstract

The development of a zygote into a functional organism requires that this single progenitor cell gives rise to numerous distinct cell types. Attempts to exhaustively tabulate the interactions within developmental signaling networks that coordinate these hierarchical cell fate transitions are difficult to interpret or fit to data. An alternative approach models the cellular decision-making process as a flow in an abstract landscape whose signal-dependent topography defines the possible developmental outcomes and the transitions between them. Prior applications of this formalism have built landscapes in low-dimensional spaces without explicit reference to gene expression. Here, we present a computational geometry framework for fitting dynamical landscapes directly to high-dimensional single-cell data. Our method models the time evolution of probability distributions in gene expression space, enabling landscape construction with minimal free parameters and precise characterization of dynamical features, including fixed points, unstable manifolds, and basins of attraction. We demonstrate the applicability of this framework to multicolor flow-cytometry and RNA-seq data. Applied to a stem cell system that models ventral neural tube patterning, we recover a family of morphogen-dependent landscapes whose valleys align with canonical neural progenitor types. Remarkably, simple linear interpolation between landscapes captures signaling dependence, and chaining landscapes together reveals irreversible behavior following transient morphogen exposure. Our method combines the interpretability of landscape models with a direct connection to data, providing a general framework for understanding and controlling developmental dynamics.

---

Biology now possesses unprecedented capability to generate genome-wide readouts of cells as they develop in embryos. It remains a challenge to parse these data into comprehensible models that elucidate the underlying principles of development and move us towards the goal of engineering target cell and tissue fates. The canalized structure of developmental programs and their robustness to perturbations offer a route forward. Hearkening back to Waddington’s “epigenetic landscapes” [1], the time course of the collective state of a developing system can be conceptualized as downhill motion in an abstract landscape, where valleys correspond to developmental outcomes (Fig. 1). Building a framework that can accurately model developmental programs as flows in landscapes whose topographies are fit directly to data promises incisive explanatory and predictive power to both understand and manipulate development.

**Figure 1.**
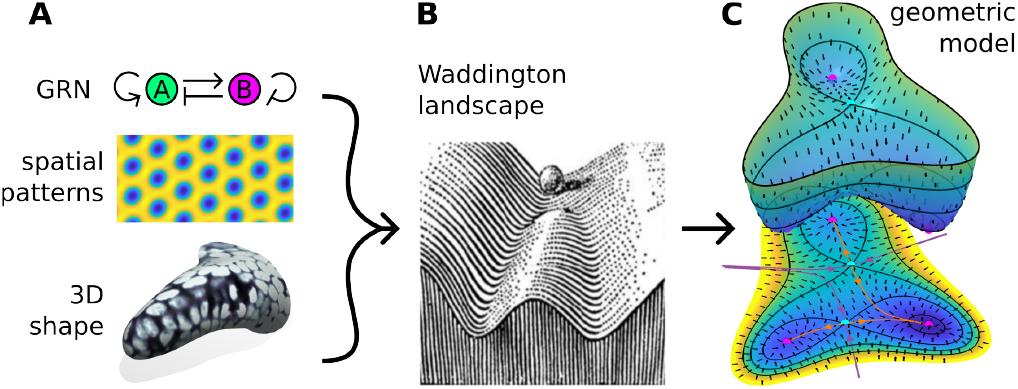
Geometric methods produce rigorous mathematical models of development in terms of a small set of phenomenological variables. The coarse-grained state of a developing system, be it gene expression levels or even the shape of an entire limb (A), can be conceptualized as valleys in an abstract landscape (B) (modified from [1]). This picture can be formalized through the introduction of a potential function and a metric tensor that fully describe the trajectories of developmental programs as they unfold in time (C).

Waddington’s landscape metaphor can be made mathematically rigorous when cast as a Morse-Smale (MS) dynamical system [2, 3]. These are flows that converge for long times to a finite number of rest points (saddles or attractors) and periodic orbits. Crucially, MS systems are structurally stable – small perturbations to the system’s parameters do not change its qualitative features. Such systems exclude chaos, guarantee robustness, and admit a Lyapunov-like function that decreases along every trajectory [2, 4]. Absent periodic orbits and provided some technical assumptions, Smale demonstrated that a MS system can be explicitly formulated as gradient-like [5], with a scalar potential defining landscape topography and a metric tensor that reorients the downhill motion. In this context, cell states correspond to equilibria. Stem-like progenitor states, represented by shallow attractors on high-up plateaus in the landscape, are connected via a sequence of lineage decisions at saddles to the attractors in valleys that represent terminally differentiated states. Lineage decisions emerge as bifurcations – minimal parameter changes that create, annihilate, or reconnect these fixed points. We expect that the generic bifurcations realized by tuning only one parameter (e.g. folds and flips [3]) dominate developmental decision spaces such that the diversity of cell fates can be encoded in a branching tree with bifurcations at the nodes.

These results unquestionably apply to systems of equations, but their relevance to real biological systems requires demonstration. Development proceeds through sequential, context-dependent lineage decisions, cells experience noise and signaling delays, and experimental readouts capture discrete time points rather than continuous trajectories. Nevertheless, enumerating every transcript and protein as a large system of coupled differential equations is computationally prohibitive and often biologically opaque. Instead, a landscape-based approach provides a principled and significant reduction in complexity, absorbing biochemical details into a topography that remains predictive yet interpretable. Prior work has shown that geometric methods can quantitatively predict the dynamics of developmental phenotypes in low-dimensional, phenomenological “fate spaces” in a variety of settings, including *C. elegans* vulval patterning [6–8], bristle patterning in *Drosophila* [9], and *in vitro* stem cell systems [10].

This work introduces a computational framework that can fit MS compatible landscapes directly to single-cell measurements, working directly in gene expression space. Rather than simulating individual cells and or cell-type proportions, we reconstruct the full time evolution of the probability distribution over gene expression space. We develop a novel way of modeling dynamic data from a series of population samples based on the path integral formulation of the Fokker-Planck equation [11], applicable in the high-dimensional spaces typical of data. It achieves an accuracy comparable to that of traditional finite-difference methods in low-dimensional spaces. Our discrete operators allow us to directly infer the dynamical structure of a system, including fixed points, unstable manifolds, and basins of attraction, with minimal preprocessing.

We apply our methods to model a mouse embryonic stem cell (mESC) system induced to mimic the patterning of the ventral neural tube [12, 13]. This setting is often reduced to the textbook “French-Flag” scheme, in which a static morphogen gradient assigns fates via a one-to-one mapping, yet *in vivo* measurements show hysteresis [14], indicating a more intricate underlying landscape with bistability. *In vitro*, the absence of cell-cell coupling allows us to isolate this landscape for an individual cell. Snapshots of gene expression profiles, captured over time using multicolor flow cytometry analysis and single-cell RNA-seq, reveal the evolving state distribution, enabling us to infer a potential that depends smoothly on morphogen dose. The resulting landscape reproduces observed fate proportions, interpolates to untested morphogen levels, and predicts responses to arbitrary morphogen histories. An examination of other methods for analyzing single-cell transcriptomics that evoke a landscape metaphor (e.g. [15–17]) is provided in the discussion.

## Algorithm

### Mathematical preliminaries

Our goal is to build a method for simulating the high-dimensional dynamics of a developing biological system that can be fit directly to experimental data. The application developed here is to single-cell data, but the formalism is broadly applicable to different types of developmental dynamics. Let **x**(*t*) ∈ ℝ^*d*^ denote the time-dependent state of a cell (e.g., the expression levels of *d* different genes at time *t*). We assume that the time evolution of the system is governed by the following Langevin equation:

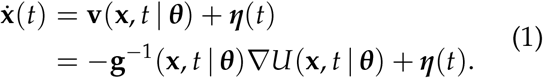

Here, the drift term **v** = − **g**^*−*1^ ∇*U* is assumed to be gradient-like for a given metric tensor **g** and potential *U*, which serves as the Lyapunov function in the Morse-Smale system we are modeling. The stochastic term ***η***(*t*) is standard white noise with mean ⟨***η***(*t*)⟩ = 0 and covariance

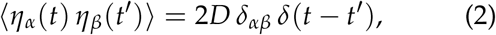

where Greek indices run over the dimensions 1, …, *d* and *D* is the (positive) diffusion constant. The biological systems we model are typically not stationary, and there is no explicit link between *U* in Eq. (1) and any equilibrium distribution (by design **g** does not appear in Eq. (2)).

We denote by ***θ*** a vector of time-dependent parameters that control the topography of the dynamical landscape (e.g. the expression levels of a set of morphogens at time *t*). Changing these parameters can destabilize existing states, introduce new states, or bias the system towards a particular outcome. We usually omit the explicit dependence on ***θ*** for the sake of notational simplicity. In the deterministic limit, the trajectories **x**(*t*) may evolve along a low-dimensional submanifold ℳ ⊂ ℝ^*d*^, but high-dimensional noise will generically blur this confinement. In this work, we do not attempt to explicitly encode the geometry of ℳ and instead extend the stochastic dynamics to ℝ^*d*^ by defining a *U*(**x**) that keeps trajectories close to ℳ.

It is not usually possible to continuously follow the state of a particular cell **x**(*t*) over time due to the destructive nature of experimental methods. A typical experiment will simultaneously measure the state of many cells at a particular time *t* subject to experimental conditions ***θ***. This defines a distribution *p*(**x**, *t* |***θ***), whose time evolution under Eq. (1) is given by [11]:

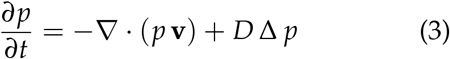

where Δ denotes the Laplacian on ℝ^*d*^. Standard finite difference or finite element methods for solving this equation are hopeless in high dimensions due to the exponential explosion of mesh elements needed for reasonable numerical accuracy.

We propose to circumvent the exponential increase in computational resources with dimension by defining a Markov process restricted to representative points sampled from the data. The time evolution of *p*(**x**, *t*) given by Eq. (3) is equivalent to the following recursion relation for *ϵ <<* 1:

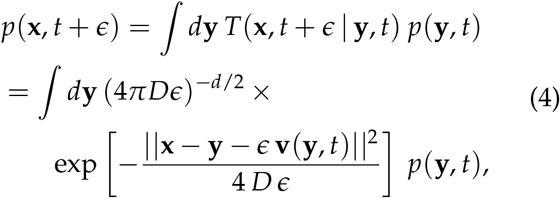

where *T*(**x**, *t*| **y**, *s*) denotes the transition probability from state **y** at time *s* ≤ *t* to state **x** at time *t*. In what follows, we adopt the simplified notation *T*(**x**, *t* | **y**, *s*) ≡ *T*_*ϵ*_(**x, y**), suppressing the dependence on time which enters only through **v** and ***θ*** for fixed *ϵ* = *t* − *s*. More details are provided in the Supplementary Information (see SI Appendix S1)

The discrete representation of our underlying dynamical manifold is a set of *N* points 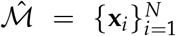 sampled from the experimental data. These points will typically be sampled from all time points and all experimental conditions in order to achieve robust coverage of all relevant regions of the gene expression space (SI Appendix S3.2). The local ‘base’ density *p*_*B*_ of the points in 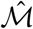 can be approximated using kernel density estimation (KDE), i.e. smoothing a sum of *δ*-functions centered on the **x**_*i*_ by convolution with a Gaussian kernel with a specified bandwidth (SI Appendix S1.2). We must also define a *d*dimensional volume *V*(**x**_*i*_) associated with each **x**_*i*_.

Several choices are possible for *V*(**x**_*i*_); the default construction in our code is *V*(**x**_*i*_) ~ 1/(*N p*_*B*_(**x**_*i*_)). The probability of finding a cell in the neighborhood 𝒱_*i*_ of **x**_*i*_ at time *t* is just

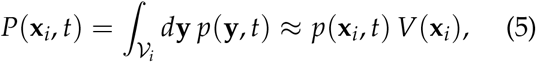

where 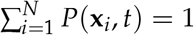. We then discretize Eq. (4) in the following way:

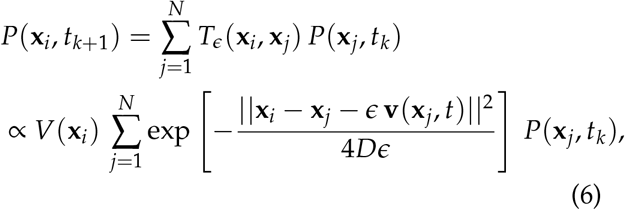

where the transition matrix is normalized so 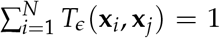 and *t*_*k*_ = *k ϵ* is the time at the *k*th step in the discrete Markov process. Precise specification of the discretization of **v** = − **g**^*−*1^ ∇*U*, alternative volume elements, and the normalization of *T*_*ϵ*_(**x**_*i*_, **x**_*j*_) can be found in SI Appendix S1.6. Suffice to say, given **v**(**x**_*i*_), we can construct a transition matrix *T*_*ϵ*_(**x**_*i*_, **x**_*j*_) that tells us how much probability transfers from **x**_*j*_ to **x**_*i*_ during the interval [*t, t* + *ϵ*]. We can therefore obtain an approximate solution to Eq. (3) at the 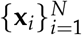 by iteratively applying *T*_*ϵ*_(**x**_*i*_, **x**_*j*_) to some initial condition *P*(**x**_*j*_, 0). Equation (6) discretizes a continuous operator, so for smooth solutions it should converge in the continuum limit. However, it remains to see how many points are needed when they are sampled from a distribution that is unrelated to the potential and diffusion constant being simulated, as will be the case with experimental data.

Fig. 2 explores the application of this method to the so-called ‘heteroclinic flip’ potential (Fig. 2A):

**Figure 2.**
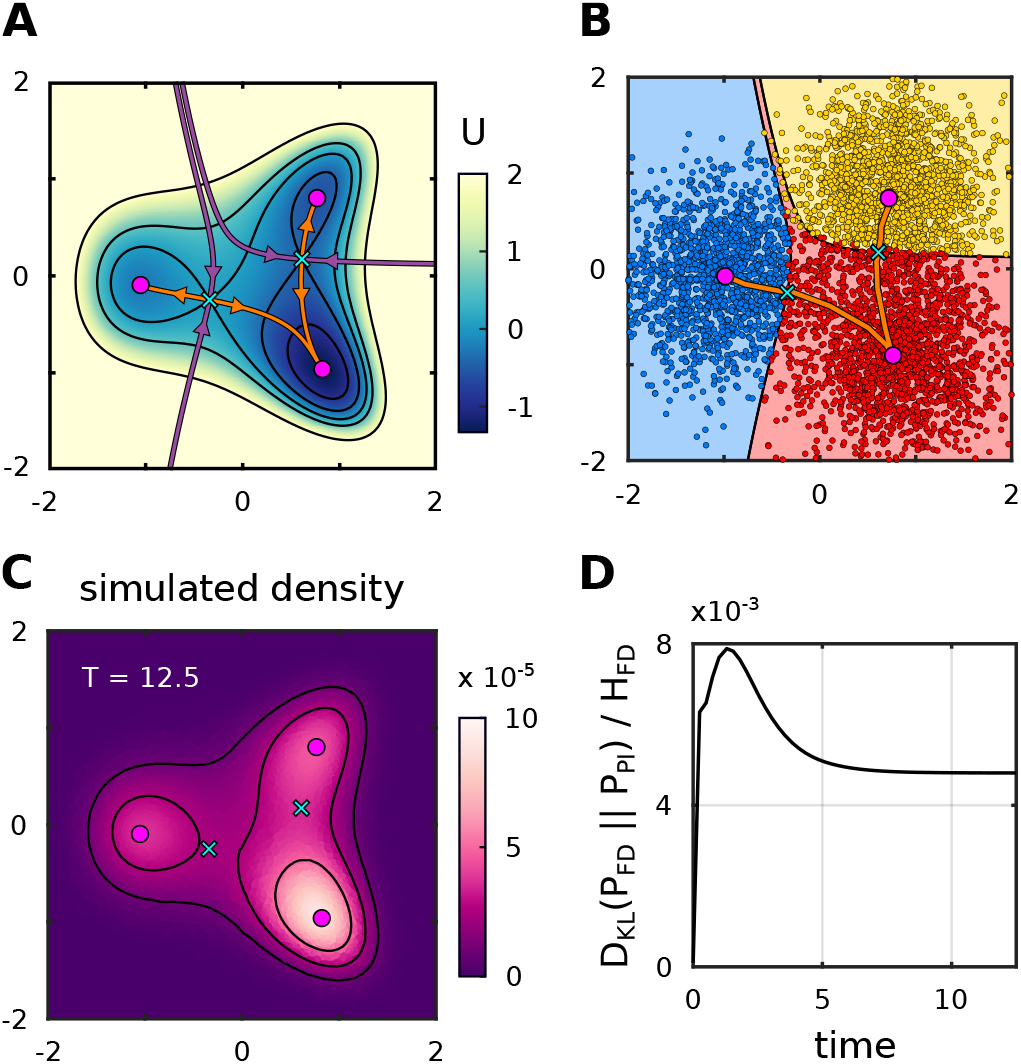
Discrete path integral formulation solves Fokker-Planck equation on non-uniform point sets. (A) The heteroclinic flip potential (Eq. (7)) for *a* = − 0.75 and *b* = 0.4. The magenta points are stable attractors, the cyan x’s are saddle points, the orange curves are unstable manifolds, and the purple curves are stable manifolds. (B) The discrete representation 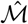 of the dynamic manifold is sampled from a three-component Gaussian mixture, with one isotropic component with covariance *σ* = 0.25 centered on each of the potential minima of shown in (A) (i.e., the distribution is not derived from the potential shown in (A)). The sinks in this panel were determined by finding the local maxima of the ground state density of the transition matrix *T*_*ϵ*_ constructed from the potential in (A) evaluated at the points in (B) for *D <<* 1. The unstable manifolds are the most probable paths between the inferred sinks. The saddles are the points along the unstable manifolds with the highest potential. Colored regions correspond to basins of attraction defined on the point set by solving the backward Kolmogorov equation using *T*_*ϵ*_ with the stable fixed points as a target set. Results almost perfectly match the segmentation inferred from the stable manifolds of the saddle points in (A), with only a few points being placed into the wrong basin. (C) The terminal density at *t* = 12.5 computed using Eq. (6) and an initial distribution centered around the ‘progenitor state’ near (−1, 0) with *D* = 1. (D) Kullback-Leibler divergences between the finite difference probabilities *P*_*FD*_ and the path integral probabilities *P*_*PI*_ normalized by the entropies, *H*_*FD*_, of the finite difference solutions of the Fokker-Planck equation. Finite difference solutions are transferred onto the non-uniform point set by area-weighted averaging for direct comparison.

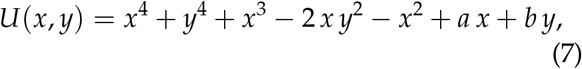

This is a crucial landscape that describes the common situation in development in which a single progenitor cell type (near (− 1, 0)) differentiates into one of two possible daughter cell types (near (1, *±*1)). The parameter *a* controls the stability of the progenitor while the parameter *b* tilts the landscape vertically and controls which daughter state receives the output of the progenitor state. Flipping the sign of *b* results in a ‘global’ bifurcation, since it entails the reconnection of the unstable manifold of the saddle nearest to the progenitor from one daughter state to the other.

The point set 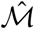 for this example consists of 5000 points sampled from a three-component Gaussian mixture, with one isotropic component centered on each of the minima of Eq. (7), evaluated with *a* = − 0.75, *b* = 0.4 (Fig. 2B). We then use the transition matrix in Eq. (6) to solve Eq. (3) with the trivial metric **g** = **I** and *D* = 1 directly on 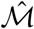 (Fig. 2C) We compare the results to a standard finite difference solution on a square grid over the same domain (SI Fig. S1) starting from the same initial condition. Despite the extreme heterogeneity of this point set (Fig. 2B), we obtain a highly faithful result relative to the finite difference solution (Fig. 2C and D).

Our construction enables us to capture multiple features of the system’s dynamical structure beyond simply solving Eq. (3). The stable fixed points of **v** can be found by examining the maxima of the stationary distribution of *T*_*ϵ*_. This distribution exists and is unique as long as *T*_*ϵ*_ is irreducible and aperiodic [18], both of which can be tested during the construction of *T*_*ϵ*_. In the limit *D* → 0, these maxima are sharply peaked and can be robustly identified using topological data analysis techniques [19] (SI Appendix S3.3). Briefly, these methods identify significant density maxima by constructing a proximity graph that connects nearby points, building a tree whose branches correspond to connected regions associated with particular local maxima, and simplifying the tree topology based on user-defined criteria like branch persistence, stability, or size. The unstable manifolds of the system are just the most probable paths between these stable fixed points, which can be found using standard methods for graph shortest paths with –log [*T*_*ϵ*_(**x**_*i*_, **x**_*j*_)] as directed edge weights (SI Appendix S3.4). The saddles are the points along the unstable manifolds with the highest potential. Fig. 2B shows that the fixed points and unstable manifolds determined using our discrete operator closely match those of the analytic system.

Finally, the basins of attraction of each stable fixed point can be computed using *T*_*ϵ*_ to solve the associated *backwards Kolmogorov equation* (BKE) (SI Appendices S1.1 and S3.6). The BKE tells us the probability that a stochastic trajectory starting at **x**_*i*_ at time *s* ≤ *t* collides with a given target set at time *t*. We can assign points in 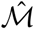 to the fixed points of **v** by solving the BKE at small *D* for a short time and then matching manifold points to the fixed point they are most likely to hit. This construction exactly produces the basins of attraction in the deterministic limit. Fig. 2D demonstrates that these methods accurately reproduce results obtained using conventional methods.

## Algorithm Applied To Simulated Data

We now demonstrate how our construction can be fit to a typical experimental data set. As a test case, we generate synthetic data from Eq. (7), using three different parameter sets (*a, b*) (denoted ***θ*** above) to represent distinct experimental conditions and sample trajectories at multiple time points (Fig. 3). To determine the dynamical structure of the data, we must build the point set manifold 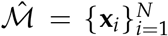 and then fit a potential that describes the dynamics of the transition from progenitor to terminal states. 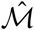 is built from data points, pooled together from all time points and experimental conditions, which we then subsample down to a tractable number (here *N* = 5000) (Fig. 3A). We then infer the point-wise density *p*_*B*_(**x**_*i*_) on 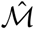 using KDE and extract prominent maxima via topological extrema detection methods, as before. We assume that the local maxima in *p*_*B*_ correspond to the set of all intermediate and terminal cell states sampled by our spectrum of conditions (Fig. 3B). KDE is also used to transform the discrete clouds of data points at each time into time-dependent probability distributions defined on 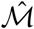 (Fig. 3C and SI Figs. S3 to S5).

**Figure 3.**
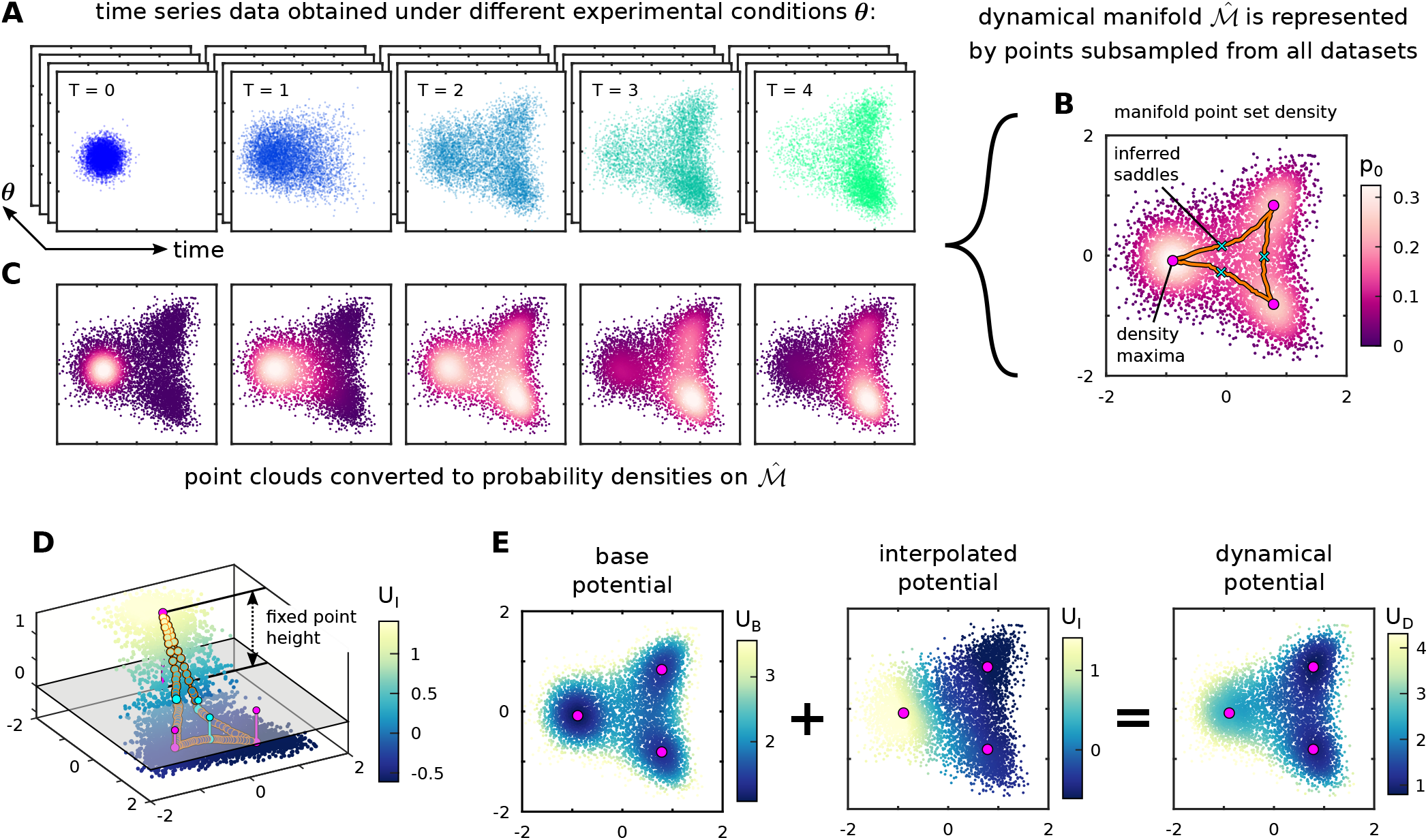
Fitting the dynamical structure of a synthetic data set. (A) A time course of points generated by simulating the gradient-like dynamics of the heteroclinic flip potential (Eq. 7) with noise *D* = 1 and ***g*** = ***I*** for three sets of parameter values *θ* = (*a, b*). Scatter plots show the dataset for *a* = *−*1.4 and *b* = 0.5 at the five sampled time points, arbitrarily labeled *T* = 0 to 4 and color coded. (B) The discrete point set 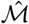 sampled from all conditions, *θ*, and times. The unstable manifolds are the most probable paths between density maxima and the saddles are the points along those paths with minimum density. (C) The actual data in (A) from T=0-4 are smoothed onto the points in 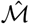 via KDE to define a density *p*(**x**) to compare with solutions of Eq. (6). (D) A schematic of our interpolation procedure used to construct a smooth potential *U*_*I*_. For a given set of fixed point heights 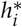, defined on the sinks and saddles, *U*_*I*_ linearly interpolates between the 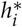 along the unstable manifolds and smoothly interpolates everywhere else by minimizing the energy Eq. (8), which penalizes curvature. The fit parameters can be iteratively optimized until we have a dynamical landscape that accurately captures the dynamics observed in the data. (E) Construction of our dynamical potential *U*_*D*_ = *U*_*B*_ + *U*_*I*_. By adding the *U*_*I*_ to a base potential *U*_*B*_ inferred from *p*_*B*_, we can build a dynamical potential *U*_*D*_, for each data set *θ*, that captures the directed, non-steady state motion in the high dimensional feature space (see Table 1).

We must now fit a condition-dependent potential to describe the non-steady state dynamics. The first ingredient is the ‘base potential’, *U*_*B*_(**x**_*i*_) := −log *p*_*B*_(**x**_*i*_) which characterizes how probability diffuses from point to point in the absence of directed motion and confines dynamics to the manifold’s interior. In order to model the target dynamics, we clearly need a function that will destabilize the base potential well near the initial condition and deepen the wells with a positive *x*-coordinate, i.e., the terminal states. Let 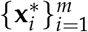 refer to the set of *m* density maxima. Also, let 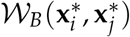 denote the most probable path between 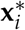 and 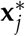 defined relative to the base

**Table 1.**
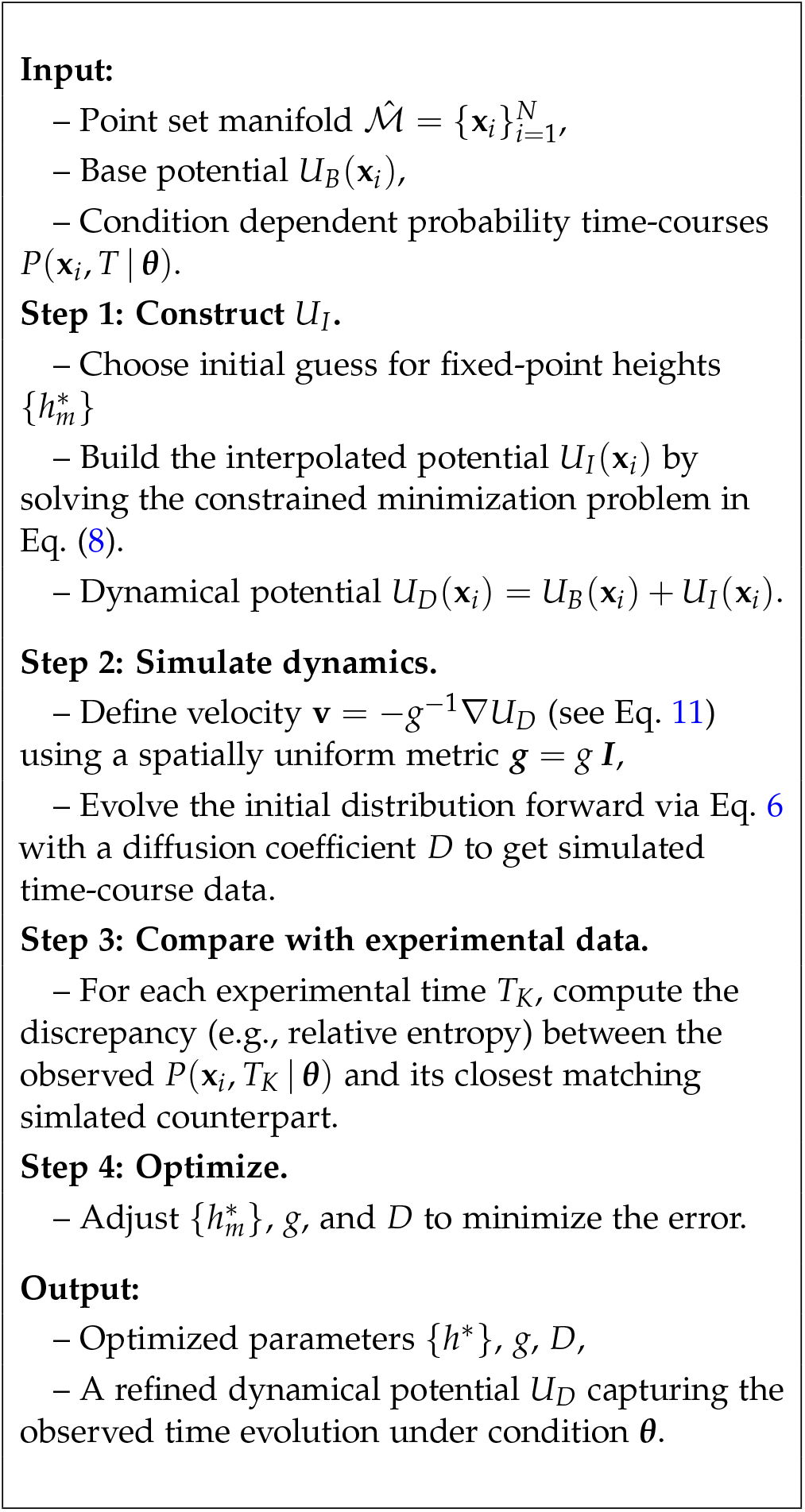
Pipeline for constructing and fitting the dynamical potential *U*_*D*_ from data acquired under experimental conditions *θ*.

potential *U*_*B*_ (i.e.,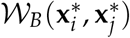 is an ordered list of points 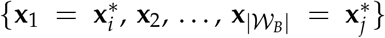 where |𝒲_*B*_ | is the number of points in the path). The function we seek can be found as the minimizer of a smoothness functional, namely the squared Laplacian energy (see, for instance, Section 3.8.1 of [20]):

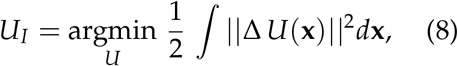

subject to:

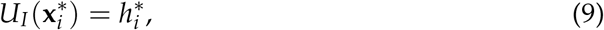

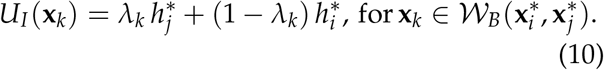

The *λ*_*k*_ *∈* [0, 1] are just the fractional position of **x**_*k*_ along 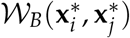. In other words, we adjust the “heights” of the fixed points 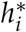, constrain the potential along the most probable paths to linearly interpolate between them, and smoothly interpolate everywhere else by minimizing Eq. (8). This framework is illustrated for a particular set of fixed point heights in Fig. 3D and E. More details about the computation of *U*_*I*_ can be found in SI Appendix S1.5. Our final dynamical potential is given by

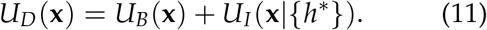

Note that all confinement comes from *U*_*B*_ as *U*_*I*_ is flat far away from the manifold ℳ. The choice of *U*_*I*_ eliminates local minima and saddle points that were inferred from *U*_*B*_, but are not relevant to the developmental dynamics under the experimental conditions being modeled. See SI Fig. S6 for an illustration of this construction.

Our numerical machinery for solving Eq. (3) in its current form can handle arbitrary conformal metrics (i.e. **g** = *e*^Ω(**x**)^**I** [21]). However, we assume a uniform scalar metric **g** = *g* **I** in all subsequent analyses, since this choice is sufficient to accurately fit all experimental data considered here. The complete set of fitting parameters are the 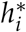 in Eq. (9), the diffusion constant *D*, the scalar metric *g* and optionally the heights of the saddle points, (whose locations are again inferred from the base potential *U*_*B*_). Note that *g* rescales *U*_*D*_ = *U*_*B*_ + *U*_*I*_ and therefore cannot simply be absorbed into the heights 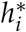. From Eq. (6), we see that *D* rescales all terms in the exponent of the transfer operator, while *g* specifically rescales **v** = –(∇*U*_*D*_)/*g* (see also SI Appendix S1.6). Fitting these two scalar values separately therefore sets the relative importance of diffusion versus drift dynamics. The complete pipeline, applied to data acquired under experimental conditions ***θ***, is schematized in Table 1.

The results of this procedure for our synthetic dataset are shown in Fig. 4 and and SI Figs. S3 to S5. The “true” potentials (Fig. 4A) are well approximated by the effective potentials *U*_*eff*_ = *–g D* log [*p*_*eq*_(***θ***)], where *p*_*eq*_(***θ***) is the equilibrium density of the transition matrix *T*_*ϵ*_ (Fig. 4B). We use *U*_*eff*_ since the ground state of our *T*_*ϵ*_ is not exactly exp [*–U*_*D*_/(*g D*)] due to discretization error and *U*_*eff*_ tends to capture dynamical features better than *U*_*D*_ (see SI Fig. S6). It is sensible that *U*_*eff*_, which is derived from an equilibrium probability density, faithfully reflects the structure of the underlying drift dynamics since our fitting procedure minimizes the error between measured and simulated probability time courses. The optimal Kullback-Leibler *D*_*KL*_ ~1e-2 bits between the simulated and observed probability time courses are a minuscule fraction of the observed entropy of the data, *H*_meas_ ~12 bits, as anticipated in Fig. 2D (SI Table S1). The inferred fixed points, unstable manifolds, and basins of attraction (Fig. 4C) closely match those of the true potential (SI Table S2).

**Figure 4.**
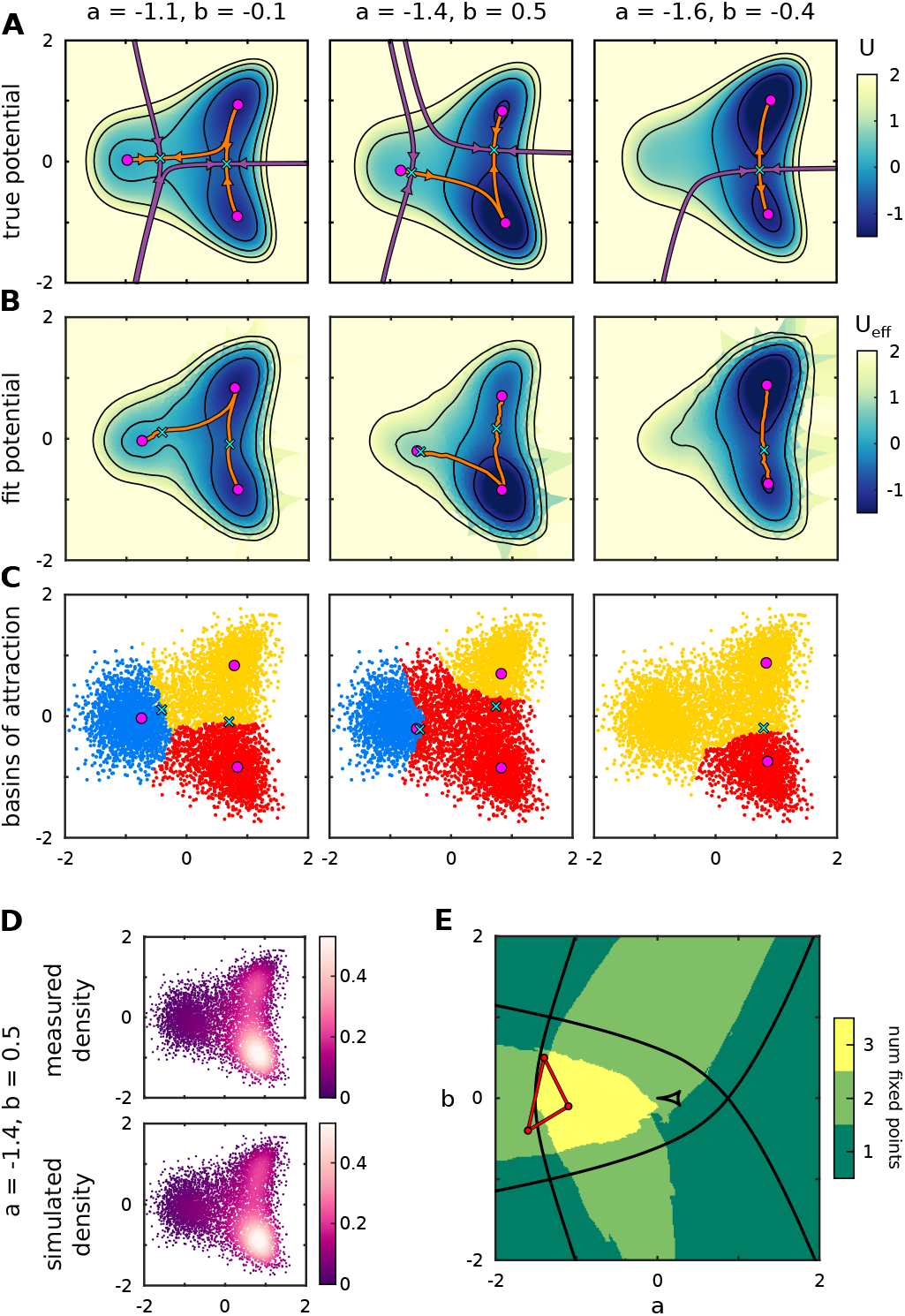
Fitting procedure accurately captures the dynamical structure of a synthetic data set. (A) The true potential used to generate the synthetic data time courses in Fig. 3A. (B) The effective potentials, *U*_*eff*_ = *−g D* log [*p*_*eq*_ (*a, b*)], fit to the synthetic data. where *p*_*eq*_ is the equilibrium density of the transition matrix *T*_*ϵ*_ constructed for a given set of (*a, b*). The sinks, saddles, and unstable manifolds are inferred directly from the data, as described in the text. (C) The basins of attraction for the fit potentials computed from the backward Kolmogorov equation closely match those in A. (D) Simulated and measured densities at the final time point for for *a* = −1.4 and *b* = 0.5. (E) A bifurcation diagram for the inferred dynamics. For each set of parameter values on a 256 × 256 grid in (*a, b*)-space a new potential is generated by linear barycentric interpolation from the three potentials fit directly to experimental data (inset triangle). The number of stable fixed points is then computed using our topological extrema detection methods. Our method produces a similar bifurcation structure to that of the true underlying dynamics despite extrapolating from only three parameter values. The small set of fold bifurcation curves near (*a, b*) = (0, 0) corresponds to a shallow repeller, which is not captured during our search for stable fixed points.

Finally, our method enables us to make out-of-sample predictions about the dynamical structure of our system, as demonstrated by the bifurcation diagram in Fig. 4D. For each set of parameter values on a 256 × 256 grid in (*a, b*)-space, we produce a new potential by linear barycentric interpolation between the three fit values of *U*_*eff*_ and determine the number of stable fixed points using our topological data analysis methods. Our method produces a similar bifurcation structure to that of the true potential over a broad range of experimental parameters despite extrapolating from only three pairs of (*a, b*)-values.

## Landscape model for neural tube patterning

### Static morphogens

We now apply our method to elucidate the dynamical structure of cell fate specification in an *in vitro* model of neural tube development. The neural tube is an embryonic structure in vertebrates that gives rise to the central nervous system, including the spinal cord. It is patterned by opposing gradients of BMP/Wnt from the dorsal side and Sonic Hedgehog (Shh) from the ventral side. The two morphogen gradients define 11 domains of neural progenitors, arranged linearly along the dorsoventral (DV) axis, each of which is uniquely identifiable by a characteristic pattern of transcription factor (TF) expression [12]. Prior work focusing on the ventral side, including *in vivo* data [14], modeled the system as a gene regulatory network (GRN) with a dynamic Shh gradient and mutual repression between the TFs that define the progenitor domains (see Fig. 5A).

**Figure 5.**
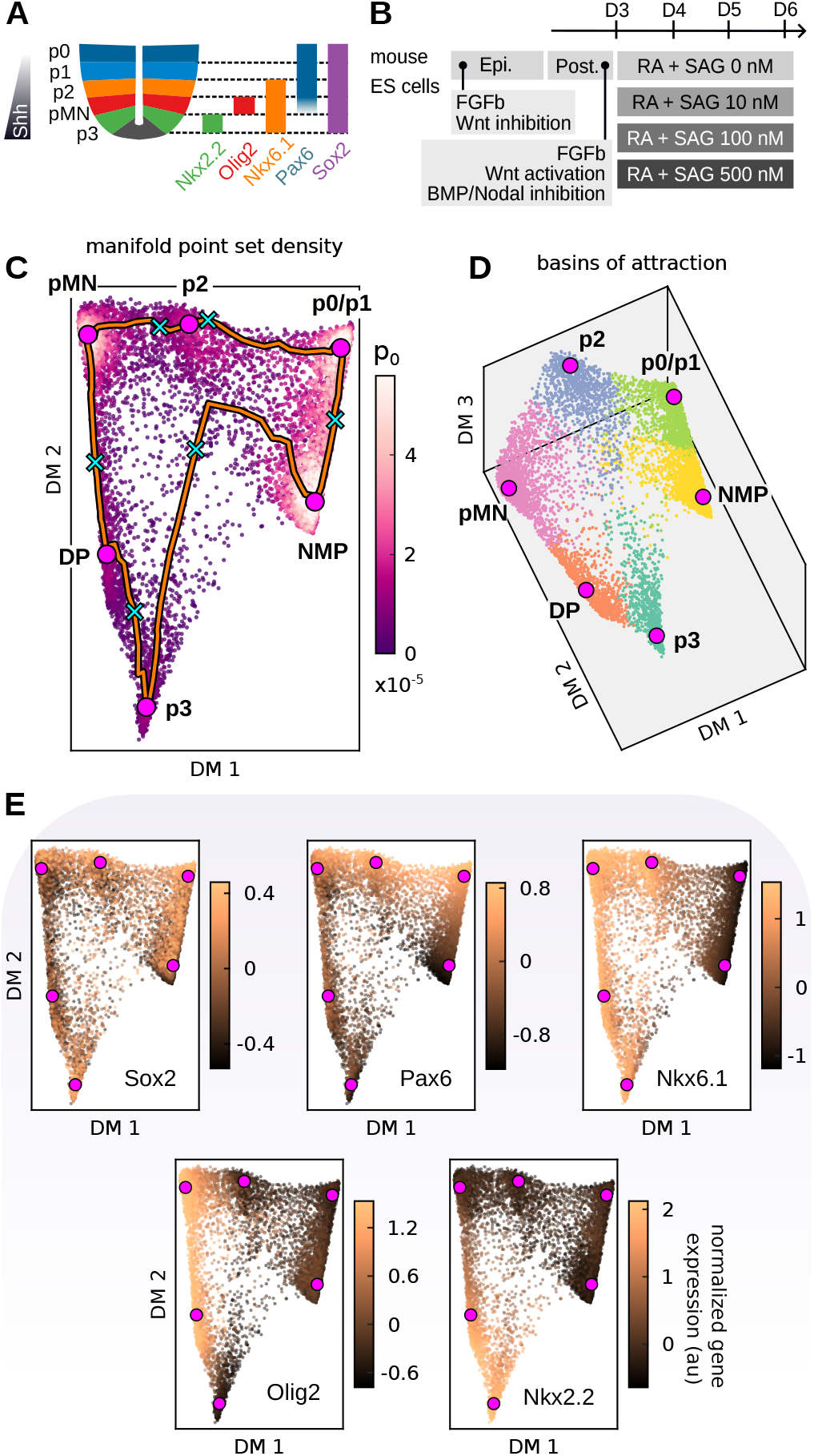
Dynamical manifold architecture of *in vitro* proneural patterning. (A) Schematic of ventral neural tube progenitor domains and the corresponding combinatorial transcription factor code used for multi-color flow cytometry. (B)Schematic of the method used to differentiate mouse embryonic stem cells into ventral neural progenitors [13].(C)Visualization of the density on the discrete dynamical manifold subsampled from all experimental conditions (SAG levels and times) projected onto the top two diffusion map dimensions. The density maxima correspond neatly to the neural progenitor types using (E). (D) The basins of attraction for the local density maximum, projected onto the three lowest diffusion map dimensions, showing the dynamical manifold is approximately two dimensional (SI Movie 1). (E) Normalized marker expression viewed on the dynamic manifold identifies the density maxima with neural progenitors in (A).

*In vitro* differentiation of stem cells offers direct control over morphogen levels and timing, thereby reducing the complexity of endogenous cell–cell interactions. This setup provides an ideal testbed for whether our landscape formalism can capture the essential patterning dynamics under externally controlled conditions. We apply our framework to an *in vitro* dataset, originally presented in [13], where mouse embryonic stem cells (mESCs) were driven toward a neuromesodermal progenitor (NMP) state (Fig. 5B). Starting on Day 3, cultures received varying concentrations of the Shh agonist SAG to mimic DV patterning *in vivo*. Fates were measured at 24-hour intervals by multidimensional flow cytometry analysis, that exploited combinatorial TF expression to define distinct progenitor domains (Fig. 5A).

Our pipeline, following Table 1, begins with the construction of a discrete manifold 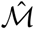, built by subsampling 8000 cells from multiple measurement times and SAG concentrations. For visualization, but not analysis, we project all data from the 5D flow cytometry data space down to two or three dimensions using diffusion maps [22] (Appendix S1.3). Our topological extrema detection methods extract six density maxima from *p*_*B*_ (Fig. 5C and D). Comparison of the point-wise normalized gene expression profiles across 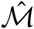 (Fig. 5E) to the known TF code in Fig. 5A shows that five of the six maxima correspond neatly to NMP and the various neural progenitor states. Since the available markers do not distinguish p0 from p1, we group them together in all subsequent analyses. The final density maximum corresponds to a persistent ‘double positive’ (DP) state, defined by simultaneous expression of Nkx6.1, Olig2, and Nkx2.2 [13].

The measured probability distributions de-fined on 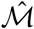 were fit following the procedure in Table 1. We determined the mapping from simulation times to experimental times by pairing each measured distributions with its best-matching simulated counterpart, then fitting these pairs using a power law whose parameters minimize the average discrepancy across all measured times (see SI Appendix S3.8 and SI Fig. S17). Note that not all progenitor types will be stable at a given morphogen level. To partition point-wise probability distributions, even in situations where a progenitor type is unstable, we assign points to the six density maxima using our backward Kolmogorov method relative to the base potential *U*_*B*_. This method produces clean basins of attraction that allow analysis of the population of cell-types (Fig. 5D) as a function of time.

The fit to the SAG 0 nM dynamics was performed as a stand alone procedure. Recognizing that landscapes must depend on morphogen levels, we imposed a simple functional constraint to simultaneously fit higher SAG levels. SAG 10 nM and SAG 500 nM were each endowed with a complete set of optimizable fit parameters, but the dynamics at the intermediate SAG 100 nM was found by linearly interpolating between the dynamical landscapes:

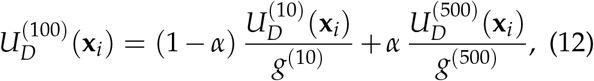

where *α* is a single interpolation parameter for all points in the discrete manifold and *g*^(100)^ = 1. The scalar diffusion coefficient was interpolated similarly as *D*^(100)^ = (1 *– α*) *D*^(10)^ + *α D*^(500)^.

Fig. 6A-D show the *U*_*eff*_ fit to each constant SAG level. Here, the highlighted points are not density maxima, but the true stable attractors for each landscape, once again determined using topological extrema detection methods. The SAG 0 nM landscape has only one attractor corresponding to p0/p1. As the SAG concentration increases, the attractors are sequentially stabilized and destabilized until, at SAG 500 nM, only the most ventral states are stable. For all SAG levels, we find that the eigenvalues of the Hessian of *U*_*eff*_ at the sinks are uniformly negative (a dynamical requirement for totally stable attractors), and all saddle points connecting them have only one unstable direction (see SI Appendix S3.10 and SI Table S4).

**Figure 6.**
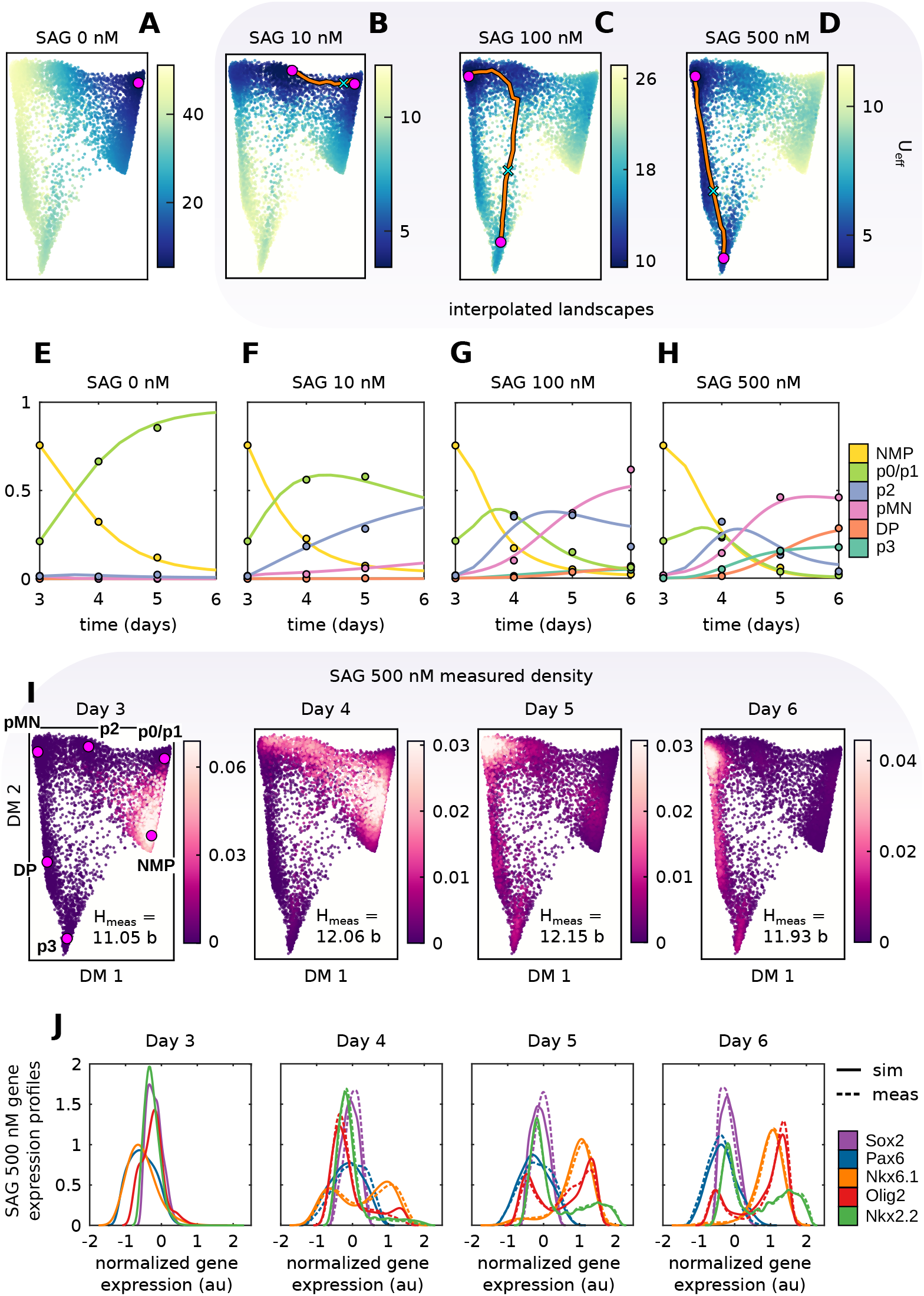
Fitting procedure accurately captures the dynamical structure of neural progenitor differentiation for constant SAG. (A)-(D) Dynamical potential landscapes that reproduce days 4-6 at the indicated SAG levels. The SAG 100 nM landscape was linearly interpolated from the landscapes obtained for SAG 10 nM and SAG 500 nM as explained in the text (*α* = 0.637). Note only certain states from Fig. 5C survive at various SAG levels. (E)-(H) Point set manifold basin probabilities over time. Filled circles show measured values and solid lines show the simulated basin probabilities. The day 6 data for SAG ¡ 500 is contaminated by ‘aging’ as explained in the text. (I) Measured time courses for SAG 500 nM. The fits are indistinguishable to the eye from the data with a K-L divergence ≲ 0.3 bits at all times compared to a measured entropy of ~12 bits for the raw data. (J)-(M) Time courses of normalized antibody expression distributions for SAG 500 nM. Dotted lines are measured distributions and solid lines are simulated distributions.

The time-dependent fits in Fig. 6E-H closely match the data (SI Figs. S10 and S11 and SI Table S3). For SAG 0 nM, probability rapidly leaves NMP and populates the p0/p1 basin. Increasing the concentration to 10 nM SAG results in more probability density reaching the p2 basin by Day

5. Further increasing the concentration to 100 nM SAG leads to a mixture composed primarily of pMN and p2. The measured dynamics at 500 nM SAG are shown in detail in Fig. 6I. By Day 4, most probability has left the NMP state and flows towards p2, largely avoiding the destabilized p0/p1 basin Fig. 6D. A smaller, but non-negligible, amount of probability appears to flow directly from NMP towards p3, a route that was not accessible at lower SAG levels (Roughly ~6% of the total probability exists in the p3 basin on Day 4. 92% of that fraction of the probability never passes through the pMN or DP basins – see SI Appendix S3.11). Probability continues to flow toward the ventral states and, by Day 6, almost all probability is concentrated in pMN, DP, and p3 The *D*_*KL*_ between the simulated and measured distributions is ≲ 0.3 bits for all conditions (see SI), a small fraction of the observed entropies (*H*_meas_ ≳ 11 bits). Remarkably, the linear interpolation method in Eq. (12) produces a fit to the SAG 100 nM dynamics with only ~1% error above fitting SAG 100 nM as a stand alone procedure (SI Table S3). We also reproduce with comparable accuracy the actual fluorescence levels as a function of time and SAG level Fig. 6J (see also SI Figs. S12 and S13).

We note that ‘aging’ effects in the system prevented a detailed analysis of the measured dynamics at low SAG concentrations on Day 6. These effects are a result of using a restricted probe set. Sox2 and Pax6 have broad expression in the developing neural tube. They transiently increase and then decrease for all SAG levels (see Fig. 6J and SI Figs. S12 and S13), which matches observations *in vivo* [23]. Since these are the only markers distinguishing p0/p1 from NMP, it appears that probability flows back uphill to NMP after Day 5 for low SAG concentrations (see SI Fig. S10). We therefore chose to ignore Day 6 in our fits for 0 nM and 10 nM SAG. The same phenomenon affects the Day 6 data for SAG 100 nM and 500 nM (Fig. 6G and SI Figs. S12 and S13), but did not prevent a high quality fit.

We tested the reproducibility of the fit parameters by running all optimizations with different random initial guesses for the fixed point heights (the initial guesses for both *D* and *g* were always set to 1) (SI Figs. S15 and S16). While the individual values of *D* and *g* were variable, the product *g D* and the heights at the fixed points were surprisingly consistent. For instance, when optimizing to find the parameters that defined the interpolated landscapes the mean and standard deviation of *D g* were *g*_10_ *D*_10_ = 1.01 *±* 0.09 and *g*_500_ *D*_500_ = 1.10 ± 0.17. Moreover, the standard deviations of the rescaled fixed point heights (both sinks and saddles) *h*_*i*_ / (*g D*) divided by their means were only ~5 –15%. This consistency was observed for both the interpolated fits and standalone fits. The only condition that exhibited notable variability was SAG 0, which is reasonable considering that SAG 0 is characterized by a unimodal population of p0/p1. In other words, it does not matter precisely what the ventral heights are for SAG 0, just that almost all the probability eventually settles near p0/p1. Reproducibility for all optimizations is provided in the SI.

We also explored the effect of reducing the number of fit parameters. All displayed results were generated by optimizing over both the heights of the density maxima and the heights of the base potential saddles (Fig. 5C). Reproducing these fits while optimizing only over the heights of the density maxima only slightly reduced the quality. Results on average were ~15% worse than the original fits, but this translates to only 0.04 extra bits, which is still small compared to the measured entropies of ~12 bits, meaning that these more parsimonious fits still explained most of the information in the measured data. The standalone fits to SAG 0 without base potential saddles were of similar high quality (SI Table S3). This suggests that fitting the heights of the saddles is not always strictly necessary.

### Time varying morphogens

We next asked whether chaining together landscapes fit to different constant SAG levels can capture the dynamical features of time-dependent perturbations. Two protocols were implemented (Fig. 7G): SAG 0-500 and SAG 500-0, each with 24 h exposure to the first concentration followed by continued exposure to the second concentration until Day 6. We analyze these experiments in two distinct ways. The first we term “static fit”, in which we simply evolve forward the measured distribution on Day 4 using the landscape previously fit to the constant SAG levels (Fig.6A and D) The ‘static fits’ in Fig. 7C and F did not accurately reflect measured dynamics on Days 5 and 6 (see below). The ‘two-stage’ landscapes were fit exclusively to data from Days 5 and 6 after the media change with the measured Day 4 data as an initial condition. The two-stage fits, which by construction capture the topography of the landscape following the perturbation, closely match the measured data for both protocols (Fig. 7A, B, D and E). See SI Figs. S18 and S19 and SI Table S5 for more details.

**Figure 7.**
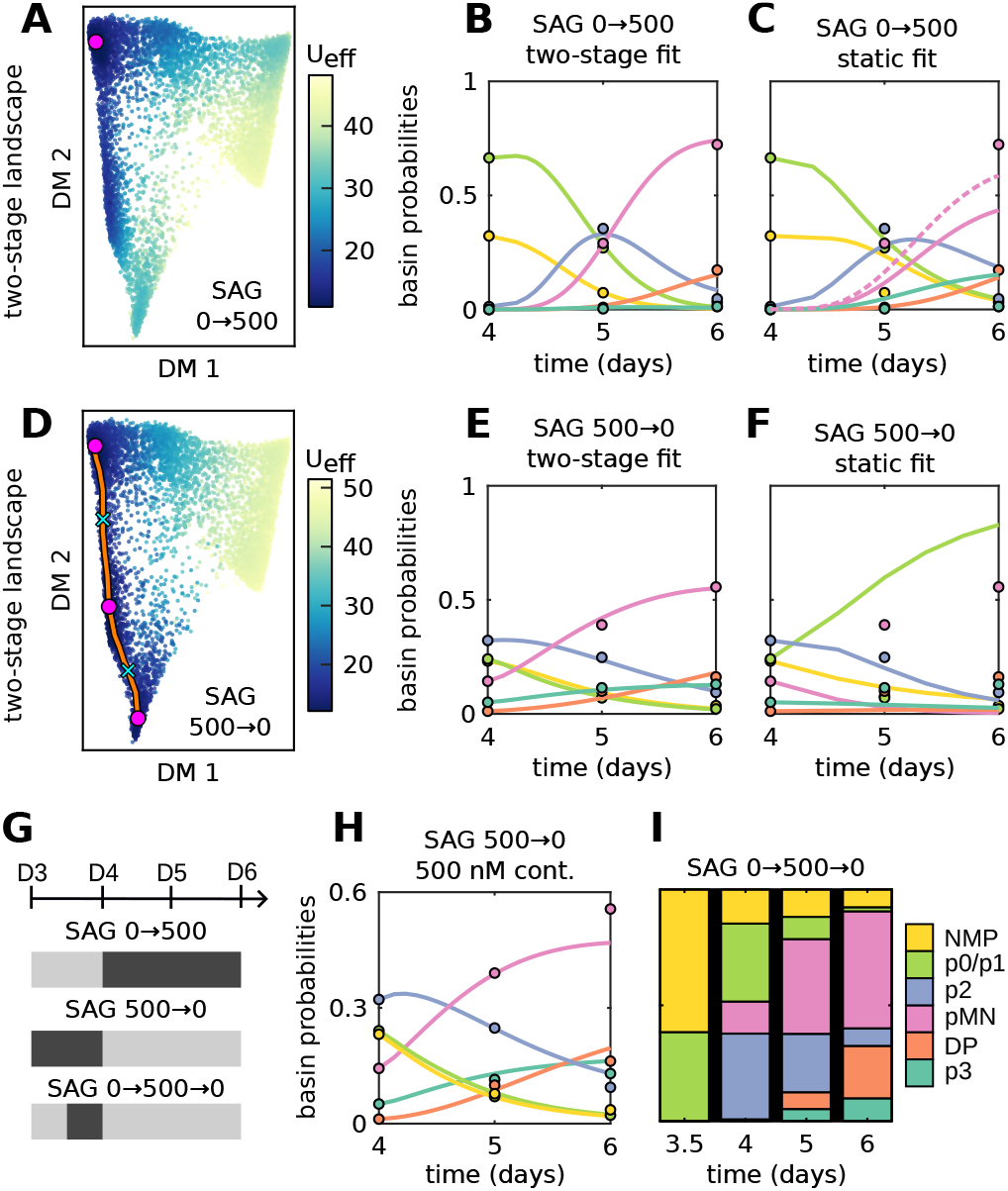
Cell type proportions for time-dependent protocols. (A-C) Fits to dynamic data for 24 h SAG 0 then SAG 500 through Day 6. The static fit, uses the models from Fig. 6. SAG 500 is initialized with the Day 4 prediction for SAG 0. The two stage fits are de-novo beginning from the measured data on Day 4 distribution. Dotted line in (C) shows the sum of the pMN and p3 basin probabilities. (D-F) The same plots for 24 h SAG 500 then SAG 0 through day 6. (G) The dynamic protocols, including 12 h pulse at SAG 500. (H) Fit of the 500 → 0 by stretching the time scale on the constant SAG 500 model initialized from the measured data on Day 4. (I) Basin proportions 12 h SAG 500 pulse data showing that the p0/p1 fraction continually decreases after SAG is reset to 0.

The failure of the static fits points to new aspects of the biology. In [13], it was shown that expression of the TF FoxA2 during Day 3 was necessary to open chromatin domains that are required to realize the p3 fate. Hence, there is some direct transfer from NMP to p3 that does not transition through more dorsal states in the constant SAG 500 nM landscape (Fig. 6D, H, and I and SI Appendix S3.11). SAG 0-500 eliminates this direct route to p3. The two stage potential, Fig. 7A, has a single minima at pMN and a higher value at p3, while for constant SAG 500 it is bistable at pMN and p3 (Fig. 6D). Despite the downregulation of the p3 fate, the relative proportions of aggregated ventral fates (p3, DP, and pMN) versus dorsal fates (p2 and p0/p1) on Day 6 are quite similar to the proportions observed at constant SAG 500 nM (compare Fig. 7B and C to Fig. 6H) Allowing the SAG 0 state to persist and computationally scanning all possible delays in the application of SAG the 500 potential beyond day 4 did not improve the fits. This corroborates that the low proportion of p3 is due to an irreversible change in the landscape, not just some change in time scale.

The discrepancy between the two-stage and static fits for SAG 500-0 nM is more marked (Fig. 7E and F). Recall that constant SAG 0 nM produces a unimodal population of p0/p1 cells (Fig. 6A and E). In stark contrast, the data for SAG 500-0 on Day 6, have almost no probability in the p0/p1 basin and instead are a mixture of the ventral states (primarily pMN). The two-stage fit landscape is tristable with attractors at pMN, DP, and p3 (Fig. 7D). Several hypotheses can be proposed to explain this behavior. It is possible that there is a single static landscape for SAG 0 nM that features a deep well at both p0/p1 and pMN separated by a high saddle around p2. In other words, the pMN basin exists at SAG 0nM but is inaccessible until the 24-hour pulse of SAG 500 nM shifts enough cells across the p2 saddle to the pMN basin. Fitting a constant landscape simultaneously to the SAG 0 nM constant data and the SAG 500-0 data from Days 4 to 6 indeed produces a landscape with attractors at both pMN and p0/p1. However, even when we exhaustively scan transient signaling delays or different values of *D* and *g*, we could never produce a distribution with negligible p0/p1 on Day 6 (SI Fig. S19). Moreover, using this common SAG 0 nM landscape somewhat degrades the fidelity of the fit to constant SAG 0 nM alone (SI Table S3).

The possibility of a single, static, multi-stable landscape is further contradicted by an additional experiment undertaken to elucidate this phenomena. The new experiment exposes cells to 12 h of SAG 0 nM to eliminate the NMP to p3 transition and force cells towards p0/p1, followed by 12 h of SAG 500 nM, before returning to SAG 0 nM until Day 6 (Fig. 7G). This experiment is specifically designed to test whether the SAG 0 nM landscape could simultaneously have two wells (p0/p1, pMN), only one of which is populated in the SAG 0 nM constant and SAG 500-0 protocols. Inspecting the measured probability time courses shows that a great deal of probability remains near the p0/p1 well on Day 4 coincident with the withdrawal of SAG 500 nM (Fig. 7I and SI Fig. S23). If an attracting state remained in the landscape some probability would linger near it. However, by Day 6 there is essentially no probability in the p0/p1 well. In light of this evidence, the most likely explanation is that the dynamical landscape explored by the cells is irreversibly changed by 12 h exposure to 500 nM SAG. Indeed, simply keeping the static 500 nM SAG landscape and refitting a time scale to data initialized from Day 4 is sufficient to reproduce the measured behavior of the SAG 500-0 protocol (Fig. 6H) and the SAG 0-500-0 protocol (SI Fig. S24) at the level of basin probabilities. In other words, the most likely explanation is that after reduction of SAG from 500 to 0, the p0/p1 attractor remained bifurcated (was not restored) and differentiation continued, albeit with a different time scale (compare Figs. S17, S22 and S24).

Finally, we used our methodology to explore the possibility of maximizing the terminal proportions of particular cell types by invoking Eq. (12) to tune the SAG level, and derive the predicted ground state basin proportions shown in Fig. 8A. With the exception of the unimodal distribution of p0/p1 at SAG 0 nM, none of the SAG concentrations have an equilibrium distribution *p*_*eq*_ characterized by a single population of progenitors. When cells are differentiated *in vitro* there is plausibly more variability in the immediate environment of each cell plus uncontrolled cell-to-cell signaling that appears as noise in our fits. We speculate that the situation *in vivo* can be modeled by reducing the diffusion constant (Fig. 8B). Similarly, Ref. [24] argues that the boundaries between differently fated regions are sharp because of dynamic properties of the GRN and implicitly invoke the relative heights of saddles and fixed points versus molecular noise. We see that the p0/p1, p2, and pMN domains effectively unimodally tile the SAG axis from 0 nM SAG up to ~250 nM SAG when we halve the diffusion constant. We never see a unimodal population of p3, and SAG 500 is saturating, so we assume that the situation *in vivo* reflects a signaling history that was not replicated in vitro, or additional signaling cues [13].

**Figure 8.**
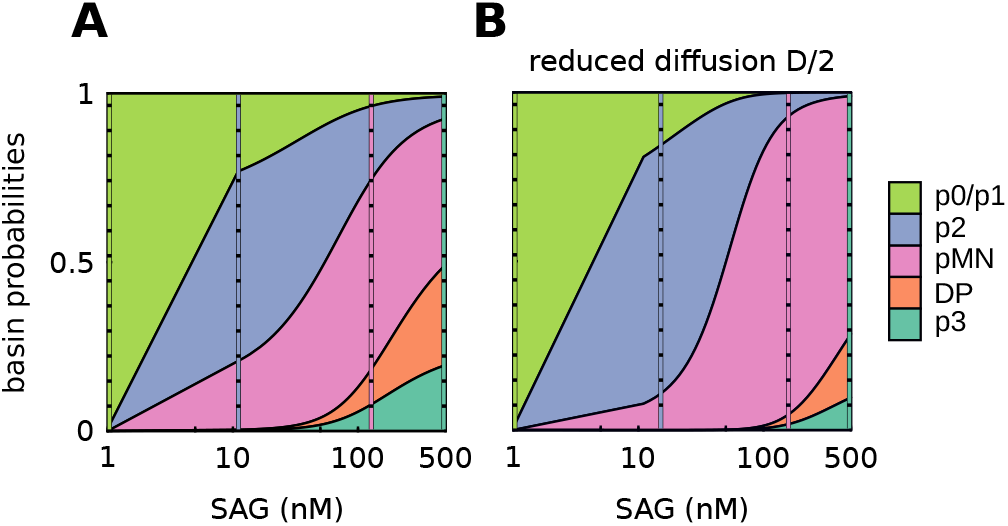
(A) Basin proportions for the equilibrium distributions as a function of SAG. Values of SAG in the range 10 nM to 500 nM are estimated by fitting a simple functional form the linear interpolation on potentials in Eq.12 (see SI Appendix S3.9). SAG 0 nM is fit separately. The x-axis is 1+SAG, on a log scale. (B) Same quantities as in (A), recalculated with a diffusion constant reduced by half to approximate the lower noise typical in *in vivo* conditions. The maximum proportions of p2 and pMN shown by the dashed vertical lines are both 56% in (A), 74% and 88%, respectively, in (B) and are 92% and 97% for D/4 (not shown).

### RNA-seq

Above, we analyzed flow cytometry data, which accurately measures a limited number of proteins. By contrast, single-cell RNA-seq can profile thousands of genes, but often at lower accuracy and with batch effects. In *in vitro* differentiation studies, the challenge remains to reduce the dimenxsionality of these data to approximate the hypothesized low-dimensional landscapes [3, 10]. We therefore turned to RNA-seq data from [25], which follows the protocol in Fig. 5B but applies SAG 500 for days 3-8.

Our goal here was to assess the efficacy of our method in analyzing the noisy, high-dimensional data typical of RNA-seq. For the sake of simplicity, we focused on a subset of 7293 cells sequenced between Day 5 to Day 8, identified by Fontaine et al. [26], that captured the differentiation of the most ventral fates (i.e. floorplate and p3). Fontaine et al. clustered these cells into putative types and identified 94 genes involved in the differentiation process, then refined this to 17 genes most correlated with ventral states. We used SANITY [27] on the 94 genes to normalize the data and then restricted analysis to the 17 gene subset. We subsampled 4000 cells from this normalized data, with equal numbers of points on each day, and characterized the structure of the resulting discrete manifold. Our topological extrema detection methods extracted three density maxima (Fig. 9B and D). Looking at the mean gene expression in the 30 nearest cells to the density maxima reveal that the 17 marker genes neatly delineate these populations into three distinct cell classes (Fig. 9A).

**Figure 9.**
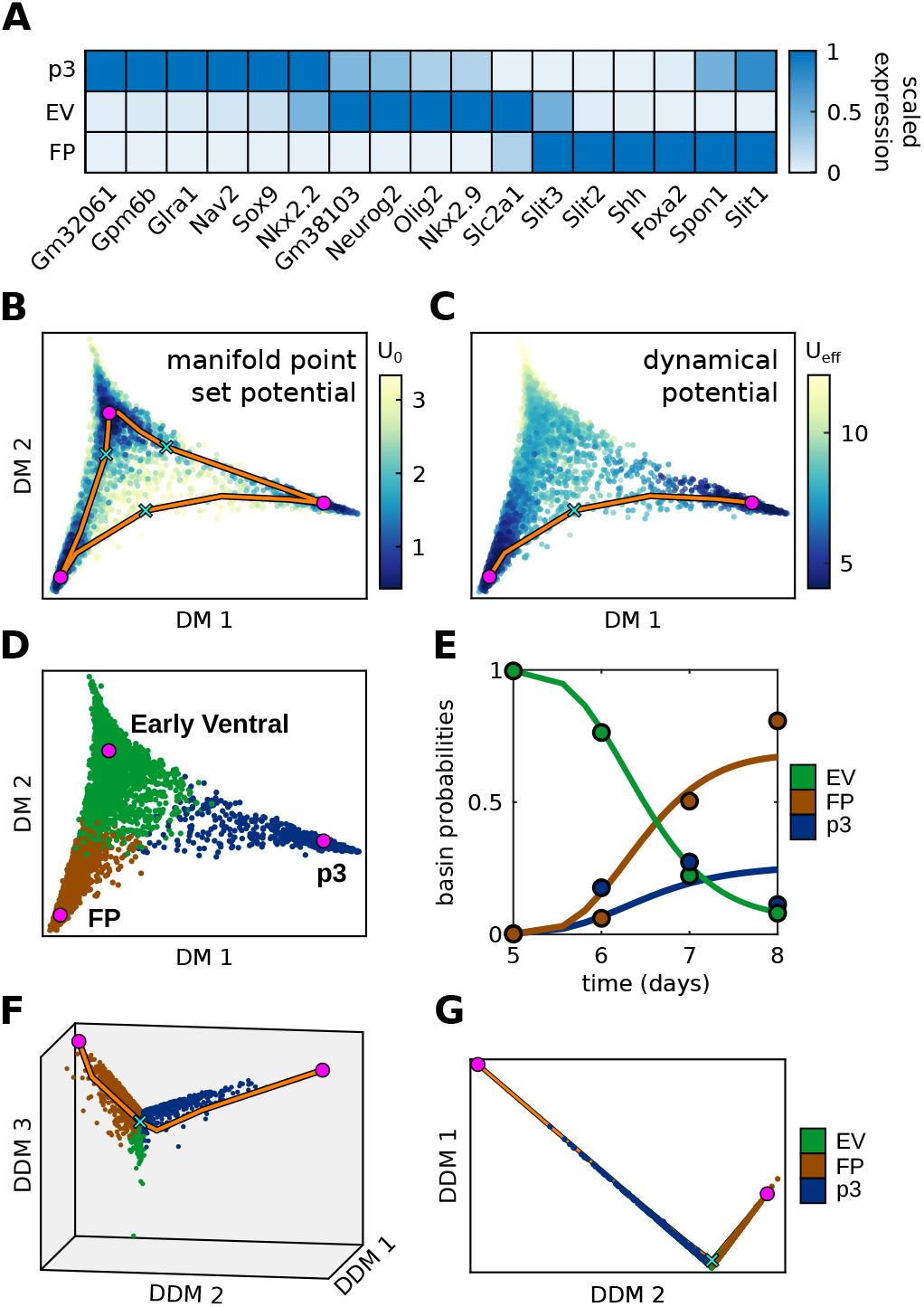
Fitting procedure captures differentiation of ventral progenitors in scRNA-seq data. All data acquired at SAG 500 nM. (A) Average normalized gene expression over the 30 nearest neighbors to the density maxima identified in (B) of the 17 genes most indicative of the three ventral states present during Days 5 to 8 [26]. Expression is scaled to lie between [0, 1] within each column. (The term ‘early ventral’ is non standard and, for us, includes the ‘double positive’ states (Olig2+) in Fig 21 in [26] Supplement.) (B) The base potential *U*_*B*_ shown on the dynamic manifold 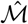 constructed in 17D and projected down to 2D using diffusion maps. Magenta circles are the density maxima (potential minima), orange curves are unstable manifolds of the base potential *U*_*B*_, and the cyan x’s are the corresponding saddles. (C) The dynamic potential, fit to the data for Days 5 to 8, destabilizes the early ventral (EV) state. (D) Assignment of points in 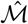 to basins of attraction corresponding to the density maxima in (B) (SI Movie 2). (E) The predicted basin proportions over time (curves) for the static fit potential in (C). Circles show the corresponding measured values. Systematic errors on Days 6 and 8 suggest the potential should be time dependent to favor floorplate (FP). (F-G) Two views of the 17D space projected onto the dynamical diffusion map coordinates based on the potential in (C), rather than the standard diffusion map methods projection shown in (B) to (D) (SI Movie 3).

We next quantified the observed time course of probability on the 17-dimensional manifold (Fig. 9E and SI Fig. S25). The cells, which are initially concentrated in the early ventral (EV) basin, flow out first towards the p3 attractor. By Day 7, the EV basin is empty and both the p3 and floor plate (FP) basins are populated, with about twice as much probability associated to the FP basin. This configuration is suggestive of the heteroclinic flip explored in the synthetic example. Indeed, the potential we fit has an unstable progenitor state (EV) and two locally stable terminal states, p3 and FP (Fig. 9C). The saddle connecting the two terminal wells is confirmed via moving least-squares fit to be an index-1 saddle (SI Table S8). The optimized *D*_*KL*_ *~* 0.3 bits for the measured and simulated distributions, compared to an *H*_*meas*_ *~* 10.5 bits (SI Table S7). Fit parameters were once again highly reproducible (modified coefficients of variation were ~5% see SI Fig. S27). This example demonstrates that our methodology is applicable to high-dimensional noisy data.

By Day 8, however, the p3 basin also empties, leaving a predominantly unimodal population of FP. We underestimate p3 and overestimate FP on Day 6, then do the reverse on Day 8 (Fig. 9E). These dynamics are inconsistent with a purely static heteroclinic flip, where probability moving into p3 would remain there rather flowing into FP. Hence, some additional signal at later time points must favor FP.

This example also illustrates the utility of *dynamical* diffusion maps based on the dynamic transition operator *T*_*ϵ*_ to dramatically simplify the data space. To construct these maps, we embed the data points in the right eigenvectors of the *T*_*ϵ*_ (treated as a left Markov matrix), as opposed to the standard diffusion map algorithm, which embeds the points in the left eigenvectors. Euclidean distances in these coordinates no longer have the interpretation of approximate diffusion distances [22], but they do highlight important dynamical features in the system. The two stable states in Fig. 9B each engage distinct eigenmodes of *T*_*ϵ*_ and appear orthogonal in Fig. 9F and G, whereas the unstable state is collapsed to the origin. These figures also suggest that the dynamics live locally in a two-dimensional space. See the SI Appendix S3.12 for more details.

## Discussion

We have developed a computational method for fitting landscape models directly to time-dependent single-cell data. Our approach accurately models how probability distributions evolve over time in high-dimensional state spaces, enabling us to exploit the intuitive geometry of developmental landscapes while preserving a direct connection to experimentally measured gene expression. Applied to a mESC model of ventral neural tube patterning, our method produced a family of dynamical potentials whose minima align with the canonical ventral neural progenitor types, linked by codimension-1 saddles. Changes in morphogen concentration sequentially create and annihilate stable states, matching the cascade of observed fate bifurcations *in vivo*. States appear to be arranged around a loop in the 5D state space (see also [26])

Our algorithms discretize the path integral form of the Fokker-Planck equation for gradient systems with additive noise. This formulation naturally captures the morphogen dependence of the dynamics through a one-parameter linear interpolation between two dynamic potentials (Fig. 6), in contrast to [10] where separate Hill functions had to be fit to each of the parameters in the interpolating polynomials. Even assuming an underlying MS structure, the result in Fig. 6 is a remarkable simplification and strongly suggests that gene epistasis can arise from minimal interactions of pathway components filtered through the nonlinear geometry of a developmental landscape, as previously shown in [6]. The ability to linearly interpolate potentials is another manifestation of canalization.

Quantitative models are valuable not only when they accurately reproduce experimental data, but also when their predictions deviate from observations in informative ways. For SAG 0–500, the static landscapes over-predicted induction of p3, supporting prior observations of a limited competence window for Shh signaling [13]. For SAG 500–0, the static potentials dramatically overestimated the final proportion of p0/p1. Together, these complementary deviations motivated the SAG 0–500–0 experiment, explicitly demonstrating that transient morphogen exposure irreversibly alters the underlying developmental landscape. Morphogen exposures do not commute, and their action is contingent on ‘developmental time’ even though the molecular timer is obscure in most instances. Dynamic data are crucial to proving that a simple model applies quantitatively.

Our methods extend to scRNA-seq data, but required a preprocessing step to focus on a subset of genes most relevant to the examined transition.

Our model in the resulting 17-dimensional space gave dynamical diffusion map embeddings that clearly revealed the attractor structure and stable states, and suggested subtle deviations from the simple flip scenario. In the future, we conjecture that focusing the RNA-seq on the 50-100 genes relevant to the particular system and gaining higher accuracy will yield greater insight than a larger gene panel.

The analysis of single-cell data is a vast enterprise and it is worthwhile to contextualize our contribution. Optimal transport approaches [16, 17] match cell state distributions at successive time points by minimizing transport costs, producing smooth trajectories that reveal the global topology of lineage bifurcations. For certain cost functions, these trajectories are equivalent to gradient flows that minimize the kinetic energy of the time-evolving distributions [28]. In contrast, our method infers dynamics by minimizing the curvature of the generating landscape by constraining the potential only along the one-dimensional unstable manifolds between attractors in gene expression space, a construction that explicitly exploits the canalization inherent in developmental dynamics. Our fitting parameters represent the potential values at the critical points. Population balance analysis [15, 29] also models drift-diffusion dynamics as a Markov process on a graph of data points, but assumes that the data are sampled from a steady-state distribution and is not applicable to the transient, non-stationary systems studied here.

Other recent methods also aim to fit MS dynamics to single-cell data. Dynamical landscape analysis [26] first clusters transcriptomics into identifiable states and then fits their occupancy over time using polynomials in two dimensions, as in [10]. Another approach builds MS systems using parameterized landscape neural networks to fit 2D potentials to time-dependent data [30]. Both methods leverage bifurcation theory to study how dynamics shift with signaling inputs, as we do, but differ in their construction and implementation.

RNA velocity methods infer transcriptional dynamics by fitting kinetic models to splicing or metabolic labeling data, thereby generating velocity vectors in gene expression space [25, 31, 32]. While these methods have the appeal of directly furnishing a velocity field, numerous steps in the analysis remain areas of active refinement [33, 34], and it is difficult to impose necessary topological constraints on the vector fields and connect them with bifurcation theory.

A GRN for the same ventral neural tube fates as we model was fit to *in vivo* data in [14]. They also measured the time-dependent signaling from endogenous Shh. This GRN exhibited bistability, which was validated experimentally. Recasting such models through the lens of our inferred landscapes, as done in other contexts in [3], could streamline the identification of minimal circuit architectures consistent with the data and focus attention on the critical regulatory interactions needed to explain observed dynamics.

Waddington’s landscape metaphor underpins many methods for processing high-dimensional gene expression data. Our approach is based on the MS formulation of these landscapes [3], a rigorous framework within dynamical systems theory, but only a hypothesis for real biological processes. In our method, we implicitly embrace the dimensionality reduction inherent to MS systems by constructing graph representations of dynamical manifolds directly from high-dimensional measured data, then constraining the interpolated potential along one-dimensional paths connecting density maxima. Dynamic signaling experiments designed to explore gene expression trajectories around saddle points offer promising routes to demonstrate that developmental dynamics can be captured by low-dimensional representations. Computational geometry frameworks for manifold learning [35–37] provide a complementary route to explicitly determine the dimensionality needed to describe such developmental processes. Ultimately, realizing the full potential of Waddington’s insight will require definitive evidence that developmental dynamics are inherently low-dimensional, with MS systems and bifurcation theory likely playing critical roles in this demonstration.

## Acknowledgments

DJC was supported in part by the Helen Hay Whitney foundation. EDS was supported in part by NSF Grant 2013131. The authors thank David Rand and Marine Fontaine for innumerable discussions and sharing their paper and data prior to publication. Francis Corson critically read a draft of the manuscript. JB and MJD were supported by the Francis Crick Institute, which receives its core funding from Cancer Research UK, the UK Medical Research Council and the Wellcome Trust (all through CC001051). MJD was additionally supported by the Wellcome Trust Career Development Award (227326/Z/23/Z).

