## Supplementary Information for "Reconstructing Waddington’s Landscape from Data"

### S1 Theoretical Background

The purpose of this text is to provide a comprehensive theoretical background that simultaneously motivates specific choices in our mathematical construction, demonstrates that our formalism correctly captures the dynamical features of the biological systems it models, and highlights connections between ours and existing methods and perspectives. We begin in Appendix S1.1 by elaborating on the dynamics of probability density distributions for stochastic dynamical systems, as introduced in the main text. We then give a brief explanation of kernel density estimation (KDE) in Appendix S1.2, a crucial method in our work that allows us to infer probability distributions defined on our discrete manifold from time courses of point cloud data. This is followed by a discussion of diffusion maps in Appendix S1.3. In this work, diffusion maps are used for nearly all visualizations of high-dimensional data. There is also a connection between the theory of stochastic processes underlying diffusion maps and our construction, which is highlighted in subsequent sections. Next, we provide a detailed description of our construction of a discrete Laplace operator in Appendix S1.4. This operator is an essential component of understanding general drift-diffusion processes and naturally encodes the geometry of our discrete manifold. It also enables robust computation of our interpolated potential, which we explain in Appendix S1.5. Finally, in Appendix S1.6, we explicitly define our discrete dynamical transition operator and volume elements. We also offer a simplified proof of convergence in the continuum limit that relies on the theory developed in Appendices S1.3 and S1.4.

#### S1.1 Mathematical preliminaries

For completeness and ease of discussion, we restate some of the mathematical exposition in the main text. Let $\mathbf{x}(t) \in \mathbb{R}^d$ denote the time-dependent state of a single cell. We assume that the time-dependence of the state is governed by the following Langevin equation:

$$\dot{\mathbf{x}}(t) = \mathbf{v}(\mathbf{x}, t\,|\,\boldsymbol{\theta}) + \boldsymbol{\eta}(t) = -\mathbf{g}^{-1}(\mathbf{x}, t\,|\,\boldsymbol{\theta})\,\nabla U(\mathbf{x}, t\,|\,\boldsymbol{\theta}) + \boldsymbol{\eta}(t). \tag{S1}$$

Here, the drift term $\mathbf{v} = -\mathbf{g}^{-1}\nabla U$ is assumed to be gradient-like for a given metric tensor $\mathbf{g}$ and potential $U$. The stochastic term $\boldsymbol{\eta}(t)$ is standard white noise with mean $\langle \boldsymbol{\eta}(t) \rangle = 0$ and covariance

$$\langle \eta_\alpha(t)\, \eta_\beta(t') \rangle = 2D\, \delta_{\alpha\beta}\, \delta(t - t'), \tag{S2}$$

where Greek indices run over the dimensions $1, \ldots, d$ and $D$ is the (positive) diffusion constant. We denote by $\boldsymbol{\theta}$ a vector of time-dependent parameters that control the topography of the dynamical landscape. We frequently omit any explicit dependence of the observables on $\boldsymbol{\theta}$ for the sake of notational simplicity. The biological systems we model are typically not stationary, and there is no explicit link between $U$ in Eq. (S1) and any equilibrium distribution (by design $\mathbf{g}$ does not appear in Eq. (S2)).

In the deterministic limit, the trajectories $\mathbf{x}(t)$ may evolve along a low-dimensional submanifold $\mathcal{M} \subset \mathbb{R}^d$, but high-dimensional noise will generically blur this confinement. In this work, we do not attempt to explicitly encode the geometry of $\mathcal{M}$ and instead extend the stochastic dynamics to $\mathbb{R}^d$ by defining a $U(\mathbf{x})$ that keeps trajectories close to $\mathcal{M}$. In our convention, $\mathbf{g}$ is not the metric of the dynamical manifold, i.e., it does not define inner products or measure distances on $\mathcal{M}$. It is a purely dynamical object that determines the trajectories $\mathbf{x}(t)$.

The probability density $p(\mathbf{x}, t\,|\,\boldsymbol{\theta})$ to measure a cell with state $\mathbf{x}$ at time $t$ evolves according to the Fokker-Planck or forward Kolmogorov equation [1]:

$$\frac{\partial p}{\partial t} = -\nabla \cdot (p\,\mathbf{v}) + D\,\Delta\, p = [-(\nabla \cdot \mathbf{v}) - \mathbf{v} \cdot \nabla + D\,\Delta]\; p = \mathcal{L}^{\dagger}\, p, \tag{S3}$$

where $\Delta$ denotes the Laplacian on $\mathbb{R}^d$. It can be shown that Eq. (S3) is equivalent to the following recursion relation for short time intervals $\epsilon << 1$ [1]:

$$\begin{aligned} p(\mathbf{x}, t+\epsilon) &= \int d\mathbf{y}\, T(\mathbf{x}, t+\epsilon \,|\, \mathbf{y}, t)\, p(\mathbf{y}, t) \\ &= \int d\mathbf{y}\, (4\pi D\epsilon)^{-d/2} \exp\left[-\frac{||\mathbf{x}-\mathbf{y}-\epsilon\,\mathbf{v}(\mathbf{y},t)||^2}{4\,D\,\epsilon}\right] p(\mathbf{y}, t) \\ &= \int d\mathbf{y}\, (4\pi D\epsilon)^{-d/2} \times \\ &\quad \exp\left[-\frac{||\mathbf{x}-\mathbf{y}||^2}{4\,D\,\epsilon} - \frac{(\mathbf{x}-\mathbf{y})\cdot\mathbf{g}^{-1}(\mathbf{y},t)\nabla U(\mathbf{y},t)}{2D} - \frac{\epsilon\,||\mathbf{g}^{-1}(\mathbf{y},t)\nabla U(\mathbf{y},t)||^2}{4\,D}\right] p(\mathbf{y}, t), \end{aligned} \tag{S4}$$

where $T(\mathbf{x}, t \,|\, \mathbf{y}, s)$ denotes the transition probability from state $\mathbf{y}$ at time $s \le t$ to state $\mathbf{x}$ at time $t$. All inner products and norms in Eq. (S4) are defined with respect to the Euclidean metric on $\mathbb{R}^d$. In the last equality, we simply substituted the gradient-like drift $\mathbf{v} = -\mathbf{g}^{-1}\nabla U$ and expanded the inner product. In what follows, we adopt the simplified notation $T(\mathbf{x}, t \,|\, \mathbf{y}, s) := T_\epsilon(\mathbf{x}, \mathbf{y})$, suppressing the dependence on time which enters only through $\mathbf{v}$ and $\boldsymbol{\theta}$ for fixed $\epsilon = t - s$. Note that Eq. (S3) is an initial value problem – solving it yields the density $p(\mathbf{x}, t \,|\boldsymbol{\theta})$ given the initial density $p(\mathbf{x}, 0 \,|\boldsymbol{\theta})$. Consider some arbitrary, smooth function $\psi(\mathbf{x}(t)) : \mathbb{R}^d \to \mathbb{R}$ of the random variable $\mathbf{x}(t)$ and let

$$u(\mathbf{y}, s) = \mathbb{E}\left[\psi(\mathbf{x}(t)) \,|\, \mathbf{x}(s) = \mathbf{y}\right] = \int d\mathbf{x}\, \psi(\mathbf{x})\, T(\mathbf{x}, t \,|\, \mathbf{y}, s). \tag{S5}$$

It is straightforward to show that $u(\mathbf{y}, s)$ obeys the backward Kolmogorov equation:

$$-\frac{\partial u}{\partial s} = \left[\mathbf{v}\cdot\nabla + D\,\Delta\right] u = \mathcal{L}\, u. \tag{S6}$$

The backward Kolmogorov equation is a final value problem whose solution $u(\mathbf{y}, s)$ is the expected value of $\psi(\mathbf{x}(t))$ given that the stochastic trajectory $\mathbf{x}(t)$ began at $\mathbf{x}(s) = \mathbf{y}$ at time $s < t$. We note that the generators of the forward and backward Kolmogorov equations are adjoint to each other, i.e. $\langle \mathcal{L}\, u,\, p\rangle = \langle u,\, \mathcal{L}^\dagger\, p\rangle$. See [2], for instance, for a derivation and a discussion of the Kolmogorov equations from an operator theoretic viewpoint.

Our numerical methods produce discrete approximations to the solutions of Eq. (S3) on high-dimensional, unstructured point clouds. Specifically, given a set of $N$ points $\hat{\mathcal{M}} = \{\mathbf{x}_i\}_{i=1}^N$, we leverage Eq. (S4) to construct a discrete-time Markov process on $\hat{\mathcal{M}}$. The result is a time course of probability vectors $P(\mathbf{x}_i, t_k) \in \mathbb{R}^N$ with $\sum_{i=1}^N P(\mathbf{x}_i, t_k) = 1$ and $t_k = k\,\epsilon$. The following sections will provide numerical details for implementing these discrete Markov processes, some applications of the resulting transition operator, and various methods needed to process and analyze single-cell data within our framework, including the construction of discrete differential operators on $\hat{\mathcal{M}}$.

#### S1.2 Kernel density estimation

Kernel density estimation (KDE) is a non-parametric method for approximating continuous probability density functions from unstructured sets of discrete random samples [3, 4]. KDE is crucial in our method, since it allows us to infer probability distributions $P(\mathbf{x}_i, t_k)$ on our discrete manifold $\hat{\mathcal{M}}$ from single-cell data. KDE also provides a conceptual starting point for approximating continuous drift-diffusion processes from discrete samples (see Appendix S1.3). Consider a set of $d$-dimensional points $\{\mathbf{x}_i\}_{i=1}^N$ sampled from a probability density $p(\mathbf{x})$. The kernel density estimate for the probability density at some point $\mathbf{z} \in \mathbb{R}^d$ is

$$\hat{p}(\mathbf{z}) = \frac{1}{N\,|\mathbf{H}|} \sum_{i=1}^{N} \mathcal{K}(\mathbf{H}^{-1}(\mathbf{z} - \mathbf{x}_i)) \tag{S7}$$

for a given $d \times d$ nonsingular bandwidth matrix $\mathbf{H}$ and a multivariate kernel function $\mathcal{K} : \mathbb{R}^d \to \mathbb{R}$. For our purposes, we always use a simple normalized isotropic Gaussian kernel

$$\hat{p}(\mathbf{z}) = \frac{1}{N\,(4\pi h)^{d/2}} \sum_{i=1}^{N} \exp\left[-||\mathbf{z} - \mathbf{x}_i||^2 \,/\, 4\,h\right] \tag{S8}$$

with scalar bandwidth $h$. This estimate will converge to the true density in the continuum limit ($N \to \infty$, $h \to 0$). The rate of convergence as a function of sample number and bandwidth has been extensively studied [4] and typically many more samples are required in higher dimensions to achieve a high quality estimate. Attempts have also been made to provide heuristics for selecting bandwidths that approach the optimal convergence rates, however, the effectiveness of these heuristics can vary drastically between data sets. In practice, after inspecting the distribution of pairwise distances in $\hat{\mathcal{M}}$, we manually select the bandwidth to achieve a base density $p_B$ that varies smoothly as one moves from point to point along the discrete manifold.

##### S1.3 Diffusion maps

Diffusion maps are a nonlinear dimensional reduction technique that leverages the properties of drift-diffusion processes to extract low-dimensional embeddings of high-dimensional data [5–7]. Although diffusion maps are not used directly in our framework to infer or analyze developmental dynamics, we do use them extensively to visualize high-dimensional single-cell data. The choice of dimensional reduction technique for visualization is somewhat arbitrary here, since all of our actual analysis is done in the high-dimensional space where the data lives. However, we prefer diffusion maps since, by construction, they preserve elements of the dynamical geometry of the data (see below and Eq. (S20) in particular). This can be contrasted with other methods, such as UMAP [8], which preserves the topological connectivity of data at the expense of geometry. More substantially, the theory of stochastic processes that underlies diffusion maps directly enables our construction of a discrete Laplace operator, which we use, for instance, to compute our interpolated potential $U_I$. We also use this theory to prove the convergence of our dynamical transition operator in the continuum limit. Finally, diffusion maps directly inspire our dynamical diffusion map method for low-dimensional embedding (see Fig. 9F and G and Appendix S3.12). For these reasons, we therefore offer a brief explanation of the standard diffusion map algorithm. Suppose that the point set $\hat{\mathcal{M}} = \{\mathbf{x}_i\}_{i=1}^N$ is sampled from a Riemannian manifold $\mathcal{M} \subset \mathbb{R}^d$ according to the (non-uniform) probability density $q(\mathbf{x}) : \mathcal{M} \to \mathbb{R}$. Roughly following the notation in [6], let

$$K_\epsilon(\mathbf{x}_i, \mathbf{x}_j) = \exp\left[-||\mathbf{x}_i - \mathbf{x}_j||^2 \,/\, 4\,D\,\epsilon\right] \tag{S9}$$

denote the components of an affinity matrix $\mathbf{K}_\epsilon$ given by a Gaussian kernel with bandwidth $D\,\epsilon$. Then $q_\epsilon(\mathbf{x}_i) = \sum_{j=1}^N K_\epsilon(\mathbf{x}_i, \mathbf{x}_j)$ is an unnormalized kernel density estimate for $q(\mathbf{x}_i)$. Note that in the standard diffusion map construction there is no explicit diffusion constant $D$, but we keep it here since it has physical meaning in our framework. We further define

$$K_\epsilon^{(\alpha)}(\mathbf{x}_i, \mathbf{x}_j) = \frac{K_\epsilon(\mathbf{x}_i, \mathbf{x}_j)}{q_\epsilon^\alpha(\mathbf{x}_i)\, q_\epsilon^\alpha(\mathbf{x}_j)}, \qquad d_\epsilon^{(\alpha)}(\mathbf{x}_i) = \sum_{j=1}^{N} K_\epsilon^{(\alpha)}(\mathbf{x}_i, \mathbf{x}_j) \tag{S10}$$

for some $\alpha > 0$. The $d_\epsilon^{(\alpha)}(\mathbf{x}_i)$ are the nonzero elements of the diagonal degree matrix $\mathbf{D}_\epsilon^{(\alpha)}$ associated to the symmetric matrix $\mathbf{K}_\epsilon^{(\alpha)}$. Finally, we define the Markov transition kernels

$$\begin{aligned} M_f^{(\alpha)}(\mathbf{x}_i, \mathbf{x}_j) &= \Pr(\mathbf{x}(t+\epsilon) = \mathbf{x}_i \,|\, \mathbf{x}(t) = \mathbf{x}_j) = \frac{K_\epsilon^{(\alpha)}(\mathbf{x}_i, \mathbf{x}_j)}{d_\epsilon^{(\alpha)}(\mathbf{x}_j)}, \\ M_b^{(\alpha)}(\mathbf{x}_i, \mathbf{x}_j) &= \frac{K_\epsilon^{(\alpha)}(\mathbf{x}_i, \mathbf{x}_j)}{d_\epsilon^{(\alpha)}(\mathbf{x}_i)} = M_f^{(\alpha)}(\mathbf{x}_j, \mathbf{x}_i), \\ M_s^{(\alpha)}(\mathbf{x}_i, \mathbf{x}_j) &= \frac{K_\epsilon^{(\alpha)}(\mathbf{x}_i, \mathbf{x}_j)}{\sqrt{d_\epsilon^{(\alpha)}(\mathbf{x}_i)\, d_\epsilon^{(\alpha)}(\mathbf{x}_j)}}. \end{aligned} \tag{S11}$$

These operators define a one-parameter family of drift-diffusion processes that depend $\alpha$. In this and subsequent sections, the specific process corresponding to $\alpha = 1$ is most relevant to us, but we include a general treatment for completeness. If we let $q(\mathbf{x}_i, t)$ represent the probability density of finding the system at location $\mathbf{x}_i$ at time $t$, then $q(\mathbf{x}_i, t+\epsilon) = \sum_j M_f^{(\alpha)}(\mathbf{x}_i, \mathbf{x}_j)\, q(\mathbf{x}_j, t)$ is the evolution of this probability to time $t + \epsilon$. Note that $\mathbf{M}_f^{(\alpha)}$ is deemed a forward operator, in the sense that it is a normalized Markov matrix that evolves probabilities forward in time and conserves probability (this follows from $\sum_i M_f^{(\alpha)}(\mathbf{x}_i, \mathbf{x}_j) = 1$). Conversely, if $g(\mathbf{x}_i)$ is an arbitrary function defined on the point set, then the action of the corresponding backwards kernel $\sum_j M_b^{(\alpha)}(\mathbf{x}_i, \mathbf{x}_j)\, g(\mathbf{x}_j)$ gives us the average value of that function, evaluated at $t = \epsilon$, for a random walk that started at $\mathbf{x}_i$ at $t = 0$. Refer back to Appendix S1.1 for more details. $M_s^{(\alpha)}(\mathbf{x}_i, \mathbf{x}_j)$ is a symmetric transition kernel that makes some subsequent calculations simpler. Crucially, these diffusion map transition operators must not be confused with our dynamical transition operator, which we will define explicitly in Appendix S1.6. A fundamental assumption of the standard diffusion map method is that data points are sampled from an equilibrium distribution $q(\mathbf{x})$, which is clearly incompatible with non-steady state developmental dynamics. The connection to drift-diffusion processes here is essentially just a way of viewing $\hat{\mathcal{M}}$ as the output of a Markov Chain Monte Carlo (MCMC) process sampling $q(\mathbf{x})$. This is also why we specifically use the symbol $q$ for the these sampling distributions – to avoid any confusion with the dynamical probability density distributions $p$ whose time evolution is governed by our discrete transition operator.

These operators naturally induce a distance between any two data points based on their dynamic proximity. Coifman and Lafon define the *diffusion distance* at time $t_k = k\,\epsilon$ as

$$D_{t_k}^2(\mathbf{x}_i, \mathbf{x}_j) = \sum_{\mathbf{z}} (q(\mathbf{z}, t_k \,|\, \mathbf{x}_i) - q(\mathbf{z}, t_k \,|\, \mathbf{x}_j))^2\, w(\mathbf{z}), \tag{S12}$$

where $q(\mathbf{z}, t_k \,|\, \mathbf{x}_i)$ is the probability of random walk that began at $\mathbf{x}_i$ at $t = 0$ ending up at location $\mathbf{z}$ at $t_k$ and $w(\mathbf{z})$ is a positive weight function. This distance can be explicitly written in terms of the eigenvectors of the matrix $\mathbf{M}_f^{(\alpha)}$. To start, since $\mathbf{M}_s^{(\alpha)}$ is a real symmetric matrix, it has $N$ real eigenvalues $\lambda_j$ and we can choose its eigenvectors $\hat{\mathbf{v}}_j$ to form an orthonormal basis of $\mathbb{R}^N$. We can define the right eigenvectors $\boldsymbol{\varphi}_j$ of $\mathbf{M}_f^{(\alpha)}$ through

$$\begin{aligned} \mathbf{M}_s^{(\alpha)}\, \hat{\mathbf{v}}_j = \left[ (\mathbf{D}_\epsilon^{(\alpha)})^{-1/2}\, \mathbf{M}_f^{(\alpha)}\, (\mathbf{D}_\epsilon^{(\alpha)})^{1/2} \right] \hat{\mathbf{v}}_j = \lambda_j\, \hat{\mathbf{v}}_j \\ \implies \mathbf{M}_f^{(\alpha)}\, \boldsymbol{\varphi}_j = \lambda_j\, \boldsymbol{\varphi}_j \quad \text{for} \quad \boldsymbol{\varphi}_j = (\mathbf{D}_\epsilon^{(\alpha)})^{1/2}\, \hat{\mathbf{v}}_j. \end{aligned} \tag{S13}$$

We can similarly define its left eigenvectors $\boldsymbol{\psi}_j$ through

$$\begin{aligned} \hat{\mathbf{v}}_j^T \, \mathbf{M}_s^{(\alpha)} &= \hat{\mathbf{v}}_j^T \, \left[ (\mathbf{D}_\epsilon^{(\alpha)})^{-1/2} \, \mathbf{M}_f^{(\alpha)} \, (\mathbf{D}_\epsilon^{(\alpha)})^{1/2} \right] = \lambda_j \, \hat{\mathbf{v}}_j^T \\ \implies \; \boldsymbol{\psi}_j^T \, \mathbf{M}_f^{(\alpha)} &= \lambda_j \, \boldsymbol{\psi}_j^T \quad \text{for} \quad \boldsymbol{\psi}_j = (\mathbf{D}_\epsilon^{(\alpha)})^{-1/2} \, \hat{\mathbf{v}}_j. \end{aligned} \tag{S14}$$

Clearly the $\boldsymbol{\varphi}_j$ and the $\boldsymbol{\psi}_j$ share eigenvalues with the $\hat{\mathbf{v}}_j$ and are biorthonormal, i.e. $\boldsymbol{\varphi}_i^T \, \boldsymbol{\psi}_j = \boldsymbol{\psi}_i^T \, \boldsymbol{\varphi}_j = \delta_{ij}$. It is easy to show that the eigendecomposition of $\mathbf{M}_f^{(\alpha)}$ implies

$$\mathbf{M}_f^{(\alpha)} = \boldsymbol{\Phi} \, \boldsymbol{\Lambda} \, \boldsymbol{\Psi}^T \implies \left( \mathbf{M}_f^{(\alpha)} \right)^k = \boldsymbol{\Phi} \, \boldsymbol{\Lambda}^k \, \boldsymbol{\Psi}^T, \tag{S15}$$

where $\boldsymbol{\Phi}$ is the matrix whose columns are the $\boldsymbol{\varphi}_j$, $\boldsymbol{\Psi}$ is the matrix whose columns are the $\boldsymbol{\psi}_j$, and $\boldsymbol{\Lambda}$ is the diagonal matrix with $\Lambda_{ii} = \lambda_i$. Therefore, the probabilities

$$q(\mathbf{z}, t_k \,|\, \mathbf{x}_i) = \left[ \left( \mathbf{M}_f^{(\alpha)} \right)^k \hat{\mathbf{e}}(\mathbf{x}_i) \right] (\mathbf{z}) = \sum_\ell \lambda_\ell^k \, \varphi_\ell(\mathbf{z}) \, \psi_\ell(\mathbf{x}_i), \tag{S16}$$

where $\hat{\mathbf{e}}(\mathbf{x}_i) \in \mathbb{R}^N$ is the vector whose value corresponding to $\mathbf{x}_i$ is one and is zero everywhere else. Eq. (S12) can therefore be written as

$$\begin{aligned} D_{t_k}^2(\mathbf{x}_i, \mathbf{x}_j) &= \sum_{\mathbf{z}} (\sum_\ell \lambda_\ell^k \, \varphi_\ell(\mathbf{z}) \, (\psi_\ell(\mathbf{x}_i) - \psi_\ell(\mathbf{x}_j)))^2 \, w(\mathbf{z}) \\ &= \sum_\ell \sum_{\ell'} \lambda_\ell^k \, \lambda_{\ell'}^k \, (\psi_\ell(\mathbf{x}_i) - \psi_\ell(\mathbf{x}_j)) \, (\psi_{\ell'}(\mathbf{x}_i) - \psi_{\ell'}(\mathbf{x}_j)) \left( \sum_{\mathbf{z}} \varphi_\ell(\mathbf{z}) \, \varphi_{\ell'}(\mathbf{z}) \, w(\mathbf{z}) \right). \end{aligned} \tag{S17}$$

For the specific choice $w(\mathbf{z}) = 1/d_\epsilon^{(\alpha)}(\mathbf{z})$ we have

$$\sum_{\mathbf{z}} \varphi_\ell(\mathbf{z}) \, \varphi_{\ell'}(\mathbf{z}) \, w(\mathbf{z}) = \sum_{\mathbf{z}} \varphi_\ell(\mathbf{z}) \, \frac{\varphi_{\ell'}(\mathbf{z})}{d_\epsilon^{(\alpha)}(\mathbf{z})} = \sum_{\mathbf{z}} \varphi_\ell(\mathbf{z}) \, \psi_{\ell'}(\mathbf{z}) = \delta_{\ell\ell'}. \tag{S18}$$

In the third equality, we used the definitions in Eq. (S13) and Eq. (S14) and, in the fourth equality, we used the biorthonormality of the $\boldsymbol{\varphi}_\ell$ and the $\boldsymbol{\psi}_\ell$. Finally, the diffusion distance becomes

$$D_{t_k}^2(\mathbf{x}_i, \mathbf{x}_j) = \sum_{\ell > 0} \lambda_\ell^{2k} \, (\psi_\ell(\mathbf{x}_i) - \psi_\ell(\mathbf{x}_j))^2 = \sum_{\ell > 0} (\lambda_\ell^k \, \psi_\ell(\mathbf{x}_i) - \lambda_\ell^k \, \psi_\ell(\mathbf{x}_j))^2. \tag{S19}$$

Note that we can restrict our sum to $\ell > 0$ since $\psi_0(\mathbf{x}_i)$ is constant for all $\mathbf{x}_i$ (which follows directly from the fact that all column sums $\sum_i M_f^{(\alpha)}(\mathbf{x}_i, \mathbf{x}_j) = 1$). If we define the diffusion coordinates at $t_k$

$$\tilde{\boldsymbol{\Psi}}_k(\mathbf{x}_i) = \left( \lambda_1^k \, \psi_1(\mathbf{x}_i), \, \lambda_2^k \, \psi_2(\mathbf{x}_i), \, \ldots, \, \lambda_N^k \, \psi_N(\mathbf{x}_i) \right), \tag{S20}$$

we see that the Euclidean distance $||\tilde{\boldsymbol{\Psi}}_k(\mathbf{x}_i) - \tilde{\boldsymbol{\Psi}}_k(\mathbf{x}_j)|| = D_{t_k}(\mathbf{x}_i, \mathbf{x}_j)$. Since all column sums $\sum_i M_f^{(\alpha)}(\mathbf{x}_i, \mathbf{x}_j) = 1$, by the Gershgorin circle theorem [9], the eigenvalues of $\mathbf{M}_f^{(\alpha)}$ must satisfy $1 = \lambda_0 \geq \lambda_1 \geq \lambda_2 \geq \ldots \geq \lambda_N \geq 0$. If these eigenvalues decrease quickly, i.e. there is a large spectral gap, then we can embed our data in a low-dimensional space $\hat{\boldsymbol{\Psi}}_k(\mathbf{x}_i) = \left( \lambda_1^k \, \psi_1(\mathbf{x}_i), \, \ldots, \, \lambda_n^k \, \psi_n(\mathbf{x}_i) \right)$ ($n = 2$ or 3) where $||\hat{\boldsymbol{\Psi}}_k(\mathbf{x}_i) - \hat{\boldsymbol{\Psi}}_k(\mathbf{x}_j)|| \approx D_{t_k}(\mathbf{x}_i, \mathbf{x}_j)$.

#### S1.4 Discretization of the Laplace-Beltrami operator

Here we discuss the construction of discrete approximations to the Laplace-Beltrami operator, i.e. a generalization of the standard Laplace operator to Riemannian manifolds. This operator plays a central role in characterizing drift–diffusion processes and naturally reflects the geometry of the underlying discrete manifold $\hat{\mathcal{M}}$. It also enables stable and robust computation of the interpolated potential $U_I$, as detailed in Appendix S1.5. Several algorithms exist that build Laplacians for unstructured point clouds that sample curved submanifolds of Euclidean space [10–12]. No discrete approximation simultaneously captures all the mathematical properties of its continuous counterpart, and care must be taken to choose a numerical construction that captures the appropriate properties for the geometry processing task at hand [13]. For our purposes, we use a construction based on the diffusion map algorithm (see Appendix S1.3).

In this section, we first provide a simplified proof of the convergence of the infinitesimal generator of diffusion processes to a family of generalized Laplacians defined over $\mathbb{R}^d$. The result is a discrete Laplacian that can be explicitly written in terms of the operators defined in Appendix S1.3. We next provide a schematic derivation of the generic form of discrete Laplacians using general geometric reasoning. We will see that our discrete Laplacian, derived from from the alternative perspective of drift-diffusion processes, matches this structure precisely and naturally yields a geometrically motivated volume element that we can associate to points in $\hat{\mathcal{M}}$. Finally, we leverage this structure to formulate the Laplacian in a way that specifically improves numerical stability.

##### S1.4.1 Diffusion processes and generalized Laplacians

Here we show that the family of diffusion processes defined in the diffusion map construction, which depend on the scalar parameter $\alpha$, converges to a corresponding family of generalized Laplacians in the continuum limit ($N \to \infty, \epsilon \to \infty$). In this and subsequent sections, the specific process corresponding to $\alpha = 1$ is most relevant to us, but we include a general treatment for completeness. What follows is a simplified proof for functions and operators defined over $\mathbb{R}^d$. A more sophisticated proof for general embedded Riemannian manifolds $\mathcal{M} \subset \mathbb{R}^d$ is given in [6]. Even more general proofs for different kernels are given in [14]. To begin, we state without proof the following result

$$\int_{\mathbb{R}^d} K_\epsilon(\mathbf{x},\mathbf{y})\, f(\mathbf{y})\, d\mathbf{y} = (4\,\pi\, D\,\epsilon)^{d/2}\, \left(f(\mathbf{x}) + D\,\epsilon\,\Delta f(\mathbf{x}) + \mathcal{O}(\epsilon^2)\right) \tag{S21}$$

where $f(\mathbf{x}) \in \mathcal{C}^3(\mathbb{R}^d)$ is a three times continuously differentiable function and $K_\epsilon$ is defined in Eq. (S9). This result is a special case of Lemma 8 in Appendix B of [6], up to a sign change which reflects our convention that $\Delta$ should be negative semi-definite. Let $q(\mathbf{x})$ be a probability density defined on $\mathbb{R}^d$. Eq. (S21) implies that

$$\begin{aligned} q_\epsilon(\mathbf{x}) &:= \int_{\mathbb{R}^d} K_\epsilon(\mathbf{x},\mathbf{y})\, q(\mathbf{y})\, d\mathbf{y} = (4\,\pi\, D\,\epsilon)^{d/2}\, \left(q(\mathbf{x}) + D\,\epsilon\,\Delta q(\mathbf{x}) + \mathcal{O}(\epsilon^2)\right) \\ &\implies q_\epsilon^{-\alpha}(\mathbf{x}) = (4\,\pi\, D\,\epsilon)^{-\alpha d/2}\, q^{-\alpha}(\mathbf{x}) \left(1 - \alpha\, D\,\epsilon\, \frac{\Delta\, q(\mathbf{x})}{q(\mathbf{x})} + \mathcal{O}(\epsilon^2)\right). \end{aligned} \tag{S22}$$

With a slight abuse of notation we define the integral operator $K_\epsilon^{(\alpha)} f(\mathbf{x})$ to be

$$\begin{aligned} K_\epsilon^{(\alpha)} f(\mathbf{x}) &:= \int_{\mathbb{R}^d} K_\epsilon^{(\alpha)}(\mathbf{x},\mathbf{y})\, f(\mathbf{y})\, q(\mathbf{y})\, d\mathbf{y} = \int_{\mathbb{R}^d} \frac{K_\epsilon(\mathbf{x},\mathbf{y})}{q_\epsilon^\alpha(\mathbf{x})\, q_\epsilon^\alpha(\mathbf{y})}\, f(\mathbf{y})\, q(\mathbf{y})\, d\mathbf{y} \\ &= \frac{1}{q_\epsilon^\alpha(\mathbf{x})} \int_{\mathbb{R}^d} K_\epsilon(\mathbf{x},\mathbf{y})\, q_\epsilon^{-\alpha}(\mathbf{y})\, f(\mathbf{y})\, q(\mathbf{y})\, d\mathbf{y} \\ &= \frac{(4\,\pi\, D\,\epsilon)^{-\alpha d/2}}{q_\epsilon^\alpha(\mathbf{x})} \int_{\mathbb{R}^d} K_\epsilon(\mathbf{x},\mathbf{y})\, q^{1-\alpha}(\mathbf{y})\, f(\mathbf{y}) \left(1 - \alpha\, D\,\epsilon\, \frac{\Delta\, q(\mathbf{x})}{q(\mathbf{x})} + \mathcal{O}(\epsilon^2)\right) d\mathbf{y} \\ &= \frac{(4\,\pi\, D\,\epsilon)^{(1-\alpha)d/2}}{q_\epsilon^\alpha(\mathbf{x})} \left(q^{1-\alpha}(\mathbf{x})\, f(\mathbf{x}) + D\,\epsilon\,\Delta\left(q^{1-\alpha}(\mathbf{x})\, f(\mathbf{x})\right) - \alpha\, D\,\epsilon\, q^{1-\alpha}(\mathbf{x})\, f(\mathbf{x})\, \frac{\Delta\, q(\mathbf{x})}{q(\mathbf{x})} + \mathcal{O}(\epsilon^2)\right) \end{aligned} \tag{S23}$$

and

$$\begin{aligned} d_\epsilon^{(\alpha)}(\mathbf{x}) &:= K_\epsilon^{(\alpha)}\, 1 \\ &= \frac{(4\,\pi\, D\,\epsilon)^{(1-\alpha)d/2}}{q_\epsilon^\alpha(\mathbf{x})} \left(q^{1-\alpha}(\mathbf{x}) + D\,\epsilon\,\Delta\, q^{1-\alpha}(\mathbf{x}) - \alpha\, D\,\epsilon\, q^{1-\alpha}(\mathbf{x})\, \frac{\Delta\, q(\mathbf{x})}{q(\mathbf{x})} + \mathcal{O}(\epsilon^2)\right). \end{aligned} \tag{S24}$$

Clearly, $K_\epsilon^{(\alpha)} f(\mathbf{x})$ is just the expected value of the product $K_\epsilon^{(\alpha)}(\mathbf{x},\mathbf{y})\, f(\mathbf{y})$ with respect to the distribution $q(\mathbf{y})$. We can use these quantities to construct the associated backwards integral transition operator

$$\begin{aligned} M_b^{(\alpha)} f(\mathbf{x}) &:= \frac{K_\epsilon^{(\alpha)} f(\mathbf{x})}{d_\epsilon^{(\alpha)}(\mathbf{x})} = \int_{\mathbb{R}^d} \frac{K_\epsilon^{(\alpha)}(\mathbf{x},\mathbf{y})}{d_\epsilon^{(\alpha)}(\mathbf{x})}\, f(\mathbf{y})\, q(\mathbf{y})\, d\mathbf{y} \\ &= \frac{q^{1-\alpha}(\mathbf{x})\, f(\mathbf{x}) + D\,\epsilon\,\Delta\left(q^{1-\alpha}(\mathbf{x})\, f(\mathbf{x})\right) - \alpha\, D\,\epsilon\, q^{1-\alpha}(\mathbf{x})\, f(\mathbf{x})\, \frac{\Delta\, q(\mathbf{x})}{q(\mathbf{x})} + \mathcal{O}(\epsilon^2)}{q^{1-\alpha}(\mathbf{x}) + D\,\epsilon\,\Delta\, q^{1-\alpha}(\mathbf{x}) - \alpha\, D\,\epsilon\, q^{1-\alpha}(\mathbf{x})\, \frac{\Delta\, q(\mathbf{x})}{q(\mathbf{x})} + \mathcal{O}(\epsilon^2)} \\ &= \left(f(\mathbf{x}) + \frac{D\,\epsilon\,\Delta\left(q^{1-\alpha}(\mathbf{x})\, f(\mathbf{x})\right)}{q^{1-\alpha}(\mathbf{x})} - \alpha\, D\,\epsilon\, f(\mathbf{x})\, \frac{\Delta\, q(\mathbf{x})}{q(\mathbf{x})} + \mathcal{O}(\epsilon^2)\right) \\ &\qquad \times \left(1 - \frac{D\,\epsilon\,\Delta\, q^{1-\alpha}(\mathbf{x})}{q^{1-\alpha}(\mathbf{x})} + \alpha\, D\,\epsilon\, \frac{\Delta\, q(\mathbf{x})}{q(\mathbf{x})} + \mathcal{O}(\epsilon^2)\right) \\ &= f(\mathbf{x}) + \frac{D\,\epsilon\,\Delta\left(q^{1-\alpha}(\mathbf{x})\, f(\mathbf{x})\right)}{q^{1-\alpha}(\mathbf{x})} - \frac{D\,\epsilon\,\Delta\, q^{1-\alpha}(\mathbf{x})}{q^{1-\alpha}(\mathbf{x})}\, f(\mathbf{x}) + \mathcal{O}(\epsilon^2). \end{aligned} \tag{S25}$$

It is immediately obvious that the infinitesimal generator of this Markov chain $\Delta_\epsilon^{(\alpha)} f(\mathbf{x}) = (M_b^{(\alpha)}\, f(\mathbf{x}) - f(\mathbf{x}))/\epsilon$ satisfies

$$\lim_{\epsilon\to 0} \Delta_\epsilon^{(\alpha)}\, f(\mathbf{x}) = D\, \frac{\Delta(f(\mathbf{x})\, q^{1-\alpha}(\mathbf{x}))}{q^{1-\alpha}(\mathbf{x})} - D\, \frac{\Delta(q^{1-\alpha}(\mathbf{x}))}{q^{1-\alpha}(\mathbf{x})} f(\mathbf{x}), \tag{S26}$$

which is equivalent to Theorem 2 in [6], modulo the overall factor of $D$. In other words, given a probability density $q(\mathbf{x})$ and a kernel $K_\epsilon$, we can construct a Markov transition operator $M_b^{(\alpha)}\, f(\mathbf{x})$ whose infinitesimal generator in the continuum limit yields a one-parameter family of generalized Laplace operators that depend on $\alpha$. In particular, we see that $\lim_{\epsilon\to 0} \Delta_\epsilon^{(1)} f(\mathbf{x}) = D\,\Delta\, f(\mathbf{x})$, i.e. we recover the Laplace-Beltrami operator on $\mathcal{M}$ with no influence from the sampling density $q(\mathbf{x})$. Furthermore, for any $t > 0$, the heat kernel $e^{t\,D\,\Delta}$ on $\mathbb{R}^d$ is given by

$$\lim_{\epsilon\to 0} \left(M_b^{(1)}\right)^{t/\epsilon} = e^{t\,D\,\Delta} \tag{S27}$$

(see Proposition 3 in [5] with $D = 1$). This construction has several advantages. Firstly, we did not have to make any explicit assumptions about the intrinsic dimensionality of $\mathcal{M}$. Secondly, the construction is theoretically insensitive to non-uniform sampling (empirically, however, extreme inhomogeneity can degrade performance). Again with some abuse of notation, we can see that the discrete operators defined in the previous section are Monte Carlo estimates for the integral operators defined in this section

$$\begin{aligned} M_b^{(\alpha)} f(\mathbf{x}) &= \int_{\mathbb{R}^d} \frac{K_\epsilon^{(\alpha)}(\mathbf{x},\mathbf{y})}{d_\epsilon^{(\alpha)}(\mathbf{x})} f(\mathbf{y})\, q(\mathbf{y})\, d\mathbf{y} \approx \frac{\frac{1}{N}\sum_{j=1}^N K_\epsilon^{(\alpha)}(\mathbf{x}_i,\mathbf{x}_j)\, f(\mathbf{x}_j)}{\frac{1}{N}\sum_{m=1}^N K_\epsilon^{(\alpha)}(\mathbf{x}_i,\mathbf{x}_m)} \\ &\implies \mathbf{M}_b^{(\alpha)}\,\mathbf{f} = \left(\mathbf{D}_\epsilon^{(1)}\right)^{-1} \mathbf{K}_\epsilon^{(1)}\,\mathbf{f}. \end{aligned} \tag{S28}$$

We therefore arrive at a discrete approximation to the Laplace-Beltrami operator defined on $\hat{\mathcal{M}}$ given by the matrix

$$\boldsymbol{\Delta} = \left( \left(\mathbf{D}_\epsilon^{(1)}\right)^{-1} \mathbf{K}_\epsilon^{(1)} - \mathbf{I} \right) / \,(D\,\epsilon) \tag{S29}$$

An important point here is that, ultimately, the Laplace-Beltrami operator is completely defined by the geometry of the manifold $\mathcal{M}$ independent of any dynamical process that may unfold on $\mathcal{M}$. Without loss of generality, we will typically set $D = 1$, absorbing its effects into $\epsilon$, in order to avoid any confusion with the diffusion coefficient of the developmental processes defined over $\mathcal{M}$ that we seek to model.

##### S1.4.2 Schematic derivation of discrete Laplacians

In this section, we provide a schematic derivation of the general form a discrete Laplace operator using geometric reasoning. This structure naturally yields a volume element that we can associate to points in our discrete manifold $\hat{\mathcal{M}}$ to define the size of their local neighborhoods. In the next section, we will leverage this form to develop a decomposition of $\boldsymbol{\Delta}$ that improves numerical stability and is critical to our method for computing the interpolated potential $U_I$ (see Appendix S1.5). We will also demonstrate how the Laplace operator can be used to compute the squared gradient of a function $||\nabla f||^2$. A more complete and rigorous treatment can be found in [15]. Consider the integral of the continuous Lapacian of a function $f(\mathbf{x})$ over a region $\mathcal{V}_i$ centered on $\mathbf{x}_i \in \mathcal{M}$

$$\int_{\mathcal{V}_i} dV(\mathbf{y})\,\Delta f(\mathbf{y}) = \int_{\mathcal{V}_i} dV(\mathbf{y})\,\nabla\cdot\nabla f(\mathbf{y}) = \oint_{\partial\mathcal{V}_i} dA(\mathbf{y})\,\hat{\mathbf{n}}\cdot\nabla f(\mathbf{y}). \tag{S30}$$

where we have made use of the divergence theorem in the final equality. $\partial\mathcal{V}_i$ denotes the boundary of $\mathcal{V}_i$, $dA$ its area element, and $\hat{\mathbf{n}}$ its outward facing normal. We can approximate the left and right hand sides of Eq. (S30) with

$$V_i\,\Delta f(\mathbf{x}_i) \approx \sum_{j\in\mathcal{N}(\mathbf{x}_i)} A_{ij}\,\frac{f(\mathbf{x}_j)-f(\mathbf{x}_i)}{\ell_{ij}} = \sum_{j\in\mathcal{N}(\mathbf{x}_i)} w_{ij}\,\left(f(\mathbf{x}_j)-f(\mathbf{x}_i)\right), \tag{S31}$$

where $\mathcal{N}(\mathbf{x}_i)$ denotes the neighborhood of $\mathbf{x}_i$, $\ell_{ij} = ||\mathbf{x}_j - \mathbf{x}_i||$, $A_{ij}$ is the portion of $\partial\mathcal{V}_i$ that we associate to $\mathbf{x}_j$ and we use a simple finite difference approximation for $\hat{\mathbf{n}}\cdot\nabla f(\mathbf{y}) \approx (f(\mathbf{x}_j) - f(\mathbf{x}_i))/\ell_{ij}$. Written out in terms of the symmetric coefficients $w_{ij} = w_{ji}$, Eq. (S31) is the general form for a graph Laplacian. Note that we have implicitly assumed that the points $\{\mathbf{x}_i\}$ and the edges between them form a manifold simplicial complex over, e.g. a Delaunay graph constructed from a Voronoi decomposition of $\mathcal{M}$ with edges connecting points that seed adjacent Voronoi cells. This is not strictly true for our diffusion map Laplacian, since our affinity matrix $\mathbf{K}_\epsilon$ (Eq. (S9)) is fully connected. For our purposes this is a technicality that does not meaningfully change any subsequent interpretation. If we define the matrix coefficients of the weak Laplace operator

$$L(\mathbf{x}_i, \mathbf{x}_j) = \begin{cases} w_{ij} & j \in \mathcal{N}(\mathbf{x}_i) \\ -\sum_{k \in \mathcal{N}(\mathbf{x}_i)} w_{ik} & i = j \\ 0 & \text{else} \end{cases} \tag{S32}$$

and its associated diagonal mass matrix

$$M(\mathbf{x}_i, \mathbf{x}_j) = V(\mathbf{x}_i)\, \delta_{ij}, \tag{S33}$$

it is easy to show by direct computation that

$$\Delta f(\mathbf{x}_i) = \frac{1}{V(\mathbf{x}_i)} \sum_{j \in \mathcal{N}(\mathbf{x}_i)} w_{ij} \, \left( f(\mathbf{x}_j) - f(\mathbf{x}_i) \right) \implies \mathbf{\Delta f} = \mathbf{M}^{-1}\, \mathbf{L}\, \mathbf{f}. \tag{S34}$$

Notice that the integral of the Laplacian over the entire manifold

$$\int_{\mathcal{M}} dV(\mathbf{y})\, \Delta f(\mathbf{y}) = \oint_{\partial\mathcal{M}} dA(\mathbf{y})\, \hat{\mathbf{n}} \cdot \nabla f(\mathbf{y}) \approx \sum_i \sum_j L(\mathbf{x}_i, \mathbf{x}_j) f(\mathbf{x}_j) = 0 \tag{S35}$$

since $\sum_i L(\mathbf{x}_i, \mathbf{x}_j) = 0$. This Laplacian construction therefore naturally induces Neumann boundary conditions ($\hat{\mathbf{n}} \cdot \nabla f(\mathbf{x}) \mid_{\partial\mathcal{M}} = 0$) on manifolds $\mathcal{M}$ with non-empty boundaries. Now consider the integral

$$\begin{aligned} \int_{\mathcal{V}_i} dV(\mathbf{y})\, ||\nabla f(\mathbf{y})||^2 &= \int_{\mathcal{V}_i} dV(\mathbf{y})\, \nabla f(\mathbf{y}) \cdot \nabla f(\mathbf{y}) \\ &= \int_{\mathcal{V}_i} dV(\mathbf{y}) \nabla \cdot (f(\mathbf{y})\, \nabla f(\mathbf{y})) - \int_{\mathcal{V}_i} dV(\mathbf{y}) f(\mathbf{y})\, \Delta f(\mathbf{y}) \\ &= \oint_{\partial\mathcal{V}_i} dA(\mathbf{y})\, \hat{\mathbf{n}} \cdot (f(\mathbf{y})\, \nabla f(\mathbf{y})) - \int_{\mathcal{V}_i} dV(\mathbf{y}) f(\mathbf{y})\, \Delta f(\mathbf{y}). \end{aligned} \tag{S36}$$

where we have once again made use of the divergence theorem over the small region $\mathcal{V}_i$. We can leverage a similar approximation scheme as before

$$\begin{aligned} \int_{\mathcal{V}_i} dV(\mathbf{y})\, ||\nabla f(\mathbf{y})||^2 &\approx \sum_{j \in \mathcal{N}(\mathbf{x}_i)} A_{ij} \frac{f(\mathbf{x}_j) + f(\mathbf{x}_i)}{2} \frac{f(\mathbf{x}_j) - f(\mathbf{x}_i)}{\ell_{ij}} - V(\mathbf{x}_i) f(\mathbf{x}_i)\, \Delta f(\mathbf{x}_i) \\ &= \frac{1}{2} \sum_{j \in \mathcal{N}(\mathbf{x}_i)} w_{ij} \, \left( f(\mathbf{x}_j)^2 - f(\mathbf{x}_i)^2 \right) - \sum_{j \in \mathcal{N}(\mathbf{x}_i)} w_{ij} \, f(\mathbf{x}_i) \, \left( f(\mathbf{x}_j) - f(\mathbf{x}_i) \right) \\ &= \frac{1}{2} \sum_{j \in \mathcal{N}(\mathbf{x}_i)} w_{ij} \, \left( f(\mathbf{x}_j) - f(\mathbf{x}_i) \right)^2. \end{aligned} \tag{S37}$$

Integrating over the entire $\mathcal{M}$ yields the famous Dirichlet energy $\mathcal{E}_D$ [15] and direct computation shows that

$$\mathcal{E}_D = \frac{1}{2} \int_{\mathcal{M}} dV(\mathbf{y})\, ||\nabla f(\mathbf{y})||^2 \approx \frac{1}{4} \sum_i \sum_{j \in \mathcal{N}(\mathbf{x}_i)} w_{ij} \, \left( f(\mathbf{x}_j) - f(\mathbf{x}_i) \right)^2 = -\frac{1}{2} \mathbf{f}^T \mathbf{L} \mathbf{f}. \tag{S38}$$

The minus sign reflects our convention that $\boldsymbol{\Delta}$ is negative semi-definite. Finally, we note that

$$\begin{aligned} ||\nabla f(\mathbf{x}_i)||^2 &= \frac{1}{2\,V(\mathbf{x}_i)} \sum_{j \in \mathcal{N}(\mathbf{x}_i)} w_{ij}\,\left(f(\mathbf{x}_j) - f(\mathbf{x}_i)\right)^2 \\ \implies ||\boldsymbol{\nabla}\mathbf{f}||^2 &= \frac{1}{2}\mathbf{M}^{-1}\,\mathbf{L}\,(\mathbf{f}\odot\mathbf{f}) - \mathbf{M}^{-1}\,(\mathbf{f}\odot(\mathbf{L}\,\mathbf{f}))\,. \end{aligned} \tag{S39}$$

where $\odot$ denotes element-wise multiplication of vectors. We utilize this fact later on in our definition of the discrete dynamical transition operator (see Appendix S1.6).

##### S1.4.3 Numerical implementation of the diffusion map Laplacian

We now synthesize the results of the last two sections to reformulate our discrete Laplace operator in a way that improved numerical stability. Starting from Eq. (S29), we see

$$\boldsymbol{\Delta} = \left(\left(\mathbf{D}_\epsilon^{(1)}\right)^{-1}\mathbf{K}_\epsilon^{(1)} - \mathbf{I}\right)\,/\,(D\,\epsilon) = \left(\mathbf{D}_\epsilon^{(1)}\right)^{-1}\,\left(\mathbf{K}_\epsilon^{(1)} - \mathbf{D}_\epsilon^{(1)}\right)\,/\,(D\,\epsilon) = \mathbf{M}^{-1}\mathbf{L}, \tag{S40}$$

where the components of the diagonal mass matrix are given by

$$M(\mathbf{x}_i, \mathbf{x}_j) = \sqrt{D\,\epsilon}\,d_\epsilon^{(1)}(\mathbf{x}_i)\,\delta_{ij} := \sqrt{D\,\epsilon}\,D_\epsilon^{(1)}(\mathbf{x}_i, \mathbf{x}_j). \tag{S41}$$

and the components of the weak Laplace operator are given by

$$L(\mathbf{x}_i, \mathbf{x}_j) = \left(K_\epsilon^{(1)}(\mathbf{x}_i, \mathbf{x}_j) - d_\epsilon^{(1)}(\mathbf{x}_i)\,\delta_{ij}\right)/\sqrt{D\,\epsilon} = \left(K_\epsilon^{(1)}(\mathbf{x}_i, \mathbf{x}_j) - D_\epsilon^{(1)}(\mathbf{x}_i, \mathbf{x}_j)\right)/\sqrt{D\,\epsilon}. \tag{S42}$$

We observe empirically that splitting the factor of $1/(D\,\epsilon)$ between the two matrices produces more numerically stable results in certain computations. Notice that $\mathbf{L}$ is a symmetric, negative semi-definite operator with all $L(\mathbf{x}_i, \mathbf{x}_j) \geq 0$ for $i \neq j$. These properties are sufficient to guarantee that $\mathbf{L}$ satisfies a discrete maximum principle [13], i.e., solutions to $\mathbf{L}\,\mathbf{f} = \mathbf{0}$ have no local extrema away from boundaries or constrained points. If $\mathcal{M}$ has a non-empty boundary, and no Dirichlet boundary conditions are explicitly set, then we should interpret $\boldsymbol{\Delta}$ as inducing Neumann boundary conditions [5]. We also see that this construction provides a natural notion of a discrete volume element $V_L(\mathbf{x}_i) := \sqrt{D\,\epsilon}\,d_\epsilon^{(1)}(\mathbf{x})$ from the components of $\mathbf{M}$ (the subscript 'L' stands for 'Laplacian' - see also Eq. (S33)). Without loss of generality, we typically set $D = 1$ to avoid any confusion with the diffusion coefficient of the developmental dynamics defined over $\hat{\mathcal{M}}$ that we seek to model.

#### S1.5 Solving for the interpolated potential $U_I$

In order to model non-steady state developmental dynamics, we build our dynamical potential from two components: $U_D = U_B + U_I$. The base potential $U_B$ characterizes how probability diffuses from point to point in the absence of directed motion and confines dynamics to the manifold's interior. It is simply $U_B = -\log p_B$, where $p_B$ is the kernel density estimate for the points that define our discrete manifold $\hat{\mathcal{M}} = \{\mathbf{x}\}_{i=1}^N$. The interpolated potential $U_I$ produces directed motion by changing the topography of the landscape. A candidate function $U_I$ should be smooth and well-behaved, but sufficiently expressive to let us selectively destabilize regions of our dynamical manifold (i.e. near the measured initial conditions) and stabilize others (i.e. the terminal conditions). Let $\{\mathbf{x}_i^*\}_{i=1}^m$ refer to a sparse subset of $\hat{\mathcal{M}}$. In practice, these are usually the set of $m$ density maxima and their associated base potential saddles. Some of the $\mathbf{x}_i^*$ are connected by paths $\mathcal{W}_B(\mathbf{x}_i^*, \mathbf{x}_j^*)$. Specifically, $\mathcal{W}_B(\mathbf{x}_i^*, \mathbf{x}_j^*)$ is an ordered list of points $\{\mathbf{x}_1 = \mathbf{x}_i^*, \mathbf{x}_2, \ldots, \mathbf{x}_{|\mathcal{W}_B|} = \mathbf{x}_j^*\}$ where $|\mathcal{W}_B|$ is the number of points in the path. These are usually the most probable paths between $\mathbf{x}_i^*$ and $\mathbf{x}_j^*$ defined relative to the $U_B$ and computed using the base potential transition operator. The function we seek can be found as the minimizer of a smoothness functional, namely the squared Laplacian energy:

$$U_I = \underset{U}{\text{argmin}}\, \mathcal{E}_{\Delta^2} = \underset{U}{\text{argmin}}\, \frac{1}{2} \int ||\Delta\, U(\mathbf{x})||^2 d\mathbf{x}, \tag{S43}$$

subject to:

$$U_I(\mathbf{x}_i^*) = h_i^*, \tag{S44}$$

$$U_I(\mathbf{x}_k) = \lambda_k\, h_j^* + (1 - \lambda_k)\, h_i^*, \quad \text{for } \mathbf{x}_k \in \mathcal{W}_B(\mathbf{x}_i^*, \mathbf{x}_j^*). \tag{S45}$$

The $\lambda_k \in [0, 1]$ are just the fractional position of $\mathbf{x}_k$ along $\mathcal{W}_B(\mathbf{x}_i^*, \mathbf{x}_j^*)$. Explicitly,

$$\lambda_k = \sum_{i=1}^{k} ||\mathbf{x}_i - \mathbf{x}_{i-1}|| \,/ \sum_{i=1}^{|\mathcal{W}_B|} ||\mathbf{x}_i - \mathbf{x}_{i-1}||. \tag{S46}$$

In words, we adjust the "heights" of the fixed points $h_i^*$, constrain the potential along the most probable paths to linearly interpolate between them, and smoothly interpolate everywhere else by minimizing Eq. (S43). This energy has been widely used in graphics, computational geometry, and image processing to model deformations [16], denoise densely sampled data [17], and interpolate sparse, scattered measurements [18]. It is analogous to a high-dimensional version of the linearized thin-plate spline energy and produces a smooth function by penalizing sharp changes in curvature [18].

Minimizing the discretized version of $\mathcal{E}_{\Delta^2}$ on $\hat{\mathcal{M}}$ turns out to be a fairly straightforward quadratic minimization problem. Using our discretation of the Laplacian $\mathbf{\Delta} = \mathbf{M}^{-1}\,\mathbf{L}$, we have

$$\begin{aligned} \mathcal{E}_{\Delta^2} &= \frac{1}{2} \int ||\Delta\, U(\mathbf{x})||^2 d\mathbf{x} \approx \frac{1}{2} \sum_i (\Delta U(\mathbf{x}_i))^2\, V(\mathbf{x}_i) \\ &= \frac{1}{2} \sum_i (\sum_j \sum_k M^{-1}(\mathbf{x}_i, \mathbf{x}_j)\, L(\mathbf{x}_j, \mathbf{x}_k)\, U(\mathbf{x}_k))^2\, V(\mathbf{x}_i) = \frac{1}{2} \sum_i (\sum_k L(\mathbf{x}_i, \mathbf{x}_k)\, U(\mathbf{x}_k)\, / \, V(\mathbf{x}_i))^2\, V(\mathbf{x}_i) \\ &= \frac{1}{2} \sum_j \sum_k U(\mathbf{x}_j) \left( \sum_i L(\mathbf{x}_j, \mathbf{x}_i)\, L(\mathbf{x}_i, \mathbf{x}_k)\, / \, V(\mathbf{x}_i) \right) U(\mathbf{x}_k) = \frac{1}{2}\, \mathbf{U}^T\, \mathbf{L}\, \mathbf{M}^{-1}\, \mathbf{L}\, \mathbf{U} = \frac{1}{2}\, \mathbf{U}^T\, \mathbf{Q} \mathbf{U} \end{aligned} \tag{S47}$$

where we have identified the diagonal elements of the mass matrix with the integration volumes $M(\mathbf{x}_i, \mathbf{x}_i) = V(\mathbf{x}_i)$ (see Eqs. (S33) and (S41)). Notice that the matrix $\mathbf{Q}$ is a symmetric, positive semi-definite matrix thanks to the properties of our discrete Laplacian. There are many methods and black-box solvers to minimize quadratic objective functions such as Eq. (S47) subject to equality constraints (Eq. (S44) and Eq. (S45)). See, for instance, MATLAB's `quadprog`. Unlike $\mathbf{L}$, the matrix $\mathbf{Q}$ will generally not exhibit a discrete maximum principle and points that are not elements of any path $\mathcal{W}_B$ may have values of $U_I$ that fall outside the range set by the equality constraints. Empirically, we observe that this can lead to isolated outliers when the equality constraints vary rapidly along a particular $\mathcal{W}_B$. We include functionality to remove outliers outside of a user-specified window by setting the value of $U_I$ at these locations to the average value of their $k$-nearest neighbors ($k = 10$ by default). We also note that we include a number of optional constraints, including the ability to fix the value of a particular $h_i^*$ and to fix the sum of all $h_i^*$. All results presented in this paper were found fixing the sum $\sum_i h_i^* = 0$, which breaks any degeneracy in the fits resulting from changing the potential by a constant shift.

#### S1.6 Discretization of the transition operator

In this section, we explicitly show how to discretize the transition operator shown in Eq. (S4) for a given $U_D(\mathbf{x})$. We begin with a derivation of a transition operator acting on probability vectors $P(\mathbf{x}_i, t_k)$ using fairly standard geometric finite-difference methods. Next, we define an alternative operator that acts on probability *densities* and show that it converges to the generator of the Fokker-Planck operator in the continuum limit using a generalization of the methods in Appendix S1.4.1. Finally, we show that these two operators are numerically equivalent so long as we choose appropriate volume elements for the points in $\hat{\mathcal{M}}$. These results synthesize all of the developments of the previous section and highlight and underlying connection between ours and existing methods for modeling discrete drift-diffusion processes.

The probability of finding a cell in the neighborhood $\mathcal{V}_i$ of $\mathbf{x}_i$ at time $t$ is just

$$P(\mathbf{x}_i, t) = \int_{\mathcal{V}_i} d\mathbf{y}\, p(\mathbf{y}, t) \approx p(\mathbf{x}_i, t)\, V(\mathbf{x}_i), \tag{S48}$$

where $\sum_{i=1}^{N} P(\mathbf{x}_i, t) = 1$ and $V(\mathbf{x}_i)$ is the volume of $\mathcal{V}_i$. We then discretize Eq. (S4) in the following way:

$$P(\mathbf{x}_i, t_{k+1}) = \sum_{j=1}^{N} T_\epsilon(\mathbf{x}_i, \mathbf{x}_j)\, P(\mathbf{x}_j, t_k) \propto V(\mathbf{x}_i) \sum_{j=1}^{N} \exp\left[-\frac{||\mathbf{x}_i - \mathbf{x}_j - \epsilon\, \mathbf{v}(\mathbf{x}_j, t)||^2}{4D\epsilon}\right] P(\mathbf{x}_j, t_k), \tag{S49}$$

where the transition matrix is normalized as $\sum_{i=1}^{N} T_\epsilon(\mathbf{x}_i, \mathbf{x}_j) = 1$ and $t_k = k\,\epsilon$ is the time at the $k$th step in the discrete Markov process. Explicitly substituting the gradient-like drift $\mathbf{v} = \mathbf{g}^{-1}\nabla U$ and isolating the transition operator yields

$$\begin{aligned} T_\epsilon(\mathbf{x}_i, \mathbf{x}_j) &= \frac{V(\mathbf{x}_i)}{\mathcal{Z}_j} \exp\left[-\frac{||\mathbf{x}_i - \mathbf{x}_j - \epsilon\, \mathbf{v}(\mathbf{x}_j, t)||^2}{4\, D\, \epsilon}\right] \\ &= \frac{V(\mathbf{x}_i)}{\mathcal{Z}_j} \exp\left[-\frac{||\mathbf{x}_i - \mathbf{x}_j||^2}{4\, D\, \epsilon} - \frac{(\mathbf{x}_i - \mathbf{x}_j)\cdot \mathbf{g}^{-1}(\mathbf{x}_j, t)\nabla\, U(\mathbf{x}_j, t)}{2\, D} - \frac{\epsilon\, ||\mathbf{g}^{-1}(\mathbf{x}_j, t)\nabla\, U(\mathbf{x}_j, t)||^2}{4\, D}\right], \end{aligned} \tag{S50}$$

where $\mathcal{Z}_j$ is a per-column normalization constant. To proceed we make a few simplifications. First, we focus on time-independent landscapes, i.e $U(\mathbf{x}, t) = U(\mathbf{x})$, $\mathbf{g}(\mathbf{x}, t) = \mathbf{g}(\mathbf{x})$. Second, we assume that the metric tensor is conformal, i.e. $\mathbf{g}(\mathbf{x}) = g(\mathbf{x})\, \mathbf{I}$, where $\mathbf{I}$ is the identity tensor and $g$ is a positive scalar function. Given these assumptions, the transition operator becomes

$$T_\epsilon(\mathbf{x}_i, \mathbf{x}_j) = \frac{V(\mathbf{x}_i)}{\mathcal{Z}_j} \exp\left[-\frac{||\mathbf{x}_i - \mathbf{x}_j||^2}{4\, D\, \epsilon} - \frac{(\mathbf{x}_i - \mathbf{x}_j)\cdot \nabla\, U(\mathbf{x}_j)}{2\, D\, g(\mathbf{x}_j)} - \frac{\epsilon\, ||\nabla\, U(\mathbf{x}_j)||^2}{4\, D\, g^2(\mathbf{x}_j)}\right]. \tag{S51}$$

We choose a simple finite-difference approximation for directional derivative in the exponent

$$\frac{(\mathbf{x}_i - \mathbf{x}_j)\cdot \nabla\, U(\mathbf{x}_j)}{2\, D\, g(\mathbf{x}_j)} \approx \frac{U(\mathbf{x}_i) - U(\mathbf{x}_j)}{D\, (g(\mathbf{x}_i) + g(\mathbf{x}_j))}. \tag{S52}$$

Notice we have replaced the metric with its average value on the edge connecting $\mathbf{x}_i$ and $\mathbf{x}_j$. Our transition operator is fully connected and this approximation will be poor for distant pairs of points. However, for sufficiently small $\epsilon$, the factor $\exp[-||\mathbf{x}_i - \mathbf{x}_j||^2/4\, D\, \epsilon]$ strongly suppresses such contributions and we do not notice any issues numerically. When the metric is uniform over all space this reduces to $(U(\mathbf{x}_i) - U(\mathbf{x}_j))\,/\,2\,D\,g$. The term $\propto ||\nabla U(\mathbf{x}_j)||^2$ can be computed using Eq. (S39). Substituting these approximations yields

$$T_\epsilon(\mathbf{x}_i, \mathbf{x}_j) = \frac{V(\mathbf{x}_i)}{\mathcal{Z}_j} \exp\left[ -\frac{||\mathbf{x}_i - \mathbf{x}_j||^2}{4\,D\,\epsilon} - \frac{U(\mathbf{x}_i) - U(\mathbf{x}_j)}{D\,(g(\mathbf{x}_i) + g(\mathbf{x}_j))} - \frac{\epsilon\,||\nabla\,U(\mathbf{x}_j)||^2}{4\,D\,g^2(\mathbf{x}_j)} \right]. \tag{S53}$$

The condition $\sum_{i=1}^N T_\epsilon(\mathbf{x}_i, \mathbf{x}_j) = 1$ tells us that

$$\mathcal{Z}_j = \sum_{i=1}^N V(\mathbf{x}_i) \exp\left[ -\frac{||\mathbf{x}_i - \mathbf{x}_j||^2}{4\,D\,\epsilon} - \frac{U(\mathbf{x}_i) - U(\mathbf{x}_j)}{D\,(g(\mathbf{x}_i) + g(\mathbf{x}_j))} - \frac{\epsilon\,||\nabla\,U(\mathbf{x}_j)||^2}{4\,D\,g^2(\mathbf{x}_j)} \right]. \tag{S54}$$

The simplified components of the transition operator can therefore be written as

$$T_\epsilon(\mathbf{x}_i, \mathbf{x}_j) = \frac{V(\mathbf{x}_i) \exp\left[ -\frac{||\mathbf{x}_i - \mathbf{x}_j||^2}{4\,D\,\epsilon} - \frac{U(\mathbf{x}_i) - U(\mathbf{x}_j)}{D\,(g(\mathbf{x}_i) + g(\mathbf{x}_j))} \right]}{\sum_n V(\mathbf{x}_n) \exp\left[ -\frac{||\mathbf{x}_n - \mathbf{x}_j||^2}{4\,D\,\epsilon} - \frac{U(\mathbf{x}_n) - U(\mathbf{x}_j)}{D\,(g(\mathbf{x}_n) + g(\mathbf{x}_j))} \right]}. \tag{S55}$$

Notice that the terms $\propto ||U(\mathbf{x}_j)||^2$ vanish after normalization since they depend only on $\mathbf{x}_j$. When the metric is uniform over all space, we could technically cancel the terms $\left[U(\mathbf{x}_j)/(D\,g)\right]$ in the exponent as well. In practice, however, we compute them regardless since keeping the difference $U(\mathbf{x}_i) - U(\mathbf{x}_j)$ in the equation improves numerical stability.

###### S1.6.1 Derivation of the Fokker-Planck transition operator from diffusion processes

Here, we will utilize the framework for modeling drift-diffusion processes developed in Appendix S1.4.1 to define an alternative transition operator that acts on probability densities rather than probabilities. We will also provide a simplified proof that this operator converges to the generator of the Fokker-Planck equation in the continuum limit ($N \to \infty$, $\epsilon \to 0$). The following derivation considers a family of drift diffusion processes that depend on a scalar parameter $\alpha$. As before, the specific process corresponding to $\alpha = 1$ is most relevant to us, but we include a general treatment for completeness. Once again, consider a probability density $q(\mathbf{x})$ defined over $\mathbb{R}^d$. We emphasize that $q(\mathbf{x})$ is just an arbitrary distribution for sampling points and is *not* necessarily related to our dynamical probability density $p(\mathbf{x}, t)$ for a given experimental condition. We define the following kernel

$$\tilde{K}_\epsilon(\mathbf{x}, \mathbf{y}) = \exp\left[ -\frac{||\mathbf{x} - \mathbf{y} - \epsilon\,\mathbf{v}(\mathbf{y})||^2}{4\,D\,\epsilon} \right] = \exp\left[ -\frac{||\mathbf{x} - \mathbf{y}||^2}{4\,D\,\epsilon} + \frac{(\mathbf{x} - \mathbf{y}) \cdot \mathbf{v}(\mathbf{y})}{2\,D} - \frac{\epsilon\,||\mathbf{v}(\mathbf{y})||^2}{4\,D} \right] \tag{S56}$$

for a time-independent vector field $\mathbf{v}(\mathbf{x})$. We would like to determine the leading order behavior in $\epsilon$ of the following integral operator

$$\int_{\mathbb{R}^d} \tilde{K}_\epsilon(\mathbf{x}, \mathbf{y})\, f(\mathbf{y})\, d\mathbf{y} = \int_{\mathbb{R}^d} \exp\left[ -\frac{||\mathbf{x} - (\mathbf{y} + \epsilon\,\mathbf{v}(\mathbf{y}))||^2}{4\,D\,\epsilon} \right] f(\mathbf{y})\, d\mathbf{y}. \tag{S57}$$

We use the change of variables $\mathbf{z}(\mathbf{y}) = \mathbf{y} + \epsilon\,\mathbf{v}(\mathbf{y})$. The Jacobian of this transformation is $\mathbf{J} = \nabla \mathbf{z}(\mathbf{y}) = \mathbf{I} + \epsilon\,\nabla \mathbf{v}(\mathbf{y})$. The volume form of our integral therefore changes according to

$$\begin{aligned} d\mathbf{y} &= |\det \mathbf{J}\,|^{-1}\, d\mathbf{z} = |\det\left[\mathbf{I} + \epsilon\,\nabla \mathbf{v}(\mathbf{y})\right]\,|^{-1}\, d\mathbf{z} \\ &= \left(1 - \epsilon\,\mathrm{Tr}\left[\nabla \mathbf{v}(\mathbf{z})\right] + \mathcal{O}(\epsilon^2)\right)\, d\mathbf{z} = \left(1 - \epsilon\,\nabla \cdot \mathbf{v}(\mathbf{z}) + \mathcal{O}(\epsilon^2)\right)\, d\mathbf{z}. \end{aligned} \tag{S58}$$

For small $\epsilon$, we approximate

$$f(\mathbf{y}) = f(\mathbf{z} - \epsilon\,\mathbf{v}(\mathbf{y})) \approx f(\mathbf{z} - \epsilon\,\mathbf{v}(\mathbf{z})) = f(\mathbf{z}) - \epsilon\nabla f(\mathbf{z})\cdot\mathbf{v}(\mathbf{z}) + \mathcal{O}(\epsilon^2) \tag{S59}$$

Combining these approximations and making use of Eq. (S21), we find

$$\begin{aligned}\int_{\mathbb{R}^d} \tilde{K}_\epsilon(\mathbf{x},\mathbf{y})\, f(\mathbf{y})\, d\mathbf{y} &= \int_{\mathbb{R}^d} K_\epsilon(\mathbf{x},\mathbf{z})\, \left(f(\mathbf{z}) - \epsilon\nabla f(\mathbf{z})\cdot\mathbf{v}(\mathbf{z}) + \mathcal{O}(\epsilon^2)\right)\, \left(1 - \epsilon\,\nabla\cdot\mathbf{v}(\mathbf{z}) + \mathcal{O}(\epsilon^2)\right)\, d\mathbf{z} \\ &= \int_{\mathbb{R}^d} K_\epsilon(\mathbf{x},\mathbf{z})\, \left(f(\mathbf{z}) - \epsilon\left(\nabla f(\mathbf{z})\cdot\mathbf{v}(\mathbf{z}) + f(\mathbf{z})\,\nabla\cdot\mathbf{v}(\mathbf{z})\right) + \mathcal{O}(\epsilon^2)\right)\, d\mathbf{z} \\ &= \int_{\mathbb{R}^d} K_\epsilon(\mathbf{x},\mathbf{z})\, \left(f(\mathbf{z}) - \epsilon\,\nabla\cdot\left(f(\mathbf{z})\,\mathbf{v}(\mathbf{z})\right) + \mathcal{O}(\epsilon^2)\right)\, d\mathbf{z} \\ &= (4\,\pi\,D\,\epsilon)^{d/2}\, \left(f(\mathbf{x}) - \epsilon\,\nabla\cdot\left(f(\mathbf{x})\,\mathbf{v}(\mathbf{x})\right) + D\,\epsilon\,\Delta f(\mathbf{x}) + \mathcal{O}(\epsilon^2)\right).\end{aligned} \tag{S60}$$

Using the same conventions as the previous sections, we define the integral operator

$$\begin{aligned}&\tilde{K}^{(\alpha)}_\epsilon\, f(\mathbf{x}) := \int_{\mathbb{R}^d} \tilde{K}^{(\alpha)}_\epsilon(\mathbf{x},\mathbf{y})\, f(\mathbf{y})\, q(\mathbf{y})\, d\mathbf{y} = \int_{\mathbb{R}^d} \frac{\tilde{K}_\epsilon(\mathbf{x},\mathbf{y})}{q^\alpha_\epsilon(\mathbf{x})\, q^\alpha_\epsilon(\mathbf{y})}\, f(\mathbf{y})\, q(\mathbf{y})\, d\mathbf{y} \\ &= \frac{1}{q^\alpha_\epsilon(\mathbf{x})} \int_{\mathbb{R}^d} \tilde{K}_\epsilon(\mathbf{x},\mathbf{y})\, q^{-\alpha}_\epsilon(\mathbf{y})\, f(\mathbf{y})\, q(\mathbf{y})\, d\mathbf{y} \\ &= \frac{(4\,\pi\,D\,\epsilon)^{-\alpha d/2}}{q^\alpha_\epsilon(\mathbf{x})} \int_{\mathbb{R}^d} \tilde{K}_\epsilon(\mathbf{x},\mathbf{y})\, q^{1-\alpha}(\mathbf{y})\, f(\mathbf{y})\, \left(1 - \alpha\,D\,\epsilon\frac{\Delta\,q(\mathbf{x})}{q(\mathbf{x})} + \mathcal{O}(\epsilon^2)\right)\, d\mathbf{y} \\ &= \frac{(4\,\pi\,D\,\epsilon)^{(1-\alpha)d/2}}{q^\alpha_\epsilon(\mathbf{x})} \times \\ &\left(q^{1-\alpha}(\mathbf{x})\, f(\mathbf{x}) - \epsilon\,\nabla\cdot\left(q^{1-\alpha}(\mathbf{x})\, f(\mathbf{x})\, \mathbf{v}(\mathbf{x})\right) + D\,\epsilon\,\Delta\left(q^{1-\alpha}(\mathbf{x})\, f(\mathbf{x})\right) - \alpha\,D\,\epsilon\, q^{1-\alpha}(\mathbf{x})\, f(\mathbf{x})\, \frac{\Delta\,q(\mathbf{x})}{q(\mathbf{x})} + \mathcal{O}(\epsilon^2)\right).\end{aligned} \tag{S61}$$

We can now construct an integral Markov transition operator

$$\begin{aligned}&\tilde{M}^{(\alpha)}_b\, f(\mathbf{x}) := \frac{\tilde{K}^{(\alpha)}_\epsilon\, f(\mathbf{x})}{d^{(\alpha)}_\epsilon(\mathbf{x})} = \int_{\mathbb{R}^d} \frac{\tilde{K}^{(\alpha)}_\epsilon(\mathbf{x},\mathbf{y})}{d^{(\alpha)}_\epsilon(\mathbf{x})}\, f(\mathbf{y})\, q(\mathbf{y})\, d\mathbf{y} \\ &= \frac{q^{1-\alpha}(\mathbf{x})\, f(\mathbf{x}) - \epsilon\,\nabla\cdot\left(q^{1-\alpha}(\mathbf{x})\, f(\mathbf{x})\, \mathbf{v}(\mathbf{x})\right) + D\,\epsilon\,\Delta\left(q^{1-\alpha}(\mathbf{x})\, f(\mathbf{x})\right) - \alpha\,D\,\epsilon\, q^{1-\alpha}(\mathbf{x})\, f(\mathbf{x})\, \frac{\Delta\,q(\mathbf{x})}{q(\mathbf{x})} + \mathcal{O}(\epsilon^2)}{q^{1-\alpha}(\mathbf{x}) + D\,\epsilon\,\Delta\, q^{1-\alpha}(\mathbf{x}) - \alpha\,D\,\epsilon\, q^{1-\alpha}(\mathbf{x})\, \frac{\Delta\,q(\mathbf{x})}{q(\mathbf{x})} + \mathcal{O}(\epsilon^2)} \\ &= \left(f(\mathbf{x}) - \epsilon\,\frac{\nabla\cdot\left(q^{1-\alpha}(\mathbf{x})\, f(\mathbf{x})\, \mathbf{v}(\mathbf{x})\right)}{q^{1-\alpha}(\mathbf{x})} + D\,\epsilon\,\frac{\Delta\left(q^{1-\alpha}(\mathbf{x})\, f(\mathbf{x})\right)}{q^{1-\alpha}(\mathbf{x})} - \alpha\,D\,\epsilon\, f(\mathbf{x})\,\frac{\Delta\,q(\mathbf{x})}{q(\mathbf{x})} + \mathcal{O}(\epsilon^2)\right) \\ &\qquad\times\left(1 - D\,\epsilon\,\frac{\Delta\,q^{1-\alpha}(\mathbf{x})}{q^{1-\alpha}(\mathbf{x})} + \alpha\,D\,\epsilon\,\frac{\Delta\,q(\mathbf{x})}{q(\mathbf{x})} + \mathcal{O}(\epsilon^2)\right) \\ &= f(\mathbf{x}) - \epsilon\,\frac{\nabla\cdot\left(q^{1-\alpha}(\mathbf{x})\, f(\mathbf{x})\, \mathbf{v}(\mathbf{x})\right)}{q^{1-\alpha}(\mathbf{x})} + D\,\epsilon\,\frac{\Delta\left(q^{1-\alpha}(\mathbf{x})\, f(\mathbf{x})\right)}{q^{1-\alpha}(\mathbf{x})} - D\,\epsilon\,\frac{\Delta\,q^{1-\alpha}(\mathbf{x})}{q^{1-\alpha}(\mathbf{x})}\, f(\mathbf{x}) + \mathcal{O}(\epsilon^2).\end{aligned} \tag{S62}$$

The infinitesimal generator of this Markov chain $\tilde{\Delta}^{(\alpha)}_\epsilon f(\mathbf{x}) = (\tilde{M}^{(\alpha)}_b\, f(\mathbf{x}) - f(\mathbf{x}))\,/\,\epsilon$ satisfies

$$\lim_{\epsilon\to 0} \tilde{\Delta}^{(\alpha)}_\epsilon\, f(\mathbf{x}) = -\frac{\nabla\cdot\left(q^{1-\alpha}(\mathbf{x})\, f(\mathbf{x})\, \mathbf{v}(\mathbf{x})\right)}{q^{1-\alpha}(\mathbf{x})} + D\,\frac{\Delta(f(\mathbf{x})\, q^{1-\alpha}(\mathbf{x}))}{q^{1-\alpha}(\mathbf{x})} - D\,\frac{\Delta(q^{1-\alpha}(\mathbf{x}))}{q^{1-\alpha}(\mathbf{x})} f(\mathbf{x}), \tag{S63}$$

which correctly reduces to Eq. (S26) when $\mathbf{v}(\mathbf{x}) = 0$. Notably, for $\alpha = 1$ we see that

$$\lim_{\epsilon\to 0} \tilde{\Delta}_\epsilon^{(1)} f(\mathbf{x}) = -\nabla \cdot (f(\mathbf{x})\,\mathbf{v}(\mathbf{x})) + D\,\Delta f(\mathbf{x}), \tag{S64}$$

which is precisely the generator of the Fokker-Planck equation (Eq. (S3)). Therefore, analogously to Eq. (S27), the integral operator $\tilde{M}_b^{(1)}\, p(\mathbf{x}, t)$ approaches the continuous transition operator defined in Eq. (S4) as $\epsilon \to 0$

$$\lim_{\epsilon\to 0} \tilde{M}_b^{(1)}\, p(\mathbf{x}, t) = \int d\mathbf{y}\, T(\mathbf{x}, t+\epsilon \,|\, \mathbf{y}, t)\, p(\mathbf{y}, t) = p(\mathbf{x}, t+\epsilon). \tag{S65}$$

To the best of our knowledge this proof is a novel mathematical result. We make no attempt to prove that this holds true for more general embedded Riemannian manifolds $\mathcal{M} \subset \mathbb{R}^d$, but conjecture that such a proof should be possible using the same techniques used in [6]. Just as in our construction of the discrete Laplace operator, a discrete transition operator can be defined from Monte Carlo estimates of the associated integral operators

$$\tilde{M}_b^{(\alpha)} f(\mathbf{x}) = \int_{\mathbb{R}^d} \frac{\tilde{K}_\epsilon^{(\alpha)}(\mathbf{x}, \mathbf{y})}{d_\epsilon^{(\alpha)}(\mathbf{x})}\, f(\mathbf{y})\, q(\mathbf{y})\, d\mathbf{y} \approx \frac{\frac{1}{N}\sum_{j=1}^N \tilde{K}_\epsilon^{(\alpha)}(\mathbf{x}_i, \mathbf{x}_j)\, f(\mathbf{x}_j)}{\frac{1}{N}\sum_{m=1}^N K_\epsilon^{(\alpha)}(\mathbf{x}_i, \mathbf{x}_m)} = \frac{\sum_{j=1}^N \tilde{K}_\epsilon^{(\alpha)}(\mathbf{x}_i, \mathbf{x}_j)\, f(\mathbf{x}_j)}{d_\epsilon^{(\alpha)}(\mathbf{x}_i)}, \tag{S66}$$

given a vector field $\mathbf{v}(\mathbf{x}_i)$ defined at each $\mathbf{x}_i \in \hat{\mathcal{M}}$. Specialization to the case of gradient-like drifts $\mathbf{v} = \mathbf{g}^{-1}\nabla U$ can be accomplished using the exact same scheme employed in the previous section.

##### S1.6.2 Connecting the two discretizations and the choice of volume element

We will now show that these two operators are numerically equivalent provided appropriate definitions of the volume element $V(\mathbf{x}_i)$. As mentioned in the main text, the default construction in our code is $V_B(\mathbf{x}_i) := 1/(N\, p_B(\mathbf{x}_i)) = \exp\left[U_B(\mathbf{x}_i)\right]/N$, where $p_B$ is the (base) kernel density estimate for the points that define our discrete manifold $\hat{\mathcal{M}} = \{\mathbf{x}\}_{i=1}^N$. This choice is intuitively appealing since for $p_B(\mathbf{x}_i) = \exp\left[-U_B(\mathbf{x}_i)\right]$ we recover a uniform probability

$$P(\mathbf{x}_i) = p_B(\mathbf{x}_i)\, V_B(\mathbf{x}_i) = 1/N. \tag{S67}$$

We pause to clarify that the diffusion coefficient in Eq. (S26) is *not* the same as the diffusion coefficient associated to the developmental dynamics we seek to model. It is just a property of the manifold $\mathcal{M}$, which we might refer to as $D_B$ since, for our discrete construction, it is solely related to the points in $\hat{\mathcal{M}}$ that define the base potential $U_B$ and not to the measured dynamics for any particular experimental condition. Without loss of generality we will set $D_B = 1$, a choice that was already implicitly made in our definition of $V_B(\mathbf{x}_i) := \exp\left[U_B(\mathbf{x}_i)/D_B\right]/N$. Another natural volume element is the diagonal elements of the mass matrix associated to the discrete Laplace-Beltrami operator $V_L(\mathbf{x}_i) := \sqrt{\epsilon}\, d_\epsilon^{(1)}(\mathbf{x})$ (Eq. (S41)). The connection between the discrete operators in Eqs. (S55) and (S66) clarifies some of the ambiguity in our choice of $V(\mathbf{x}_i)$. Recall that the matrix operator $\tilde{\mathbf{M}}_b^{(1)}$ in Eq. (S66) acts on discrete probability densities. We can consider constructing an associated transition operator that acts on discrete probabilities in the following way

$$\tilde{T}_\epsilon(\mathbf{x}_i, \mathbf{x}_j) = \frac{V_L(\mathbf{x}_i)}{\tilde{\mathcal{Z}}_j}\, \tilde{M}_b^{(1)}(\mathbf{x}_i, \mathbf{x}_j)\, \frac{1}{V_L(\mathbf{x}_j)}. \tag{S68}$$

In other words, given a probability distribution $P(\mathbf{x}_j)$, we compute the corresponding density $p(\mathbf{x}_j) = P(\mathbf{x}_j)/V_L(\mathbf{x}_j)$, evolve this density forward in time using $\tilde{\mathbf{M}}_b^{(1)}$, and then convert back to a probability distribution via multiplication by $V_L(\mathbf{x}_i)$. $V_L(\mathbf{x})$ is the natural choice of volume element in this context given the derivation of Eq. (S66). We also include a per-column normalization constant $\tilde{\mathcal{Z}}_j$ to ensure that $\tilde{\mathbf{T}}_\epsilon$ is a properly normalized left Markov matrix. Explicitly, we have

$$\tilde{T}_\epsilon(\mathbf{x}_i, \mathbf{x}_j) = \frac{1}{\tilde{\mathcal{Z}}_j} \frac{\tilde{K}_\epsilon^{(1)}(\mathbf{x}_i, \mathbf{x}_j)}{d_\epsilon^{(1)}(\mathbf{x}_j)} = \frac{1}{\tilde{\mathcal{Z}}_j} \frac{\exp\left[-\frac{||\mathbf{x}_i - \mathbf{x}_j||^2}{4D\epsilon} + \frac{(\mathbf{x}_i - \mathbf{x}_j)\cdot\mathbf{v}(\mathbf{x}_j)}{2D} - \frac{\epsilon\,||\mathbf{v}(\mathbf{x}_j)||^2}{4D}\right]}{q_\epsilon(\mathbf{x}_i)\, q_\epsilon(\mathbf{x}_j)} \frac{1}{d_\epsilon^{(1)}(\mathbf{x}_j)}. \tag{S69}$$

Normalization $\sum_i \tilde{T}_\epsilon(\mathbf{x}_i, \mathbf{x}_j) = 1$ demands that

$$\tilde{\mathcal{Z}}_j = \frac{\exp\left[-\frac{\epsilon\,||\mathbf{v}(\mathbf{x}_j)||^2}{4D}\right]}{q_\epsilon(\mathbf{x}_j)\, d_\epsilon^{(1)}(\mathbf{x}_j)} \sum_{i=1}^{N} \frac{\exp\left[-\frac{||\mathbf{x}_i - \mathbf{x}_j||^2}{4D\epsilon} + \frac{(\mathbf{x}_i - \mathbf{x}_j)\cdot\mathbf{v}(\mathbf{x}_j)}{2D}\right]}{q_\epsilon(\mathbf{x}_i)} \tag{S70}$$

and substituting this back into Eq. (S69) yields

$$\begin{aligned} \tilde{T}_\epsilon(\mathbf{x}_i, \mathbf{x}_j) &= \frac{\frac{1}{q_\epsilon(\mathbf{x}_i)} \exp\left[-\frac{||\mathbf{x}_i - \mathbf{x}_j||^2}{4D\epsilon} + \frac{(\mathbf{x}_i - \mathbf{x}_j)\cdot\mathbf{v}(\mathbf{x}_j)}{2D}\right]}{\sum_{m=1}^{N} \frac{1}{q_\epsilon(\mathbf{x}_m)} \exp\left[-\frac{||\mathbf{x}_m - \mathbf{x}_j||^2}{4D\epsilon} + \frac{(\mathbf{x}_m - \mathbf{x}_j)\cdot\mathbf{v}(\mathbf{x}_j)}{2D}\right]} \\ &= \frac{V_B(\mathbf{x}_i) \exp\left[-\frac{||\mathbf{x}_i - \mathbf{x}_j||^2}{4D\epsilon} + \frac{(\mathbf{x}_i - \mathbf{x}_j)\cdot\mathbf{v}(\mathbf{x}_j)}{2D}\right]}{\sum_{m=1}^{N} V_B(\mathbf{x}_m) \exp\left[-\frac{||\mathbf{x}_m - \mathbf{x}_j||^2}{4D\epsilon} + \frac{(\mathbf{x}_m - \mathbf{x}_j)\cdot\mathbf{v}(\mathbf{x}_j)}{2D}\right]}, \end{aligned} \tag{S71}$$

which exactly matches Eq. (S55) after specialization to gradient-like drift velocities and application of our finite-difference approximation scheme given the choice $V(\mathbf{x}_i) = V_B(\mathbf{x}_i)$. To arrive at the final equality, we used the fact that $q_\epsilon(\mathbf{x}_i) \propto p_B(\mathbf{x}_i)$ and simply canceled the overall constant factors.

So which is the correct volume element? Clearly, in the context of the Laplace-Beltrami operator we must use $V_L(\mathbf{x}_i)$. However, Eq. (S71) suggests that the appropriate volume element for our discrete dynamical transition operator in Eq. (S55) should be $V_B(\mathbf{x}_i)$. This is the primary reason that $V_B(\mathbf{x}_i)$ is the default choice in our code. Empirically, the two produce very similar fit results under the majority of conditions. See Fig. S2 for an illustration using the discrete manifold $\hat{\mathcal{M}}$ built for the synthetic data example in Algorithm Applied To Simulated Data (see Fig. 3B). The two elements share similar values in dense regions, but differ in sparse regions, with $V_B$ typically having a much larger dynamic range. Sparse regions, however, are typically void of probability during simulations anyway, which explains why the time courses of probability fit using the two different volume elements remain similar. In high dimensions (e.g. our RNA-seq example) and for very sparse points, the exponential $V_B(\mathbf{x}_i)$ can rapidly diverge and so the renormalized $V_L(\mathbf{x}_i)$ produces more stable results at smaller $\epsilon$.

### S2 Materials

Detailed descriptions of the experimental materials and methods used to generate the data in this paper can be found in [19, 20].

### S3 Methods

#### S3.1 Synthetic data generation

In the main text section Algorithm Applied To Simulated Data, we use our method to fit the dynamical structure of a landscape derived from the heteroclinic flip potential

$$U(x, y \mid a, b) = x^4 + y^4 + x^3 - 2\,x\,y^2 - x^2 + a\,x + b\,y \tag{S72}$$

at three different values of $(a, b) = \{(-1.6,\ -0.4),\ (-1.4,\ 0.5), (-1.1,\ -0.1)\}$. We for each set of $(a, b)$-values, we simulated 60,000 points using the Euler-Maryuama method with a time step of $dt = 0.0001$ according to Eq. (S1) with $\mathbf{g} = \mathbf{I}$ and $D = 1$. The initial conditions for all points were randomly drawn from a Gaussian distribution with variance $\sigma = 0.05$ centered on the fixed point of $U(x, y \mid a, b)$ with negative $x$-value ($(x, y) \sim (-1, 0)$). For $(a, b) = (-1.6,\ -0.4)$, which has no fixed point with negative $x$-value, we use the corresponding fixed point from $(a, b) = (-1.4,\ 0.5)$. At five non-uniformly spaced time points $t \in \{0, 0.2, 0.8, 1.6, 5\}$, corresponding to the arbitrarily labeled $T$ in main text Fig. 3, we randomly select 12,000 points as a representative sample for that time point and remove them from the simulation. This was done to prevent any issues with possible correlations if we had just simulated 12,000 points and reported their positions at the requisite time points. This is functionally the same as starting five separate simulations of 12,000 points and running them to the specified end points, although marginally faster by taking advantage of MATLAB's vectorization optimization. This also mirrors real experiments where the cell profiles captured on Day 4, say, are completely distinct from cells at any other measured time points and are not simply the time-evolved states of the cell profiles captured on Day 3, for instance.

#### S3.2 Subsampling methods for building $\hat{\mathcal{M}}$ from data

Our code comes packaged with two methods for subsampling point clouds to produce a discrete representation of the underlying dynamical manifold $\hat{\mathcal{M}}$: (1) simple uniform random subsampling and (2) SPARTAN [21], an optimal transport based method specifically designed to improve convergence of kernel density estimation on the subsampled point clouds over uniform sampling. Briefly, SPARTAN uses a fast approximate optimal transport computation [22] to compute a map $\phi(\mathbf{x}) : \mathbb{R}^d \to U[0, 1]^d$ from the high-dimensional space where the point cloud lives to the uniform distribution over the unit hypercube. $N$ quasirandom uniform samples (see, for instance, MATLAB's `sobolset`) are then drawn over $U[0, 1]^d$ and the $N$ unique points whose mapped locations $\phi(\mathbf{x})$ are closest to these quasirandom feature points are picked as subsamples of the original cloud.

The primary concerns when picking points comprising $\hat{\mathcal{M}}$ are to ensure broad coverage of the relevant regions of gene expression space, such that faint yet important features are not drowned out, and that regions that are heavily populated solely due to the structure of the experiment (i.e. the initial conditions) do not dominate the base potential $U_B$. For our synthetic data example (Appendix S3.1), we used SPARTAN to sample $\{200, 300, 300, 300, 900\}$ points from each time point, respectively, and for each set of $(a, b)$-values for a total of 6000 points out of 180,000 initial points. This was done to ensure that the terminal distributions (which were different for each set of $(a, b)$-values) was not overwhelmed by the initial condition (which was effectively identical for all parameter values). For the 5D flow cytometry data, the full point set consisted of 1,109,152 cells pooled together from seven data sets, including the constant SAG experiments, the SAG 0-500 and SAG 500-0 experiments, and the SAG 0-500-0 experiment. The subsampled $\hat{\mathcal{M}}$ consisted of 8000 points obtained using SPARTAN, with

$\{1500, 1000, 2000, 3500\}$ sampled on Days, 3, 4, 5, and 6 respectively. For the RNA-seq analysis, we just used uniform random sampling to reduce the 7293 cells processed by Fontaine et al. [20] down to 4000 cells. Complete specification of all data analysis choices is available in our codebase.

##### S3.3 Topological data analysis methods for estimating extrema

Consider a scalar function $f_i := f(\mathbf{x}_i)$ defined on an unstructured point set $\hat{\mathcal{M}} = \{\mathbf{x}_i\}_{i=1}^N \in \mathbb{R}^d$ (e.g. a pointwise density or potential). The extrema of these functions can be robustly identified using topological data analysis techniques [23]. For concreteness here, we focus on finding the local minima of the $f_i$. Locating the local maxima be accomplished by applying the exact same minimum finding procedure after a sign-flip and a suitable constant shift. The first step in this procedure is the construction of a proximity graph, i.e. a set of points and edges $\{\hat{\mathcal{M}}, E\}$ which define connectivity between points based on local neighborhood structure. Familiar examples are the k-NN graph or the Delaunay graph. For our purposes, we use the relaxed Gabriel graph. For two points $\mathbf{x}_i$, $\mathbf{x}_j \in \hat{\mathcal{M}}$, the standard Gabriel graph contains an edge $E_{ij} \in E$ connecting $\mathbf{x}_i$ and $\mathbf{x}_j$ if the open ball of diameter $||\mathbf{x}_i - \mathbf{x}_j||$ centered on the midpoint $\mathbf{m}_{ij} = (\mathbf{x}_i + \mathbf{x}_j)/2$ contains no other points, i.e.

$$||\mathbf{x}_i - \mathbf{x}_k||^2 + ||\mathbf{x}_j - \mathbf{x}_k||^2 \geq ||\mathbf{x}_i - \mathbf{x}_j||^2, \quad \forall\, \mathbf{x}_k \in \hat{\mathcal{M}}. \tag{S73}$$

Correa and Lindstrom define a relaxed Gabriel graph that is constructed by sequentially applying the condition Eq. (S73) to points $\mathbf{x}_k$ in order of increasing distance from $\mathbf{x}_i$ and only rejecting edges $E_{ik}$ if Eq. (S73) is violated with respect to previously accepted neighbors [24]. The standard Gabriel graph is therefore always a subset of the relaxed Gabriel graph, which, in turn, is a subset of the Delaunay graph. We also ensure that the graph always contains only one connected component, which can be accomplished, for instance, by joining disconnected points to their nearest neighbor.

Once the proximity graph is built, we construct the corresponding join tree with respect to the $f_i$. The join tree is a topological data structure that captures how the basins of attraction of the local minima (regions connected by the proximity graph in which all points flow downhill to the same local minimum) merge as we progressively raise a threshold $c$ defining the sub-level sets $\{i : f_i \leq c\}$. Practically, points are sorted from lowest to highest $f_i$, and each point is processed sequentially. Initially, all basins are disconnected, i.e there is no way to travel from one local minima to another along edges of the proximity graph between vertices contained within the incomplete join tree. As we continue to add points into the join tree, we eventually reach vertices that join two previously disconnected basins, which triggers a merge event that connects two branches in the tree. This scheme results in a tree with multiple branches, whose roots are the local minima of the proximity graph, internal nodes (join saddles) marking basin mergers, and a single leaf at the global maximum, at which all basins ultimately combine.

Once the join tree is constructed, we can perform a robust extraction of the "true" minima by pruning branches that fail to meet certain topological quality standards. The most important such criteria for our purposes are size (number of points in a branch), stability (sum of function values along a branch), and persistence (defined here as the difference between the maximum and minimum values along a branch). Branches that fail to meet user specified thresholds in these quantities are systematically merged until only a small set of prominent minima remain.

##### S3.4 Determining unstable manifolds

In this work, we restrict our attention to systems of the form Eq. (S1) with $\mathbf{v}(\mathbf{x}, t) = -(\nabla U(\mathbf{x})/g)$, where $g$ is a spatially uniform scalar that can be absorbed into $U$. For such systems, i.e. gradient dynamics with additive noise, the unstable manifolds of the deterministic component of the dynamics are equivalent to the most probable paths between fixed points in the limit $D \to 0$ [25, 26]. Discrete approximations to these analytic manifolds can be found as the most probable paths between points in $\hat{\mathcal{M}}$ for small $D$. In particular the total probability of transitioning between two points $\mathbf{x}_i^*$ and $\mathbf{x}_j^*$ along the path $\mathcal{W}_B(\mathbf{x}_i^*, \mathbf{x}_j^*) = \{\mathbf{x}_1 = \mathbf{x}_i^*, \mathbf{x}_2, \ldots, \mathbf{x}_{|\mathcal{W}_B|} = \mathbf{x}_j^*\}$, where $|\mathcal{W}_B|$ is the number of points in the path, is given by

$$\begin{aligned} P(\mathbf{x}_i^* \to \mathbf{x}_j^* | \mathcal{W}_B) &= \prod_{k=1}^{|\mathcal{W}_B|-1} T(\mathbf{x}_{k+1}, \mathbf{x}_k) \\ \implies -\log P(\mathbf{x}_i^* \to \mathbf{x}_j^* | \mathcal{W}_B) &= \sum_{k=1}^{|\mathcal{W}_B|-1} \left(-\log T(\mathbf{x}_{k+1}, \mathbf{x}_k)\right) \end{aligned} \tag{S74}$$

The minus log components $-\log T_\epsilon(\mathbf{x}_i, \mathbf{x}_j)$ of the transition operator can be thought of as defining the (non-symmetric) weights of a directed graph. Note that these components are always positive since all $0 < T_\epsilon(\mathbf{x}_i, \mathbf{x}_j) < 1$. The shortest path between any two specified points $\mathbf{x}_i^*$ and $\mathbf{x}_j^*$ along the edges of this weighted directed graph will therefore always correspond to the path with the greatest total transition probability $P(\mathbf{x}_i^* \to \mathbf{x}_j^* | \mathcal{W}_B)$. These shortest paths can be found using any number of well established graph shortest path algorithms (e.g. Dijkstra's algorithm - see, for instance, MATLAB's `shortestpath`). Note that for more general dynamical systems, the most probable path will typically no longer coincide with the unstable manifolds (see [27] for examples and discussion).

##### S3.5 Stable solution of the Fokker-Planck equation

Approximate solutions to Eq. (S3) can be computed by iteratively acting the transition operator $\mathbf{T}_\epsilon$ on an initial probability vector

$$P(\mathbf{x}_i, t_{k+1}) = \sum_{j=1}^{N} T_\epsilon(\mathbf{x}_i, \mathbf{x}_j)\, P(\mathbf{x}_j, t_k). \tag{S75}$$

For small $\epsilon$ or small $D$, the repeatedly multiplied exponential factors in $T_\epsilon^k(\mathbf{x}_i, \mathbf{x}_j)$ can rapidly underflow or overflow due to finite numerical precision, leading to degradation in the quality of the results. We employ a commonly used log-stabilization trick to avoid these problems. From Eq. (S75), we see that the time evolution of the log probabilities is given by

$$\begin{aligned} \log P(\mathbf{x}_i, t_{k+1}) &= \log \left[ \sum_{j=1}^{N} \exp\left[\log T_\epsilon(\mathbf{x}_i, \mathbf{x}_j) + \log P(\mathbf{x}_j, t_k)\right] \right] \\ &= \mathrm{LSE}\left[\log T_\epsilon(\mathbf{x}_i, \mathbf{x}_j) + \log P(\mathbf{x}_j, t_k)\right], \end{aligned} \tag{S76}$$

where LSE denotes the LogSumExp function, a familiar operation in numerical methods that can be computed in a way that prevents numerical underflow or overflow [28]. Solutions to Eq. (S3) can therefore be computed stably in the log-domain and then converted back via exponentiation after a predetermined number of time steps. A completely analogous operation allows us to stably compute solutions to the backward Kolmogorov equation.

##### S3.6 Computation of basins of attraction

Let $\psi(\mathbf{x}) : \mathbb{R}^d \to \mathbb{R}$ be some scalar function defined on $\mathbb{R}^d$ that we evaluate at time $t$ based on the position of a cell $\mathbf{x}(t)$, which evolves according to Eq. (S1). The backwards Kolmogorov equation (BKE) tells us the average value of this function given that the trajectory of the cell began at location $\mathbf{x}(s)$ at time $s \leq t$. When $\psi(\mathbf{x}) = \mathbf{1}_{\mathcal{U}}$ is the indicator function for a region $\mathcal{U} \subset \mathbb{R}^d$, i.e. it equals 1 if $\mathbf{x} \in \mathcal{U}$ and zero otherwise, the BKE tells us which locations can possibly be connected to the target set by a stochastic trajectory beginning at $\mathbf{x}(s)$. The higher the value, the more likely it is that such a trajectory reaches $\mathcal{U}$ at time $t$. Approximate solutions can be found simply by using the transpose transition operator

$$\psi(\mathbf{x}_i, t_{k-1}) = \sum_{j=1}^{N} T_\epsilon(\mathbf{x}_j, \mathbf{x}_i)\, \psi(\mathbf{x}_j, t_k). \tag{S77}$$

For a given potential $U_D$ and corresponding transition operator $\mathbf{T}_\epsilon$, let $\{\mathbf{x}^*_m\}$ denote the stable fixed points of the dynamics. These can be found by using topological extrema identification methods to locate the minima of $U_D$ (or of the effective potential $U_{eff} = -g\, D \log[p_{eq}(\boldsymbol{\theta})]$, where $p_{eq}(\boldsymbol{\theta})$ is the equilibrium density of $\mathbf{T}_\epsilon$). See Appendix S3.3 for more details. We can then iteratively solve the BKE with the trivial indicator function $\mathbf{1}_{\mathbf{x}^*_m}$ using the stable method described in Appendix S3.5. The default number of time steps is 50. We can then associate each $\mathbf{x}_i \in \hat{\mathcal{M}}$ to one of the $\{\mathbf{x}^*_m\}$ by asking which fixed point had the highest corresponding BKE value at $\mathbf{x}_i$. In the continuum limit ($N \to \infty$, $\epsilon \to 0$) and for $D \to 0$ this method will approach the true dynamical basins of attraction.

##### S3.7 Choice and computation of loss function

Our framework allows users to pick one of six different loss functions to compare simulated pointwise probabilities $P^{(sim)}$ to the measured pointwise probabilities $P^{(meas)}$. The Kullback-Leibler divergence for two distributions $P$ and $Q$ is defined to be

$$D_{KL}(P||Q) = \sum_{i=1}^{N} P(\mathbf{x}_i) \log\left[\frac{P(\mathbf{x}_i)}{Q(\mathbf{x}_i)}\right]. \tag{S78}$$

Users can choose from $D_{KL}(P^{(meas)}||P^{(sim)})$ ('reverse' K-L divergence), $D_{KL}(P^{(sim)}||P^{(meas)})$ ('forward' K-L divergence), or the symmetrized K-L divergence

$$(D_{KL}(P^{(meas)}||P^{(sim)}) + D_{KL}(P^{(sim)}||P^{(meas)}))/2. \tag{S79}$$

Users can also use the mean squared error

$$\text{MSE}(P^{(meas)}||P^{(sim)}) = \frac{1}{N}\sum_{i=1}^{N} ||P^{(meas)}(\mathbf{x}_i) - P^{(sim)}(\mathbf{x}_i)||^2, \tag{S80}$$

which treats the distributions as vectors in $\mathbb{R}^N$. We also implement a 'geodesic' error metric. Since $\sum_i P(\mathbf{x}_i) = \sum_i (\sqrt{P(\mathbf{x}_i)})^2 = 1$ for all probability distributions, the vector whose elements are the square-roots of the probabilities lives on the $(N-1)$-sphere in $N$-dimensional space. Our geodesic error metric is defined to be

$$\text{GEO}(P^{(meas)}||P^{(sim)}) = \cos^{-1}\left[\sum_{i=1}^{N} \sqrt{P^{(meas)}(\mathbf{x}_i)\, P^{(sim)}(\mathbf{x}_i)}\right], \tag{S81}$$

which is just the geodesic distance between the two corresponding points on the $(N-1)$-sphere. Finally, we also provide an implementation of the weighted optimal mass transport distance, although the computation of this error metric is too slow to be of any practical use. The symmetrized $D_{KL}$ is the default loss function in our code. Empirically, we observe that the symmetrized $D_{KL}$, MSE, and GEO loss functions all produce similar, high-quality results with similar computation times. The reverse and forward K-L divergences have a known tendency to be mean or mode seeking, respectively, and users should have a specific reason to prefer them over their symmetrized counterpart. For all examples in this work, we used the symmetrized $D_{KL}$ as the error metric to produce our fits. In tables and figures, however, we report the reverse K-L divergence $D_{KL}(P^{(meas)}||P^{(sim)})$.

For each data set and experimental condition, we simulate $k_{max}$-steps (a user specified positive integer), which produces $k_{max}+1$ column vectors $P_0^{(sim)}$, $P_1^{(sim)}$, ... , $P_{k_{max}}^{(sim)}$. These must be compared with the $K$ measured probability distributions $P_0^{(meas)}$, $P_1^{(meas)}$, ... , $P_{K_{max}}^{(meas)}$, i.e. the probability distributions on Day 0 to Day $K_{max}$. Typically we use the measured condition at the earliest time point as the initial condition for the simulations so $P_0^{(meas)} = P_0^{(sim)}$. By default, we associate a $P_k^{(sim)}$ to each $P_K^{(meas)}$ by just exhaustively searching over the error for all simulated probability distributions and choosing the one that minimizes the error for that measured time point. Technically, this means that it is possible for measured distributions $P_{K_1}^{(meas)}$ and $P_{K_2}^{(meas)}$ to be associated to two simulated distributions $P_{k_1}^{(sim)}$ and $P_{k_2}^{(sim)}$ where $K_1 < K_2$ and $k_2 < k_1$. In practice, such violations of causality never occurred during the fitting of any synthetic or experimental data set. We do, however, do include a non-default option to perform more stringent checks so that causality violations never occur. Finally, with all this in hand, the total loss at a given optimization step is the average loss over all data sets, experimental conditions, and time points.

##### S3.8 Converting simulation time to experimental time

Our method produces approximate solutions to Eq. (S3) at discrete simulated times $t_k^{(sim)} = k\epsilon$. To clearly distinguish simulated and experimental in this section, we explicitly label all times and distributions accordingly. Given a data set containing $K$ measured probability distributions $P_I^{(meas)}$, $I = 1, \dots, K$, our fitting procedure identifies corresponding simulated distributions $P_I^{(sim)}(t_{k_I}^{(sim)})$ that best match these measured data. We assume a monotonically increasing relationship between indices, such that if $I < J$, then $t_{k_I}^{(sim)} < t_{k_J}^{(sim)}$ (see Appendix S3.7). To map simulated times $t^{(sim)}$ to experimental times $t^{(exp)}$ (e.g., measured in days), we use a simple power-law relationship:

$$t^{(exp)} = c_1 \left(t^{(sim)}\right)^{c_2} + c_3. \tag{S82}$$

For all data sets, the parameters $c_1$, $c_2$, and $c_3$ are determined by minimizing the average error between measured and simulated distributions. For data sets with only three experimental time points (such as SAG 0 and SAG 10, where we exclude aging effects, and the SAG 0-500 and SAG 500-0 two-stage fits), these parameters are trivially determined by explicitly matching the three simulated points that minimize this error. For data sets with more than three experimental time points (e.g., SAG 100, SAG 500, and RNA-seq data), we perform a more general optimization procedure using MATLAB's `fmincon`. This approach utilizes either a hard assignment (selecting the closest simulated time point) or a soft assignment (softmax-weighted averaging across simulated points), allowing for robust fitting even when later time points exhibit broad, shallow error minima due to equilibration (see, for example, Fig. S9D-F).

##### S3.9 Converting the linear interpolation on fit potentials to SAG concentrations

Recognizing that landscapes must depend on morphogen levels, we imposed a simple functional constraint to interpolate SAG levels in the flow cytometry data. SAG 10 nM and SAG 500 nM were each endowed with a complete set of optimizable fit parameters, but the dynamics at the intermediate SAG 100 nM was found by linearly interpolating between the dynamical landscapes:

$$U_D^{(100)}(\mathbf{x}_i) = (1 - \alpha_{100})\,\frac{U_D^{(10)}(\mathbf{x}_i)}{g^{(10)}} + \alpha_{100}\,\frac{U_D^{(500)}(\mathbf{x}_i)}{g^{(500)}}, \tag{S83}$$

where $\alpha_{100} = 0.637$ is a single interpolation parameter (note the slight change of notation from the main text) for all points in the discrete manifold and $g^{(100)} = 1$. The scalar diffusion coefficient was interpolated similarly as $D^{(100)} = (1 - \alpha)\,D^{(10)} + \alpha\,D^{(500)}$. A conversion from the abstract linear interpolation parameter $\alpha$ to concrete SAG concentration levels is necessary to make out-of-sample predictions (e.g. Fig. 8). Based on the observation that the dynamics are roughly saturating above SAG 500 nM (data not shown) we use the following functional form

$$\mathrm{SAG}(\alpha) = \frac{b\,(\alpha - c)}{c + a - \alpha}, \tag{S84}$$

where $a$, $b$, and $c$ are constants that can be uniquely solved for by enforcing the conditions that $\mathrm{SAG}(0) = 10$, $\mathrm{SAG}(\alpha_{100}) = 100$, and $\mathrm{SAG}(1) = 500$. For values of SAG between 0 nM and 10 nM, we just linearly interpolate between the landscapes fit to those concentrations (see Fig. 8).

##### S3.10 Moving least squares for derivative computation

We implemented a moving least-squares (MLS) method to compute gradients of scalar functions defined on unstructured point clouds. A more complete expanation of MLS methods can be found in [29]. Briefly, given $N$ points at positions $\mathbf{x}_i \in \mathbb{R}^d$, the goal of MLS methods are to obtain a globally defined function $f(\mathbf{x})$ that approximates the scalar values $f_i$ defined at the points $\mathbf{x}_i$. For our purposes, we obtain this global function as the solution to the following optimization problem

$$f(\mathbf{x}) := f_{\mathbf{x}}(\mathbf{x}) = \underset{f_{\mathbf{x}} \in \Pi_m^d}{\operatorname{argmin}} \sum_{i=1}^{N} \theta(||\mathbf{x} - \mathbf{x}_i||)\,||f_{\mathbf{x}}(\mathbf{x}_i) - f_i||^2, \tag{S85}$$

where $f_{\mathbf{x}}$ is drawn from $\Pi_m^d$, the space of polynomials of total degree $m$ in $d$ spatial dimensions, and $\theta(d)$ is a non-negative weight function that sets the relative importance of each $f_i$ in shaping the global function. Following [29], we use the notation $f_{\mathbf{x}}(\mathbf{x})$ to emphasize that this function is redefined locally for each query point $\mathbf{x}$ through the weight function $\theta(d)$. For all analysis in this work, we used a simple Gaussian weight function $\theta(d) = \exp[-d^2/4\,\epsilon]$, where the bandwidth $\epsilon$ is the same one used to build the transition operator $\mathbf{T}_\epsilon$. We note that our implementation enables users to fit polynomials of arbitrary spatial dimension $k \leq d$ by first fitting a $k$-plane to a local neighborhood around $\mathbf{x}$ and then optimizing Eq. (S85), however, for our purposes we always just used the full dimensions of the ambient space. We used this MLS compute the stability of the various fixed points in our fit potentials. For a given dynamical potential $U_D$ and scalar metric $g$ the elements of the Jacobian of the velocity are given by

$$J_{ij} = -\frac{\partial^2 (U_D(\mathbf{x})/g)}{\partial x_i \partial x_j}. \tag{S86}$$

It is trivial to then compute the eigensystem of this matrix for each putative sink and saddle to determine their stability. We note for completeness that all saddles fit to both synthetic and experimental data were index-1 saddles with only one positive eigenvalue.

##### S3.11 Computation of direct probability flow from NMP to p3

The components of the transition operator $T_\epsilon(\mathbf{x}_i, \mathbf{x}_j)$ define the probability of transitioning from $\mathbf{x}_j$ to $\mathbf{x}_i$ in one step of our discrete Markov process. Let $\hat{\mathcal{M}}_{inc} = \text{NMP} \cup \text{p0/p1} \cup \text{p2} \cup \text{p3}$ denote the union of the points $\mathbf{x}_i \in \hat{\mathcal{M}}$ assigned to the corresponding basins using the method in Appendix S3.6. Let $\hat{\mathcal{M}}_{exc} = \text{pMN} \cup \text{DP}$ denote the complement of $\hat{\mathcal{M}}_{inc}$ such that $\hat{\mathcal{M}}_{inc} \cup \hat{\mathcal{M}}_{exc} = \hat{\mathcal{M}}$ and $\hat{\mathcal{M}}_{inc} \cap \hat{\mathcal{M}}_{exc} = \emptyset$. We define $\hat{\mathcal{M}}_{inc}$ in this way since the measured probability in the flow cytometry data on Day 3, $P(\mathbf{x}, 0)$, has a significant amount of probability in the p0/p1 basin and a small, but non-negligible, amount of probability in p2 basin (Fig. 6E-H). Let $\mathbf{1}_{inc}$ denote the column vector whose elements are one if they correspond to points in $\hat{\mathcal{M}}_{inc}$ and zero otherwise. Then $\hat{\mathbf{P}}_{inc} = \text{diag}[\mathbf{1}_{inc}]$ is a projection operator onto $\hat{\mathcal{M}}_{inc}$. The matrix $\mathbf{T}_\epsilon^{inc} = \hat{\mathbf{P}}_{inc}\, \mathbf{T}_\epsilon\, \hat{\mathbf{P}}_{inc}$ is an unnormalized, projected transition operator that zeros flow into and out of $\hat{\mathcal{M}}_{exc}$. The total probability in the p3 basin at the $k$th simulation step is of course

$$P(\text{p3},\, t_k) = \mathbf{1}_{\text{p3}}^T\, (\mathbf{T}_\epsilon)^k\, \mathbf{P}(0), \tag{S87}$$

while the probability in the p3 basin that never entered $\hat{\mathcal{M}}_{exc}$ is given by

$$P(\text{p3},\, t_k \,|\, \hat{\mathcal{M}}_{inc}) = \mathbf{1}_{\text{p3}}^T \left(\mathbf{T}_\epsilon^{inc}\right)^k \hat{\mathbf{P}}_{inc}\, \mathbf{P}(0). \tag{S88}$$

This becomes our definition of the 'direct' route from NMP to p3. Using the landscape fit to the SAG 500 nM flow cytometry data (using the interpolated SAG 10-100-500 protocol), for which Day 4 corresponds to the $k = 8$, we have $P(\text{p3, Day 4} \,|\, \hat{\mathcal{M}}_{inc}) = 0.053$ and $P(\text{p3, Day 4}) = 0.058$, which has a ratio of 0.92. This is equivalent to the ratio we would approach if we simulated many $k$-step single cell Monte Carlo trajectories beginning in $\hat{\mathcal{M}}_{inc}$ and asked which fraction terminated in the p3 basin without ever entering $\hat{\mathcal{M}}_{exc}$.

##### S3.12 Dynamical diffusion map embeddings

The dynamical diffusion map embedding used in Fig. 9 is a nonlinear dimensional reduction technique that is related to the standard diffusion map embedding (see Appendix S1.3). Given a dynamic transition operator $\mathbf{T}_\epsilon$ that we fit to data, we take the embedding coordinates to be

$$\tilde{\mathbf{\Phi}}_k(\mathbf{x}_i) = \left(\lambda_1^k\, \varphi_1(\mathbf{x}_i),\, \lambda_2^k\, \varphi_2(\mathbf{x}_i),\, \ldots,\, \lambda_n^k\, \varphi_n(\mathbf{x}_i)\right), \tag{S89}$$

where $\varphi_i$ is the $i$th right eigenvector of $\mathbf{T}_\epsilon$, $\lambda_i$ is the associated eigenvalue, and $n = 2$ or 3. Euclidean distances in these coordinates no longer have the interpretation of approximate diffusion distances, as they do for standard diffusion maps [5], but they do highlight important dynamical features in the system. Empirically, they compress unstable regions towards the origin and highlight the stable states, which appear at the ends of roughly orthogonal lines emanating from the origin.

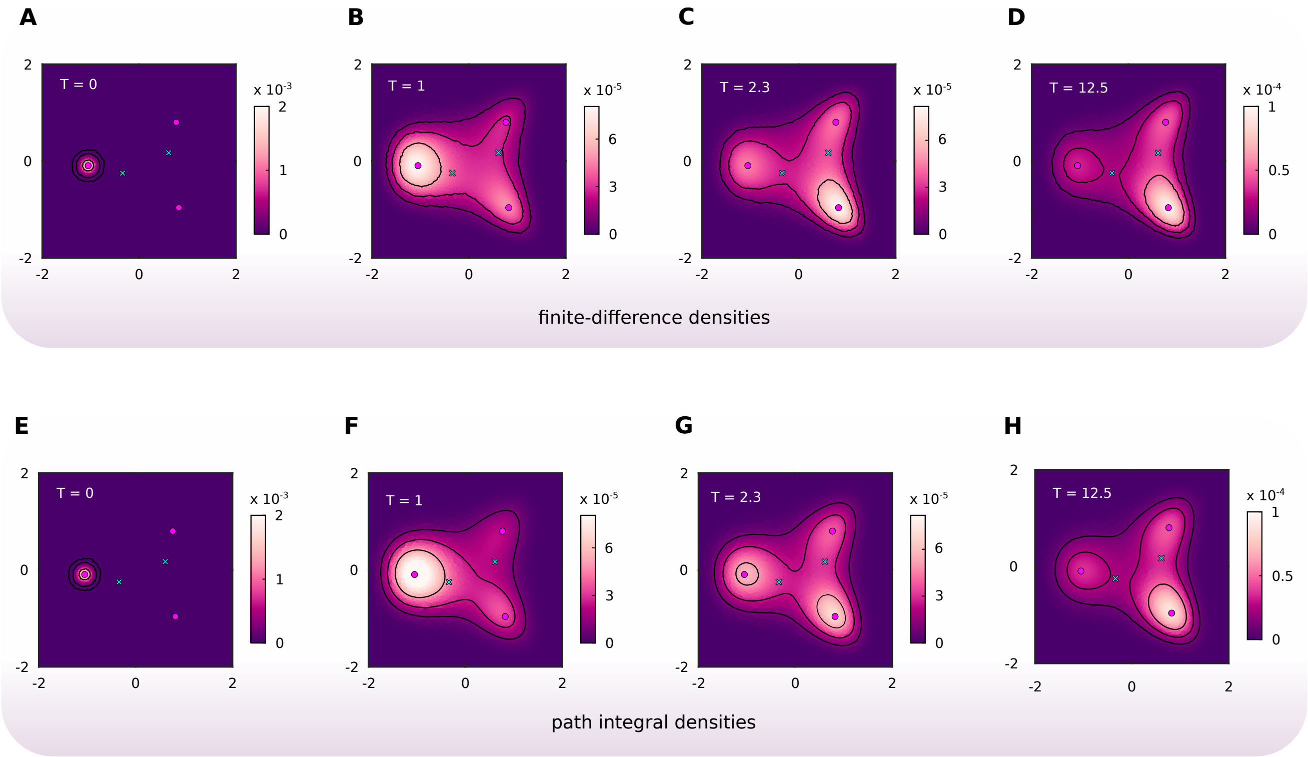

**Figure S1.** A time course of probability density solutions to the Fokker-Planck equation for the example described in Mathematical preliminaries and Fig. 2. (A)-(D) The results of solving Eq. (S3) using finite-difference methods on a $100 \times 100$ regular grid over the domain $[-2, 2] \times [-2, 2]$. Finite-difference solution are then transferred onto the inhomogenous point used to define $\hat{\mathcal{M}}$ via area weighted averaging. (E)-(H) The results of solving Eq. (S3) using our discrete path integral formalism directly on $\hat{\mathcal{M}}$

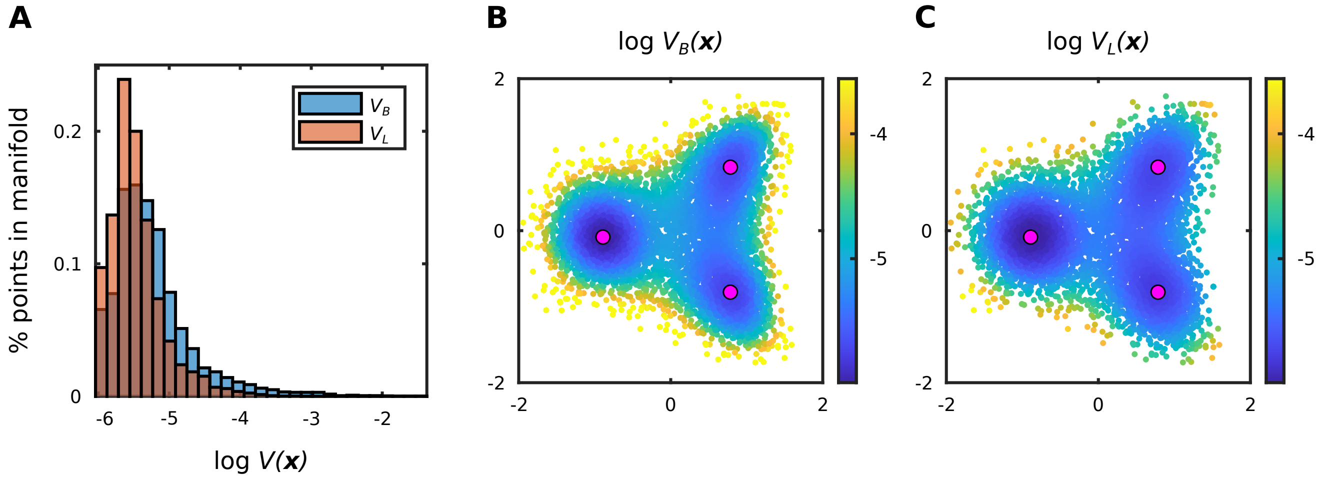

**Figure S2.** A comparison of the different volume elements for the discrete manifold $\hat{\mathcal{M}}$ built for the synthetic data example in Algorithm Applied To Simulated Data (see Fig. 3B). See Appendix S1.6.2 for a detailed discussion of the different volume elements. (A) A histogram comparing the log-values of the volume elements $V_B$ and $V_L$. (B) Shifted log-values of the volume element $V_B = 1/(N\,p_B)$, where $N$ is the the number of points in $\hat{\mathcal{M}}$ and $p_B$ is the base density of $\hat{\mathcal{M}}$. The log-values have been shifted so that the minimum aligns with the minimum of $\log V_L$ in (C). Color range is saturated between the 2nd and 98th percentile. (C) Log-values of the volume element $V_L$. See Appendices S1.4.3 and S1.6.2 for an explicit definition. Color range is identical to the one in (B).

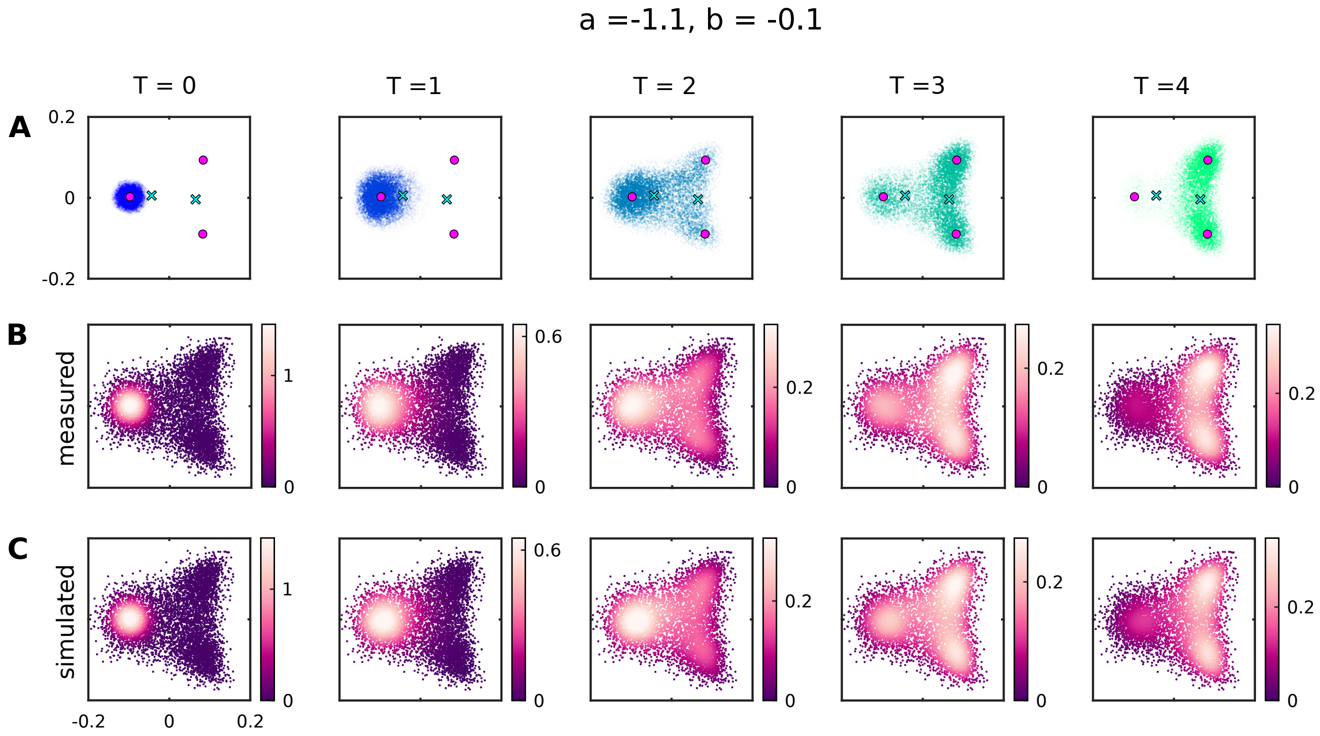

**Figure S3.** Time course of results of the synthetic example discussed in Algorithm Applied To Simulated Data and Figs. 3 and 4 for parameter values $a = -1.1$ and $b = -0.1$. Errors and entropies are reported in Table S1. (A) The actual simulated points at times $T = 0$ to 4 (see Appendix S3.1). Each time point contains 12,000 unique points. (B) The time course of measured densities on $\hat{\mathcal{M}}$ computed using kernel density estimation with the points in (A) as sources. (C) Time course of simulated densities.

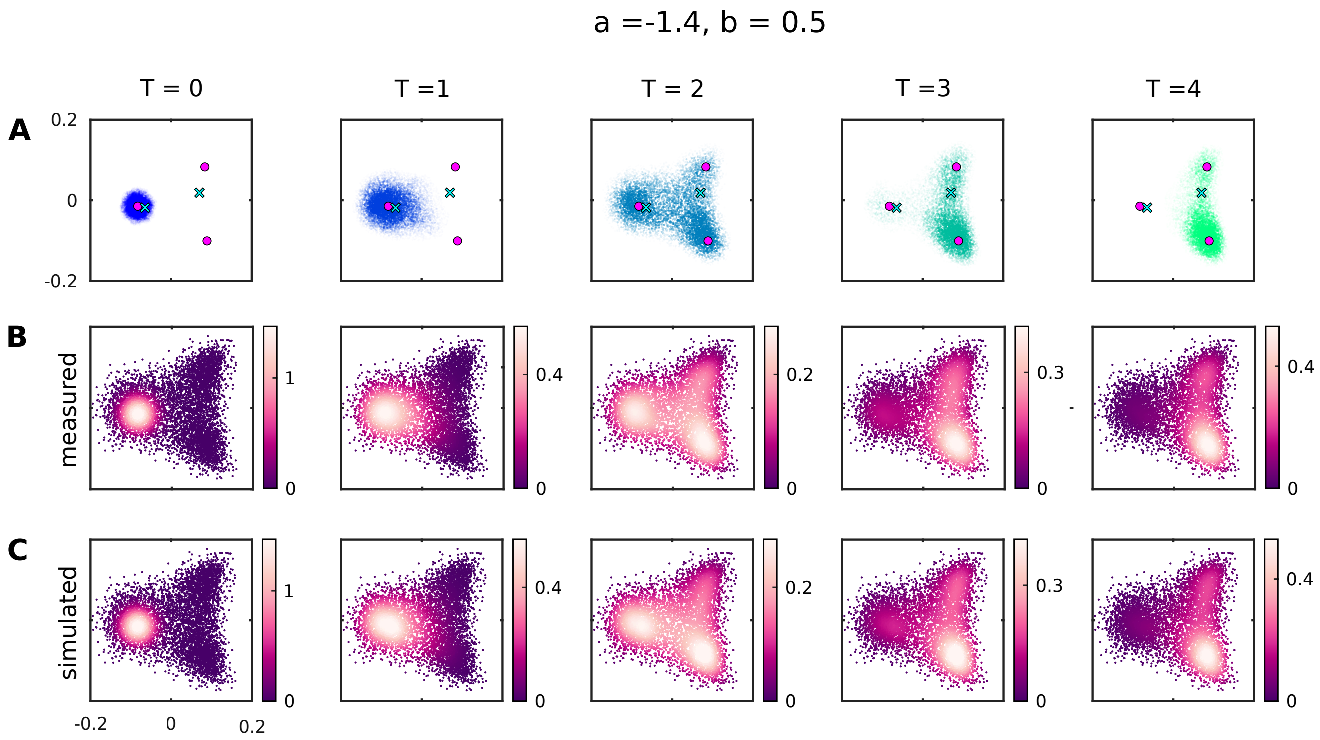

**Figure S4.** Time course of results of the synthetic example discussed in Algorithm Applied To Simulated Data and Figs. 3 and 4 for parameter values $a = -1.4$ and $b = 0.5$. Errors and entropies are reported in Table S1. (A) The actual simulated points at times $T = 0$ to 4 (see Appendix S3.1). Each time point contains 12,000 unique points. (B) The time course of measured densities on $\hat{\mathcal{M}}$ computed using kernel density estimation with the points in (A) as sources. (C) Time course of simulated densities.

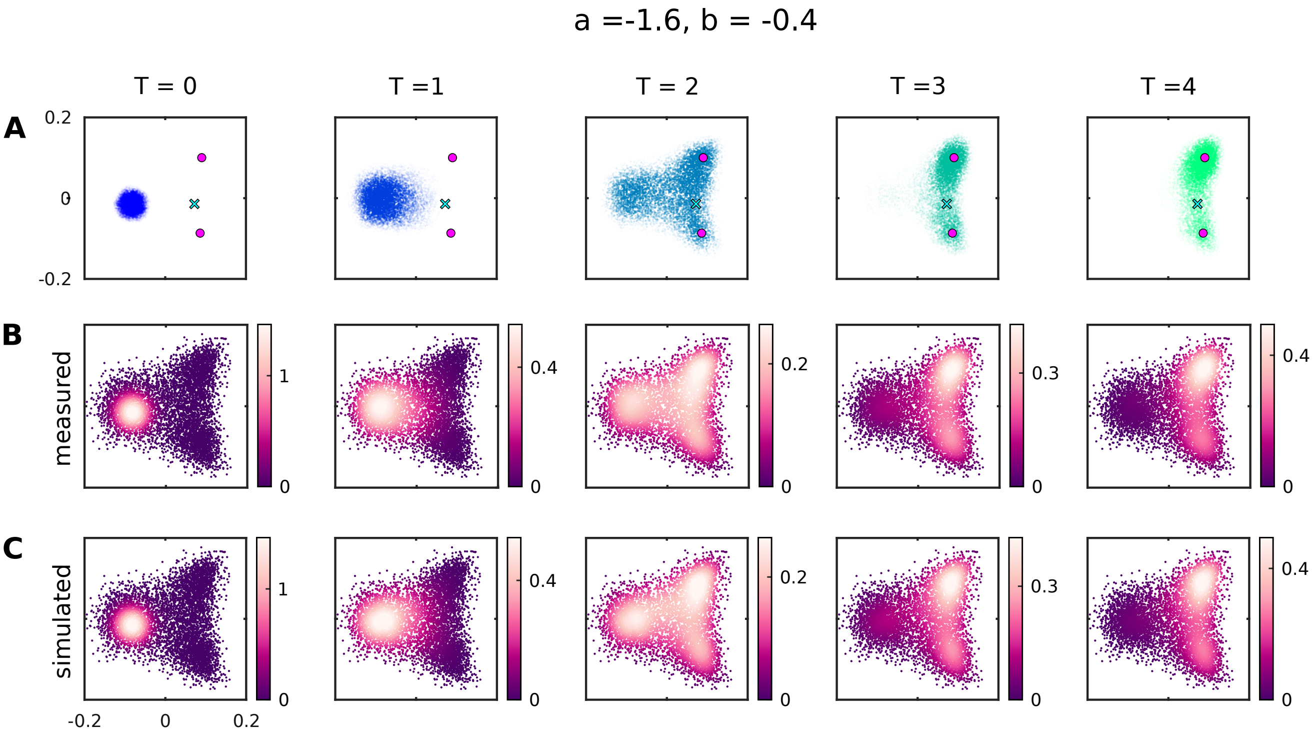

**Figure S5.** Time course of results of the synthetic example discussed in Algorithm Applied To Simulated Data and Figs. 3 and 4 for parameter values $a = -1.6$ and $b = -0.4$. Errors and entropies are reported in Table S1. (A) The actual simulated points at times $T = 0$ to 4 (see Appendix S3.1). Each time point contains 12,000 unique points. (B) The time course of measured densities on $\hat{\mathcal{M}}$ computed using kernel density estimation with the points in (A) as sources. (C) Time course of simulated densities.

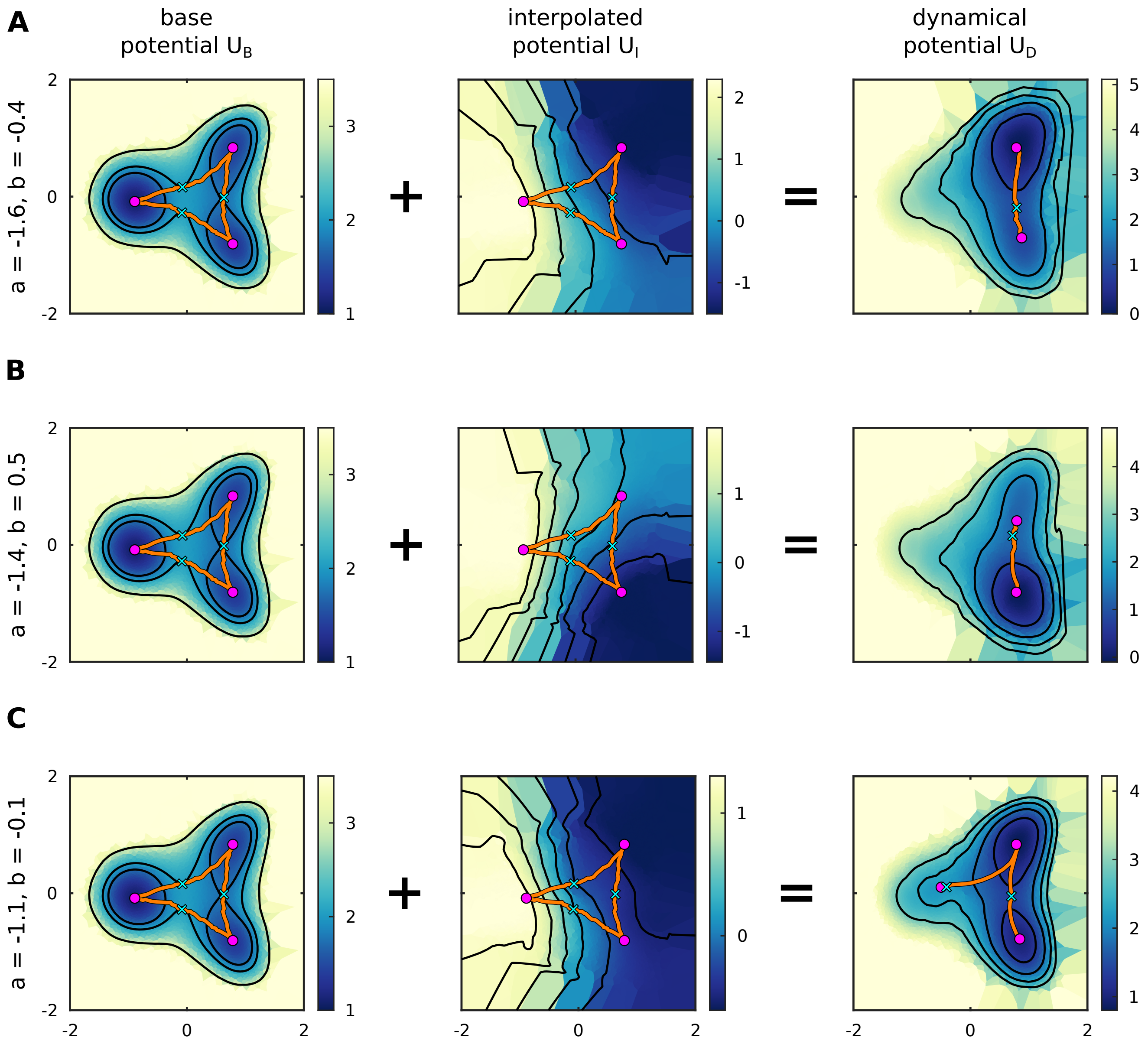

**Figure S6.** Base potential $U_B$, interpolated potential $U_I$, and dynamical potential $U_D$ for each parameter condition of the the synthetic example discussed in Algorithm Applied To Simulated Data and Figs. 3 and 4. For both $U_B$ and $U_I$ the unstable manifolds and fixed points are derived from the base potential. For $U_D$ we use our topological minimum finding techniques to determine the stable attractors and then infer the unstable manifolds between them. See also Fig. 4. (A) Results for $a = -1.6$ and $b = -0.4$. (B) Results for $a = -1.4$ and $b = 0.5$. Notice that $U_D$ does not contain the shallow minimum near $\sim (-1, 0)$ that is present both in the analytic potential $U$ and in the optimized effective potential $U_{eff}$. (C) Results for $a = -1.1$ and $b = -0.1$.

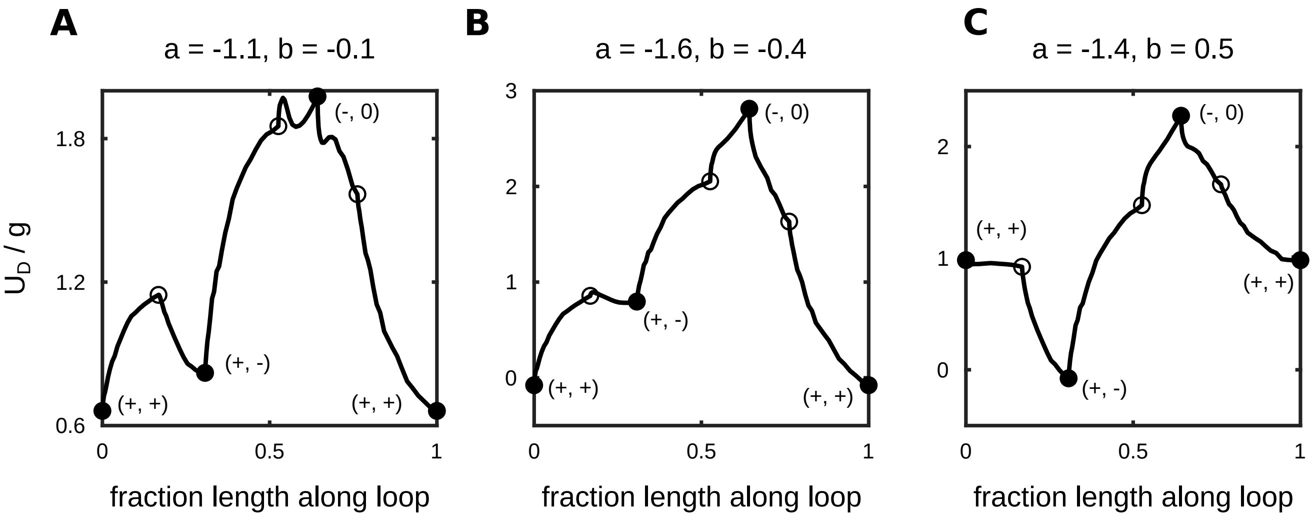

**Figure S7.** Rescaled dynamical potential $U_D/g$ along the loop defined by the base potential unstable manifolds for the synthetic data example in Algorithm Applied To Simulated Data (see Fig. 3B). Filled circles correspond to base potential minima and empty circles correspond to the associated saddles. The symbol $(+,+)$ denotes the minimum near $\sim(1,1)$, $(+,-)$ denotes the minimum near $\sim(1,-1)$, and $(-,0)$ denotes the minimum near $\sim(-1,0)$. (A) Results for $a=-1.1$ and $b=-0.1$. Optimized $g=1.21$. (B) Results for $a=-1.6$ and $b=-0.4$. Optimized $g=1.20$. (C) Results for $a=-1.4$ and $b=0.5$. Optimized $g=1.34$.

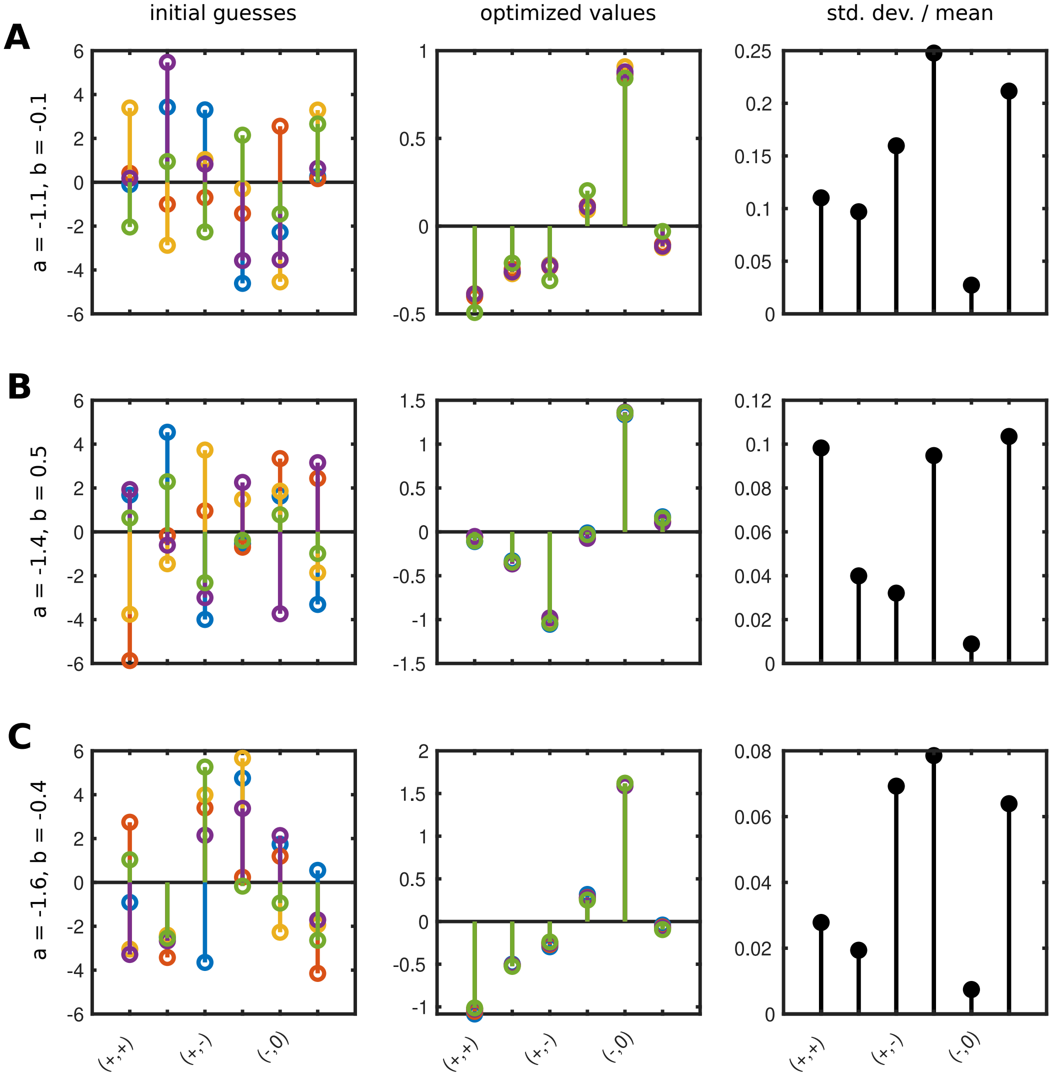

**Figure S8.** Parameter reproducibility for the synthetic data example in Algorithm Applied To Simulated Data. Displayed are the fit heights rescaled by the product of the metric tensor and diffusion coefficient $h/(g\,D)$. Left column shows the initial guesses (five random initial conditions). Note that the initial guesses for $g$ and $D$ are always 1. Middle column shows optimized values. Right column shows the standard deviation of the optimized $h_i/(g\,D)$ divided by $\max\left[\langle h_i/(g\,D)\rangle, \max\left[\langle h_j/(g\,D)\rangle\right]/5\right]$, i.e. standard deviations are rescaled by a minimum of 20% of the maximum mean height. This rescaling reduces irrelevant variability for heights with small absolute values. The symbol $(+,+)$ denotes the minimum near $\sim(1,1)$, $(+,-)$ denotes the minimum near $\sim(1,-1)$, and $(-,0)$ denotes the minimum near $\sim(-1,0)$ (see Fig. 3B). (A) Results for $a=-1.1$ and $b=-0.1$. Optimized $g=1.21$, $D=1.25$. (B) Results for $a=-1.6$ and $b=-0.4$. Optimized $g=1.20$, $D=1.19$. (C) Results for $a=-1.4$ and $b=0.5$. Optimized $g=1.34$, $D=1.09$.

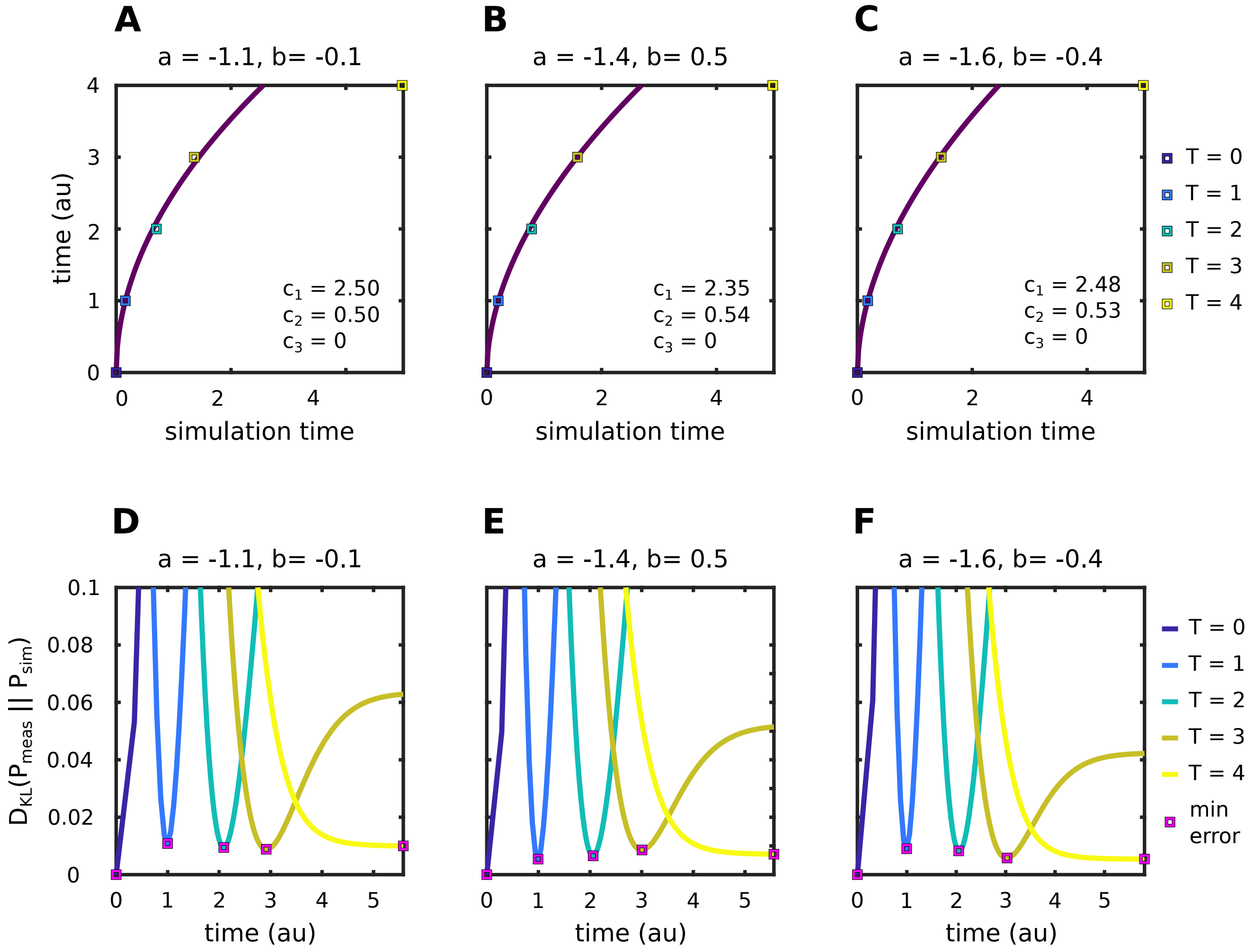

**Figure S9.** Time scales fit to the synthetic data example in Algorithm Applied To Simulated Data (see Appendix S3.8). Top row shows the actual power law that maps simulation time to experimental time ($c_1\, t^{c_2} + c_3$). Bottom row shows the K-L divergence $D_{KL}(P_{meas}||P_{sim})$ for each of the measured probability distributions on $T = 0$ to 4. The magenta squares show the time of minimum error for each data point, which is not necessarily equal to the associated fit time since our power law only has three parameters. Different parameter values produce nearly identical time scale fits, which reflects the fact that the time scales of the simulations that generated the synthetic data were identical for all conditions.

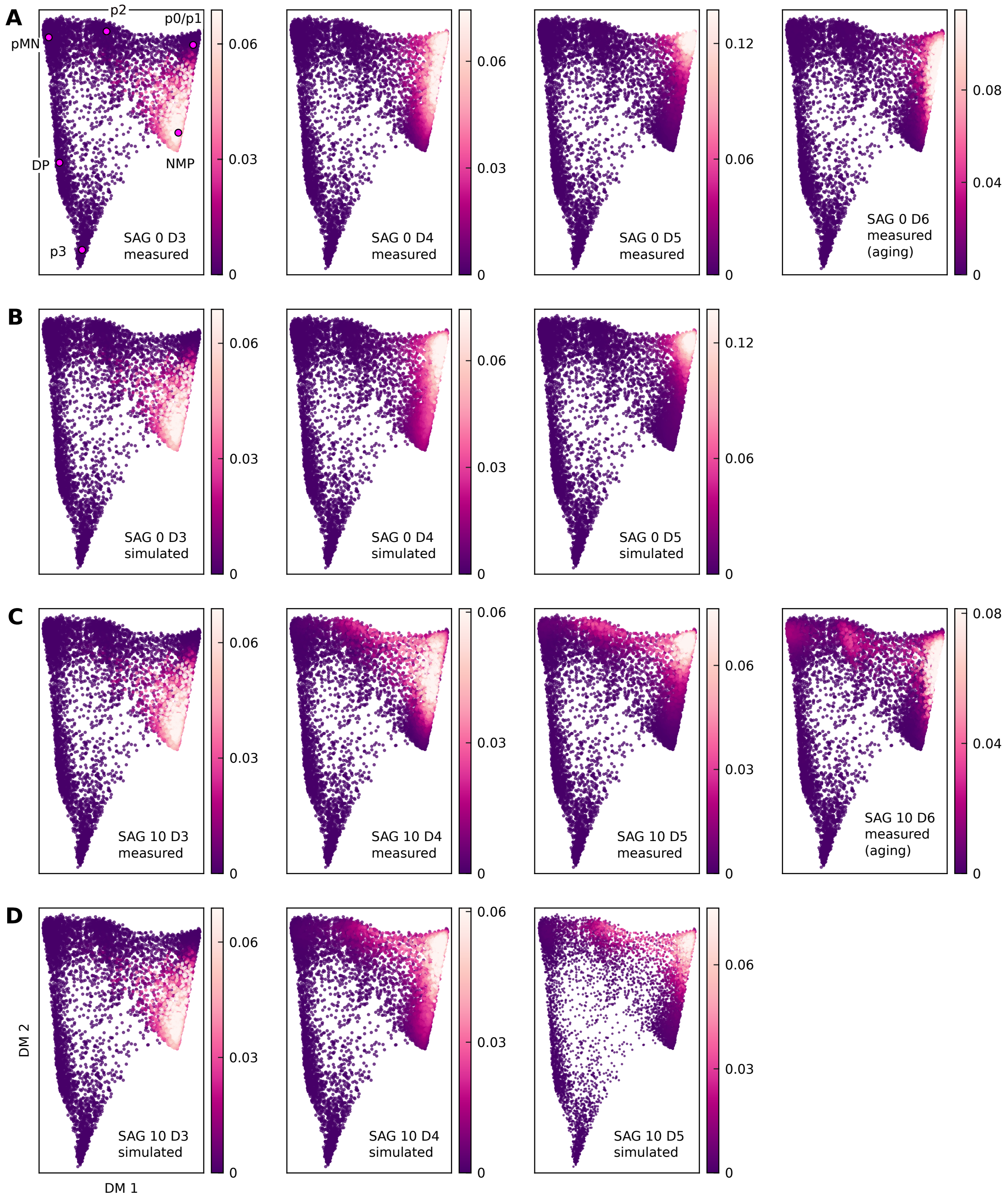

**Figure S10.** Probability density time courses for the constant SAG flow cytometry data. Errors and entropies are reported in Table S3 (A) Measured data for SAG 0 nM. Note the aging effects on Day 6 where probability appears to flow back towards the NMP state. (B) Simulated density time course for SAG 0 nM. Aging effects preclude a fit to Day 6. (C) Measured data for SAG 10 nM. Aging effects are also visible in this data on Day 6. (D) Simulated density time course for SAG 10 nM. Once again, aging effects preclude a fit to Day 6. SAG 10 nM fits shown here were generated using the interpolated scheme (see Appendix S3.9).

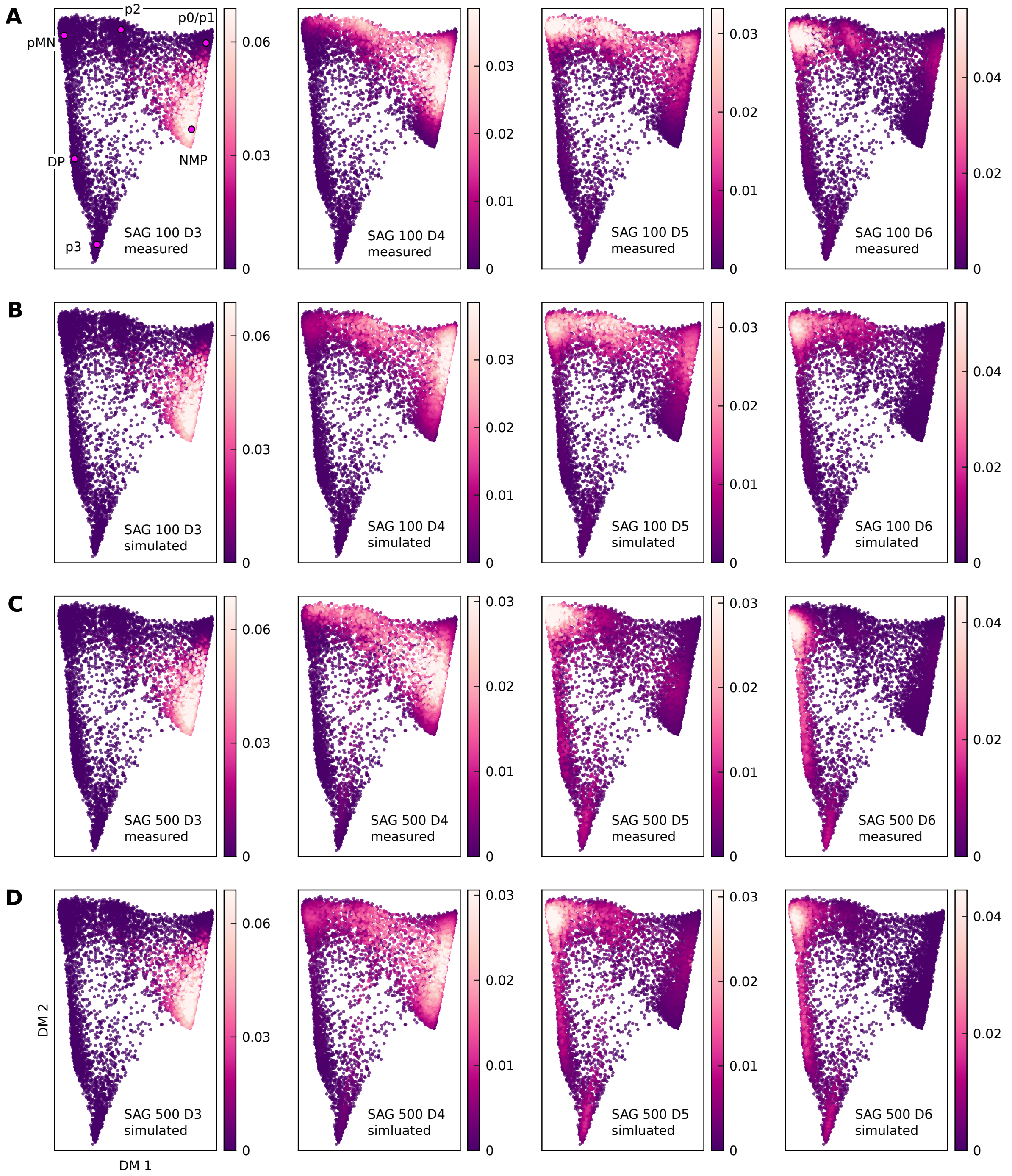

**Figure S11.** Probability density time courses for the constant SAG flow cytometry data. Some aging effects are visible on Day 6 in the measured data for both conditions, but does not preclude a fit. Errors and entropies are reported in Table S3. (A) Measured data for SAG 100 nM. (B) Simulated density time course for SAG 100 nM. SAG 100 nM fits shown here were generated using the interpolated scheme (see Appendix S3.9). (C) Measured data for SAG 500 nM. (D) Simulated density time course for SAG 500 nM. SAG 500 nM fits shown here were generated using the interpolated scheme (see Appendix S3.9).

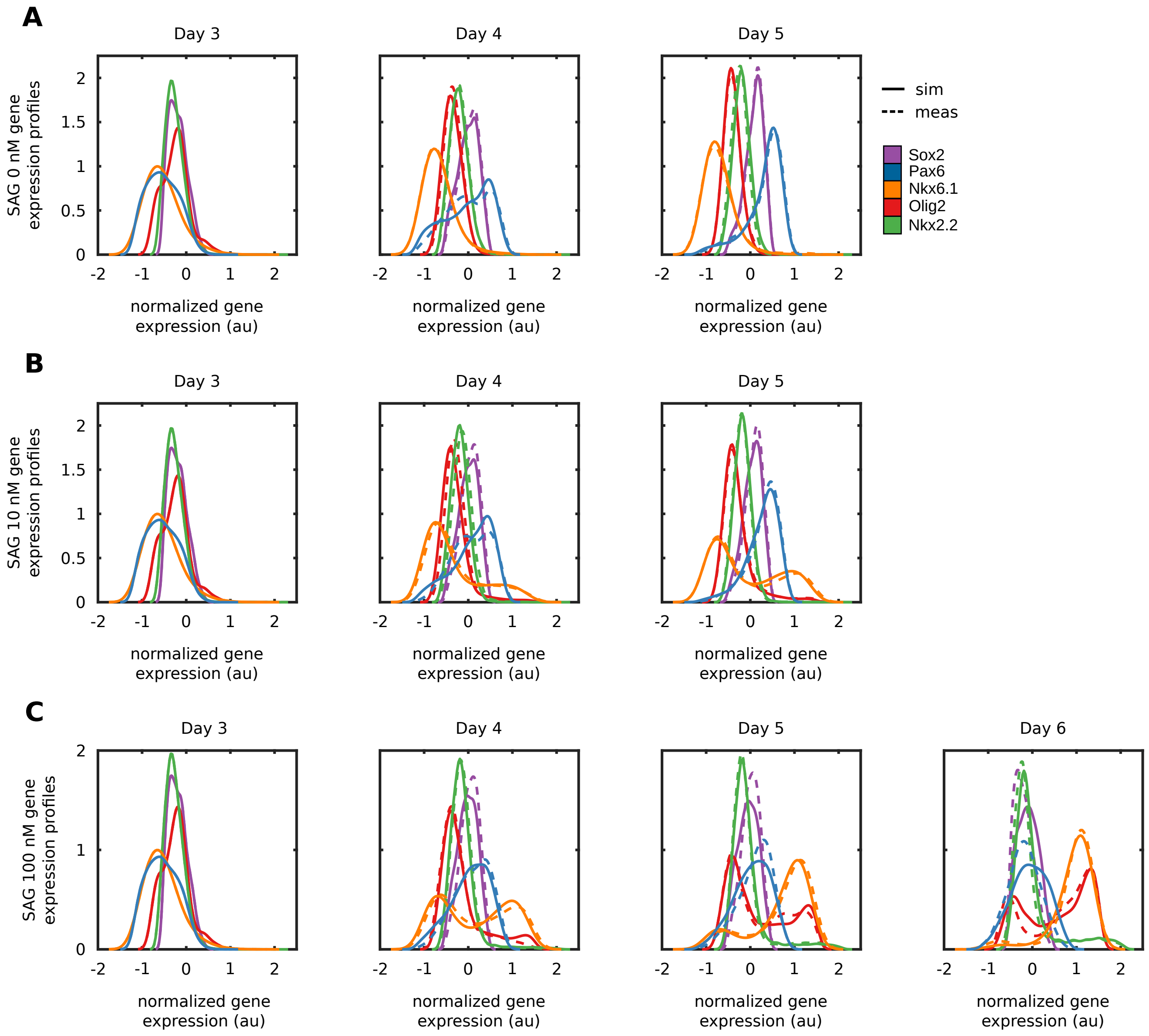

**Figure S12.** Time courses of normalized antibody expression distributions for constant SAG flow cytometry data. Dotted lines denote measured distributions, solid lines denoted simulated distributions. SAG 10 nM and 100 nM fits shown here were generated using the interpolated scheme (see Appendix S3.9). (A) Time course for SAG 0 nM (Day 6 not shown). (B) Time course for SAG 10 nM (Day 6 not shown). (C) Time course for SAG 100 nM.

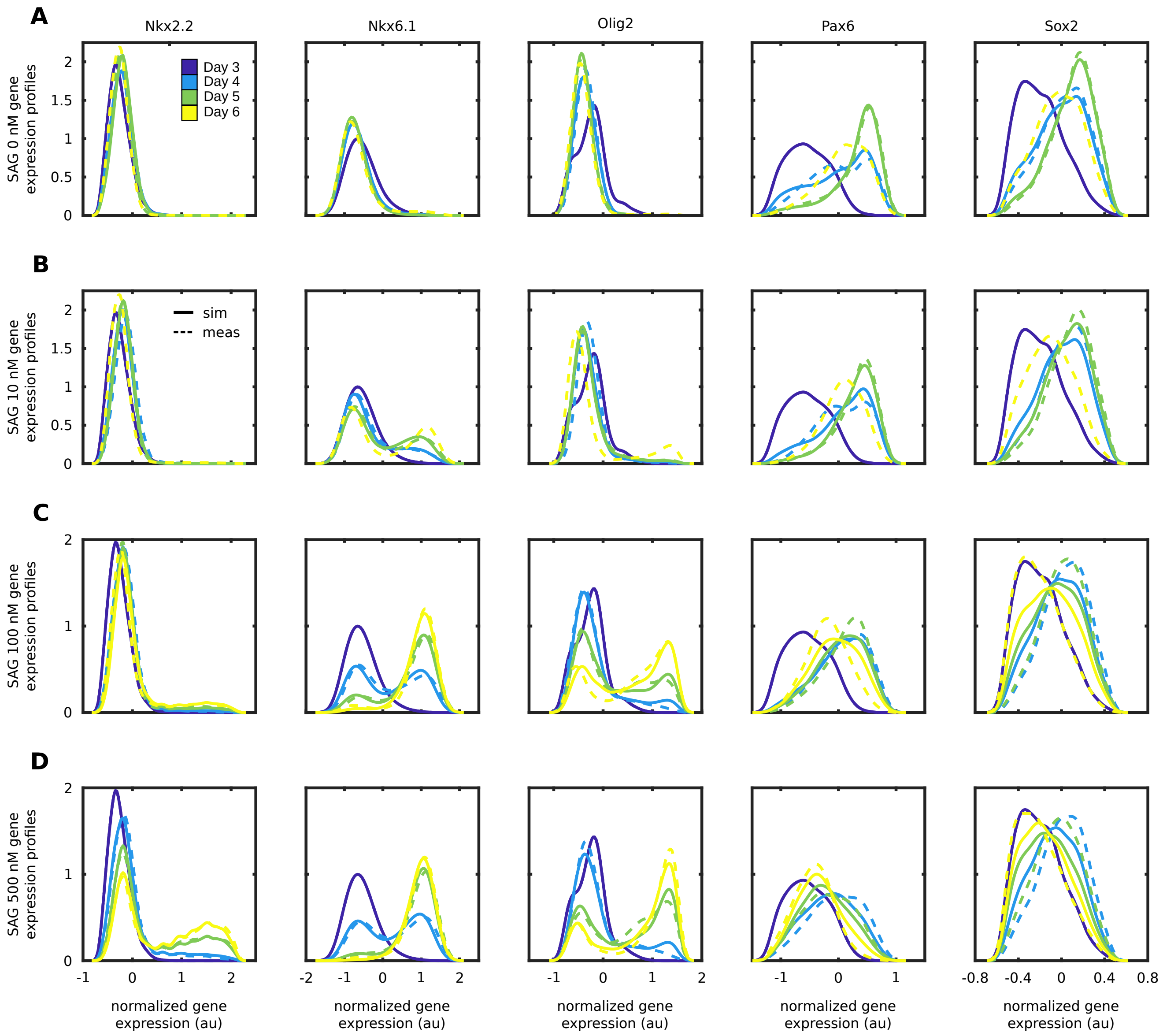

**Figure S13.** Time courses of normalized antibody expression distributions for constant SAG flow cytometry data, with each panel containing the full time course for a given marker and SAG concentration. Dotted lines denote measured distributions, solid lines denoted simulated distributions. SAG 10 nM, 100 nM, and 500 nM fits shown here were generated using the interpolated scheme (see Appendix S3.9). (A) Data for SAG 0 nM. Only measured data is shown on Day 6 since aging effects preclude a fit. (B) Data for SAG 10 nM. Only measured data is shown on Day 6 since aging effects preclude a fit. (C) Data for SAG 100 nM. (D) Data for SAG 500 nM.

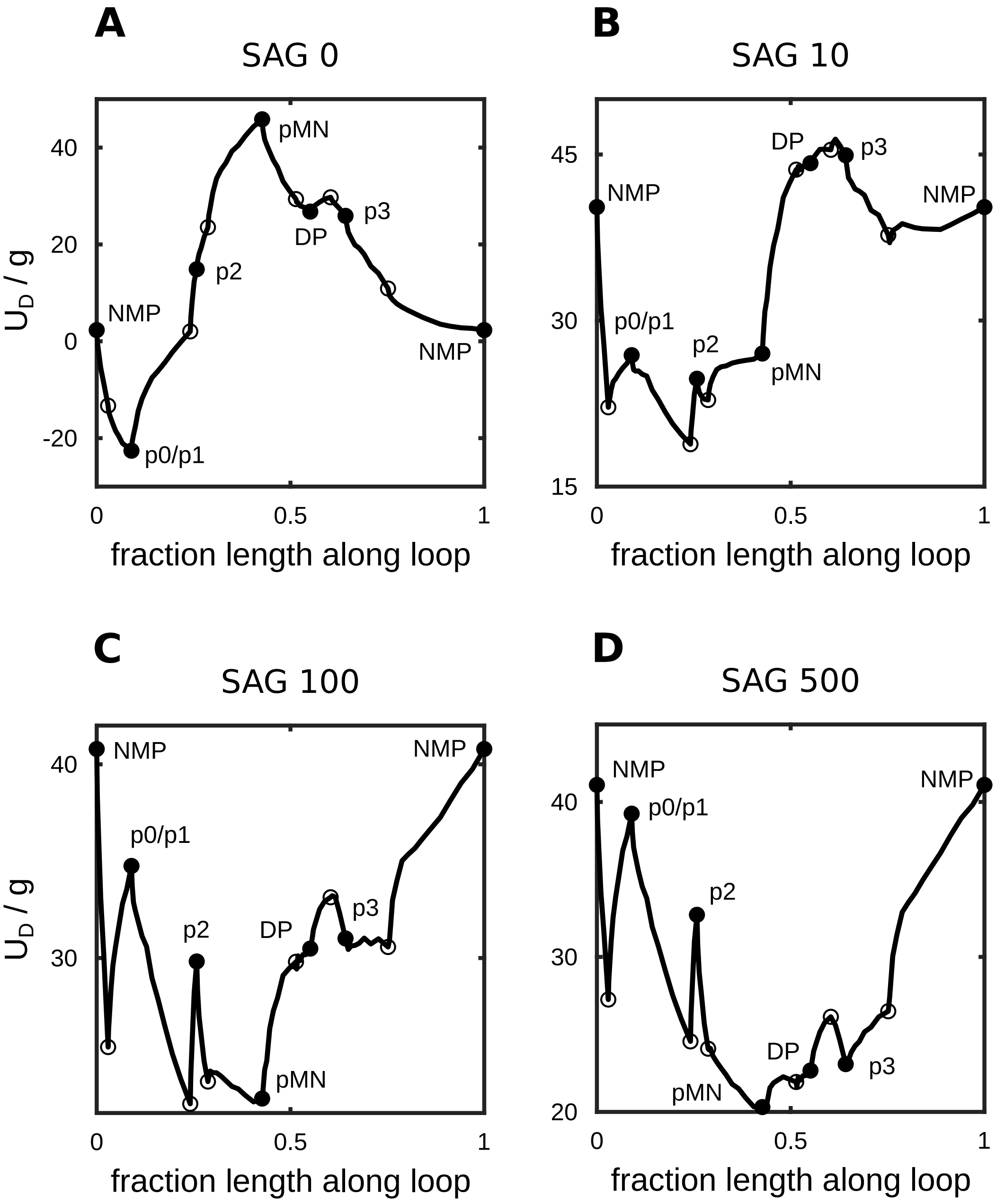

**Figure S14.** Rescaled dynamical potential $U_D/g$ along the loop defined by the base potential unstable manifolds for the flow cytometry data manifold (see Fig. 5C). Filled circles correspond to base potential minima and empty circles correspond to the associated saddles. SAG 10 nM, 100 nM, and 500 nM fits shown here were generated using the interpolated scheme (see Appendix S3.9). (A) Results for SAG 0 nM. Optimized $g^{(0)} = 0.75$. (B) Results for SAG 10 nM. Optimized $g^{(10)} = 0.34$. (C) Results for SAG 100 nM. Recall that $U_D^{(100)}$ is interpolated between $U_D^{(10)}/g^{(10)}$ and $U_D^{(500)}/g^{(500)}$ and that we define $g^{(100)} = 1$ (see Eq. (12) in the main text). (D) Results for SAG 500 nM. Optimized $g^{(500)} = 0.40$

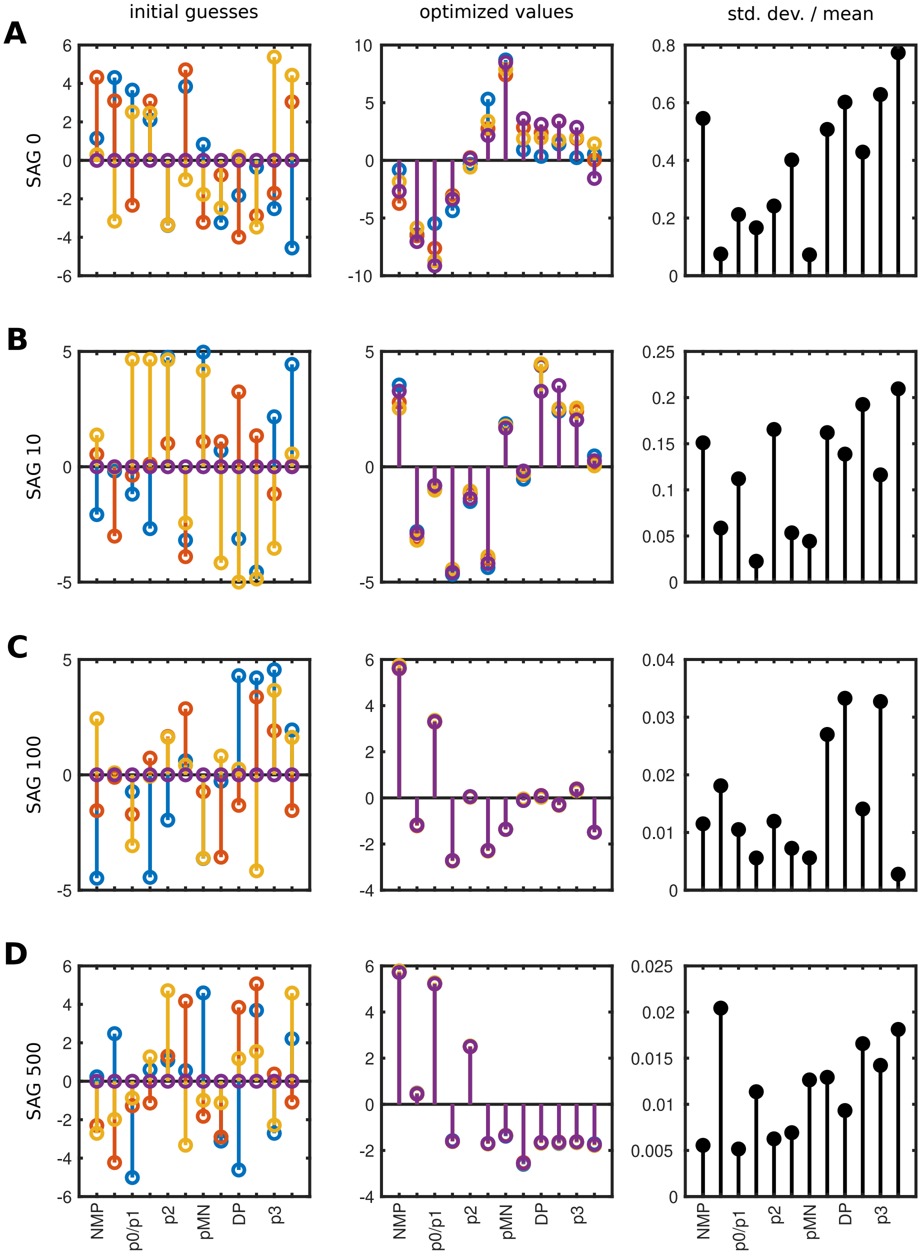

**Figure S15.** Parameter reproducibility for stand alone fits to constant SAG flow cytometry data. Displayed are the fit heights rescaled by the product of the metric tensor and diffusion coefficient $h/(g\,D)$. Left column shows the initial guesses (three random initial conditions and a homogeneous initial condition with all heights equal to zero). Note that the initial guesses for $g$ and $D$ are always 1. Middle column shows optimized values. Right column shows the standard deviation of the optimized $h_i/(g\,D)$ divided by $\max\left[\langle h_i/(g\,D)\rangle, \max\left[\langle h_j/(g\,D)\rangle\right]/5\right]$, i.e. standard deviations are rescaled by a minimum of 20% of the maximum mean height. This rescaling reduces irrelevant variability for heights with small absolute values. (A) Results for stand alone fits to SAG 0 nM. Optimized $g^{(0)} = 0.75$, $D^{(0)} = 3.87$. This condition exhibits the greatest variability reflecting its unimodal population around p0/p1. In other words, the precise heights do not matter so long as the vast majority of the probability ends up in the p0/p1 basin. (B) Results for stand alone fits to SAG 10 nM. Optimized $g^{(10)} = 0.27$, $D^{(10)} = 4.00$. (C) Results for stand alone fits to SAG 100 nM. Optimized $g^{(100)} = 0.28$, $D^{(100)} = 3.28$. (D) Results for stand alone fits to SAG 500 nM. Optimized $g^{(500)} = 0.59$, $D^{(500)} = 2.25$

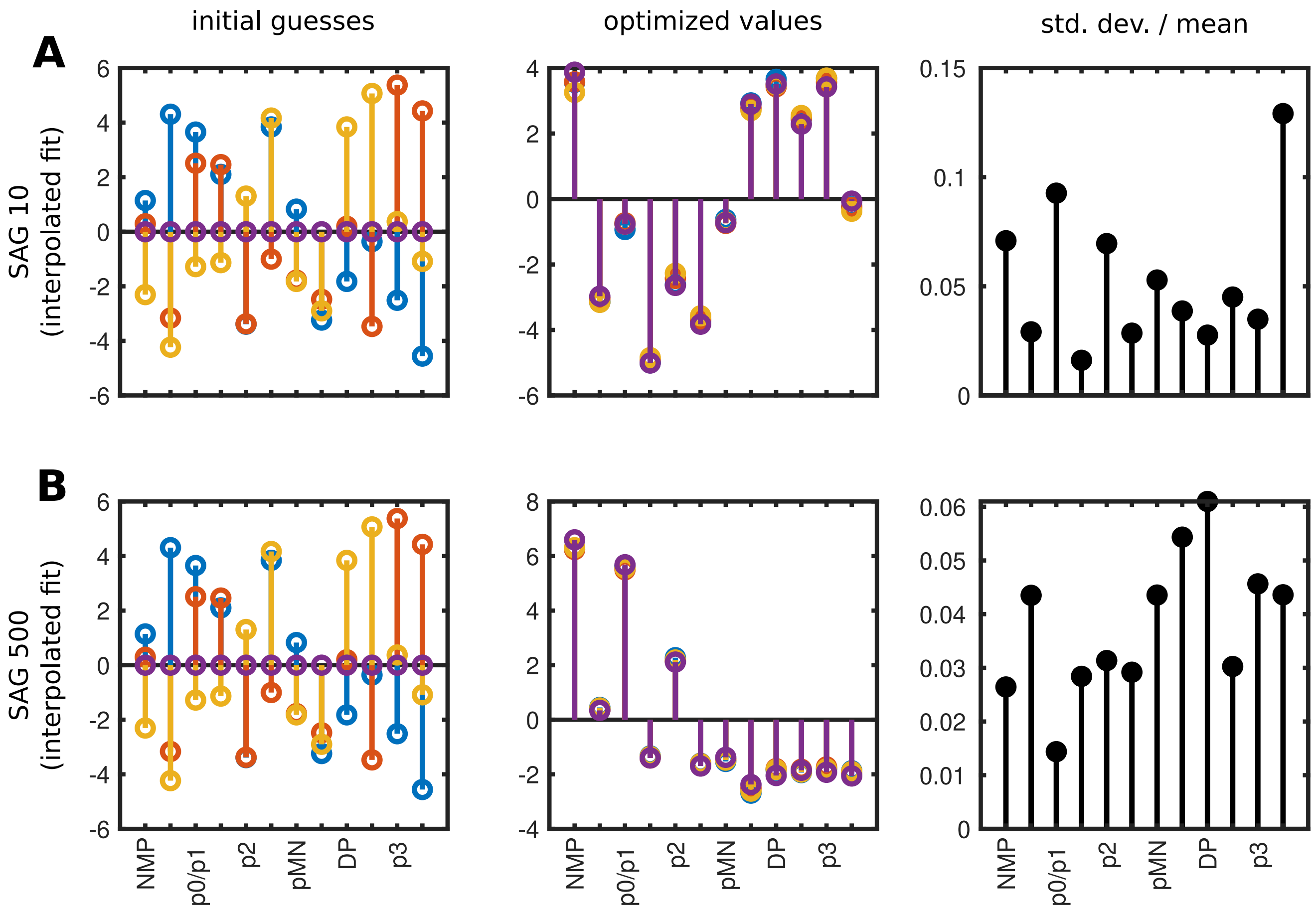

**Figure S16.** Parameter reproducibility for interpolated fits to constant SAG flow cytometry data. Displayed are the fit heights rescaled by the product of the metric tensor and diffusion coefficient $h/(g\,D)$. Left column shows the initial guesses (three random initial conditions and a homogeneous initial condition with all heights equal to zero). Note that the initial guesses for $g$ and $D$ are always 1. Middle column shows optimized values. Right column shows the standard deviation of the optimized $h_i/(g\,D)$ divided by max $\left[\langle h_i/(g\,D)\rangle, \max\left[\langle h_j/(g\,D)\rangle\right]/5\right]$, i.e. standard deviations are rescaled by a minimum of 20% of the maximum mean height. This rescaling reduces irrelevant variability for heights with small absolute values. (A) Results for interpolated fits to SAG 10 nM. Optimized $g^{(10)} = 0.34, D^{(10)} = 3.16$. (B) Results for interpolated fits to SAG 500 nM. Optimized $g^{(500)} = 0.40, D^{(500)} = 2.75$. Interestingly, the parameter values for SAG 500 nM are nearly identical to their counterparts in the SAG 500 nM stand alone fits. The values for SAG 10 nM are similar to their counterparts in the stand alone fits, but exhibit less variability.

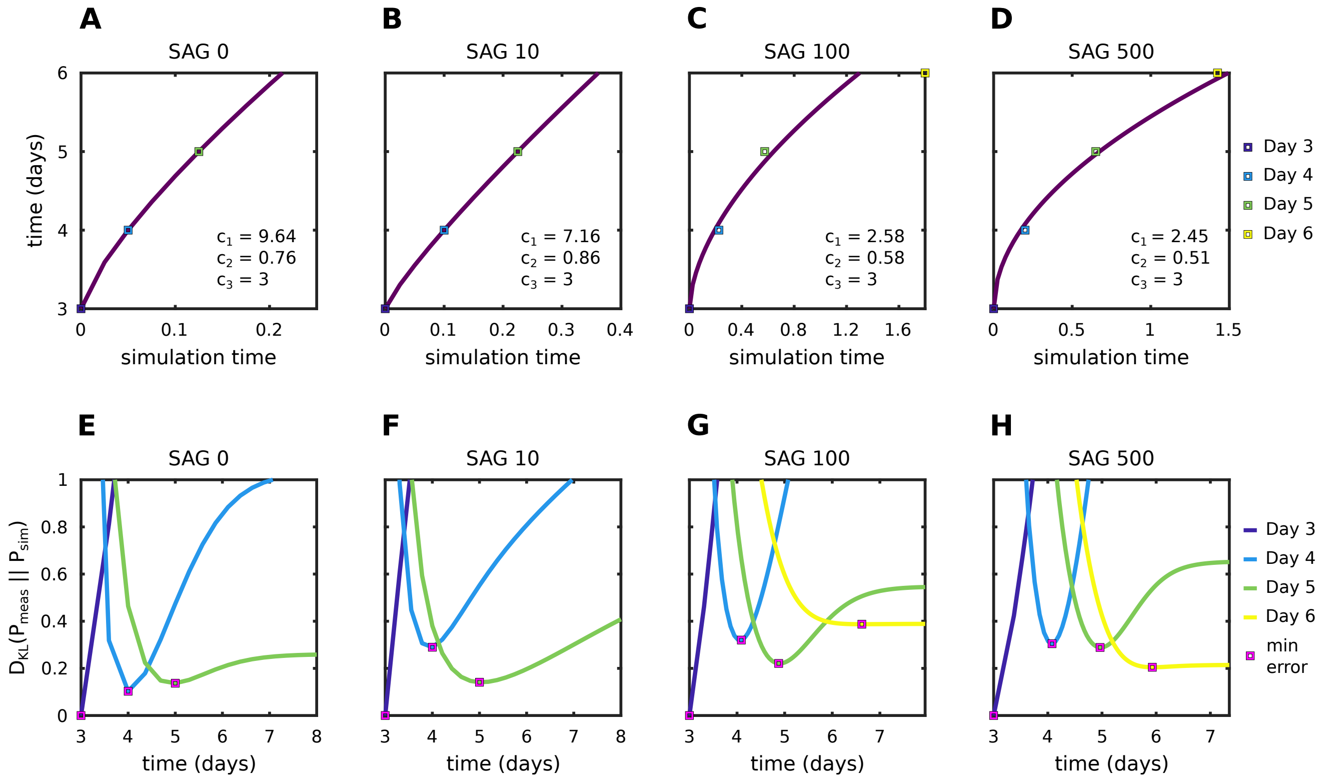

**Figure S17.** Time scales fit to the constant SAG flow cytometry data (see Appendix S3.8). Top row shows the actual power law that maps simulation time to experimental time ($c_1\, t^{c_2} + c_3$). Bottom row shows the K-L divergence $D_{KL}(P_{meas}||P_{sim})$ for each of the measured probability distributions on Days 3 to 6. The magenta squares show the time of minimum error for each data point, which is not necessarily equal to the associated fit time for SAG 100 nM and 500 nM since we fit the times scale for four days and our power law only has three parameters. SAG 10 nM, 100 nM, and 500 nM fits shown here were generated using the interpolated scheme (see Appendix S3.9).

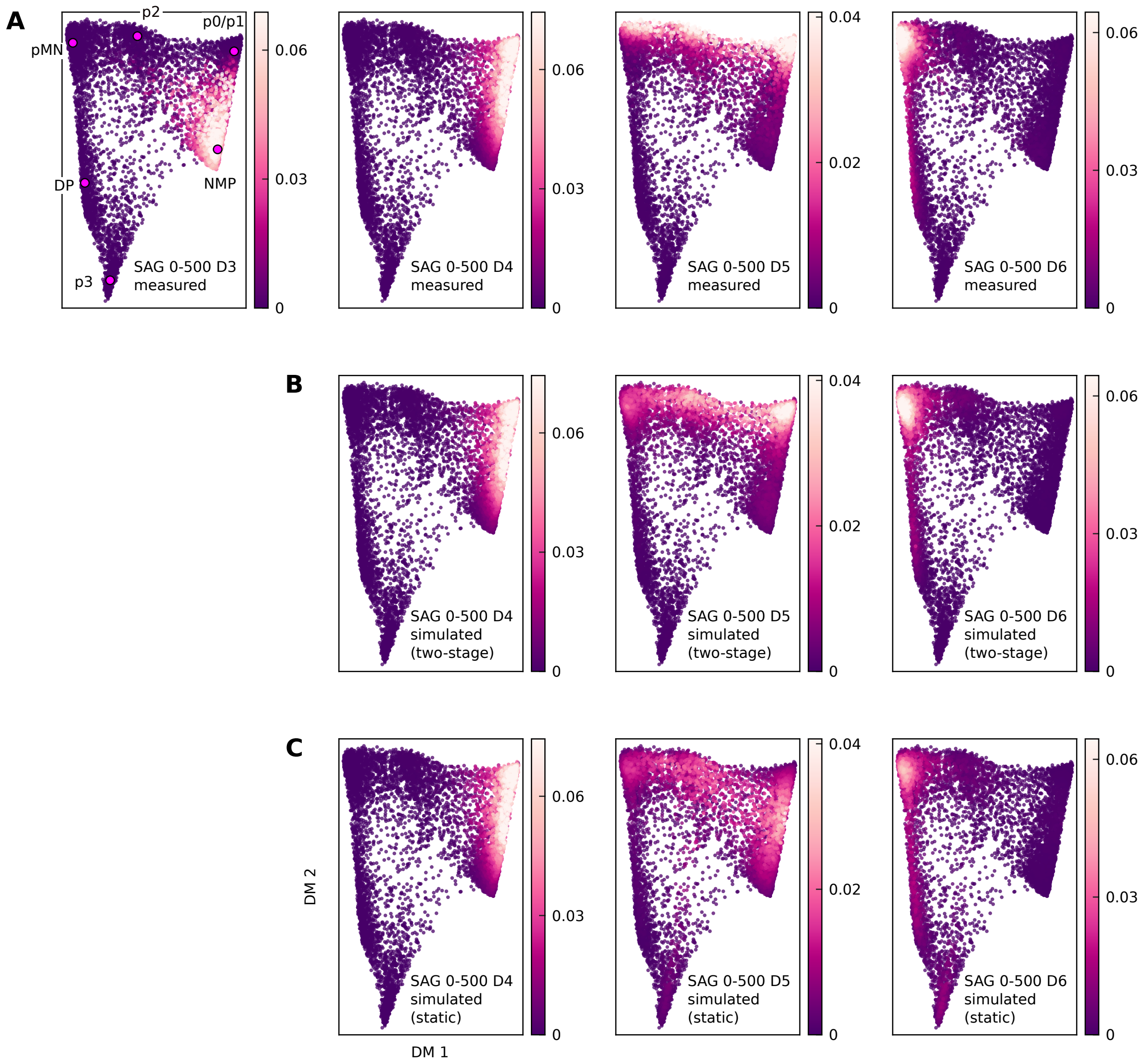

**Figure S18.** Probability density time courses for the SAG 0-500 data. Fits use the Day 4 measured probabilities as an initial condition. Errors and entropies are reported in Table S5. (A) Measured data. (B) Probability density time course for the "two-stage" fit, which is fit directly to the measured data in (A) on Days 4 to 6. (C) Probability density time course for the "static" fit, which simulates a time course using the constant SAG 500 nM landscape using the Day 4 measured probability in (A) as an initial condition.

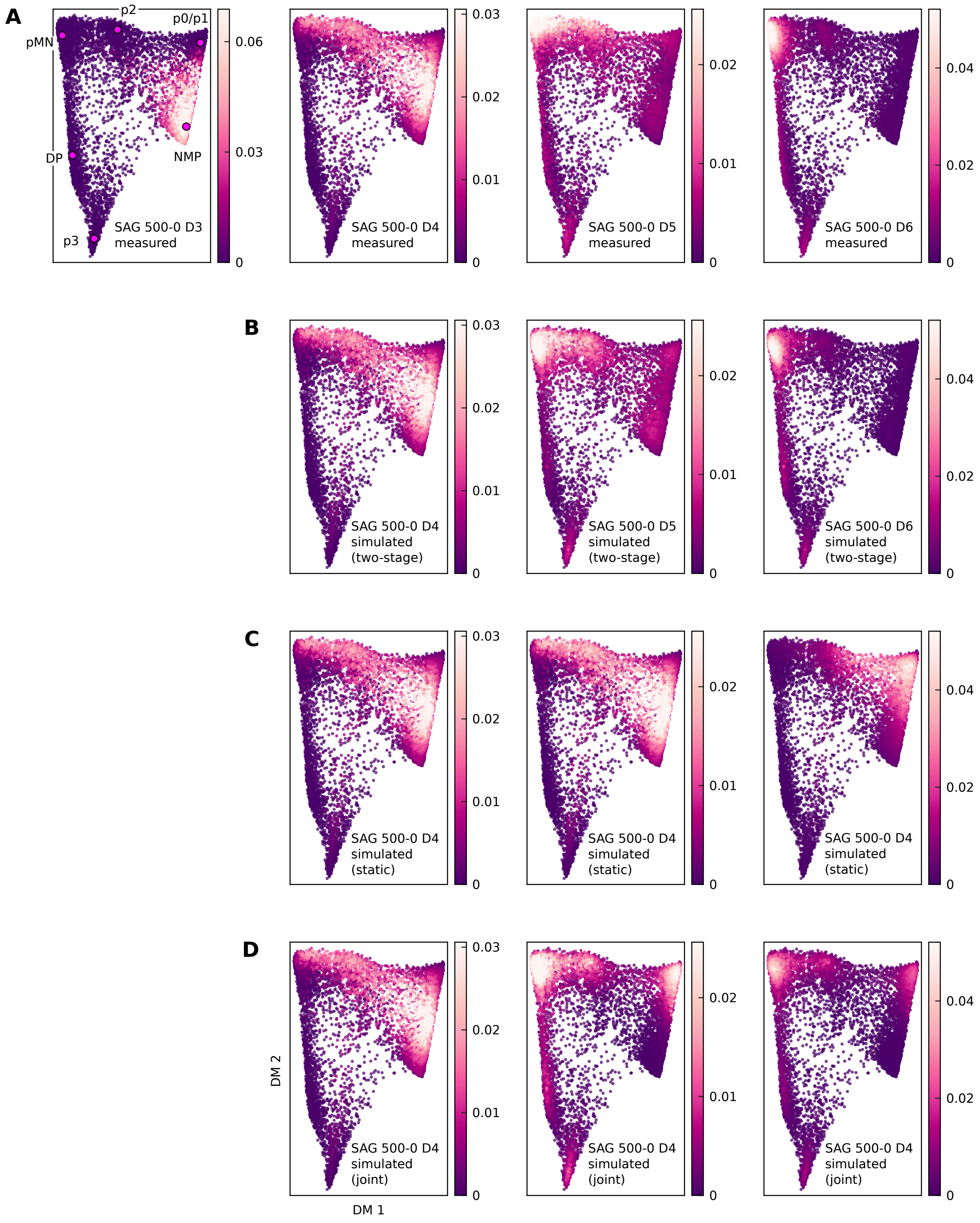

**Figure S19.** Probability density time courses for the SAG 500-0 data. Fits use the Day 4 measured probabilities as an initial condition. Errors and entropies are reported in Table S5. (A) Measured data. (B) Probability density time course for the "two-stage" fit, which is fit directly to the measured data in (A) on Days 4 to 6. (C) Probability density time course for the "static" fit, which simulates a time course using the constant SAG 0 nM landscape using the Day 4 measured probability in (A) as an initial condition. (D) Probability density time course for the "joint" fit, which is fit both to the measured data in (A) and the measured probabilities for the constant SAG 0 nM data (Fig. S10A) on Days 4 to 6.

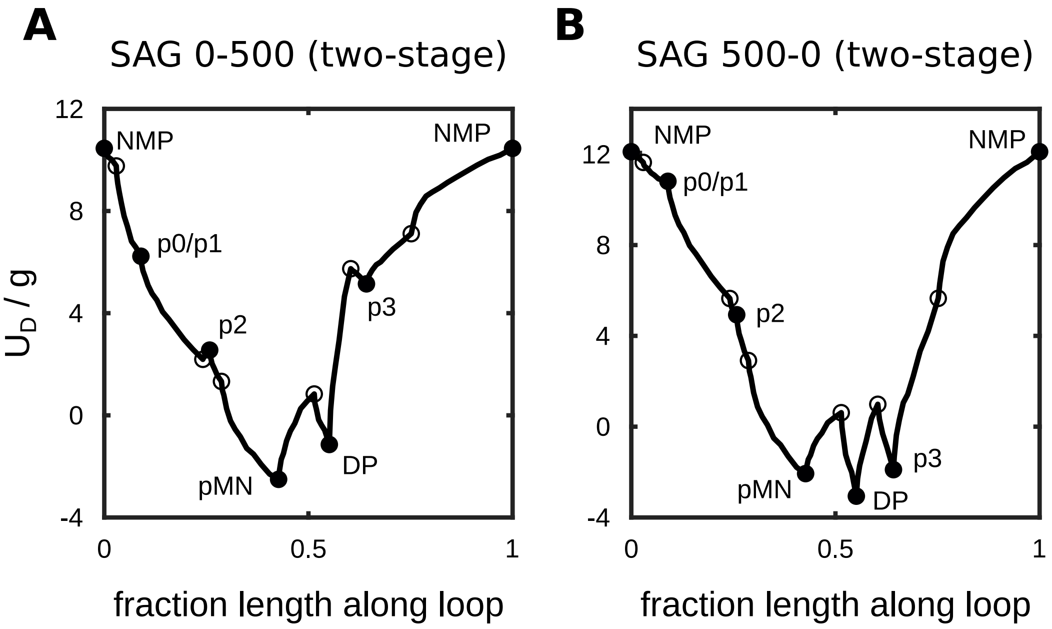

**Figure S20.** Rescaled dynamical potential $U_D/g$ along the loop defined by the base potential unstable manifolds for the flow cytometry data manifold (see Fig. 5C). Filled circles correspond to base potential minima and empty circles correspond to the associated saddles. (A) Results for SAG 0-500. Optimized $g = 2.75$. (B) Results for SAG 500-0. Optimized $g = 2.72$.

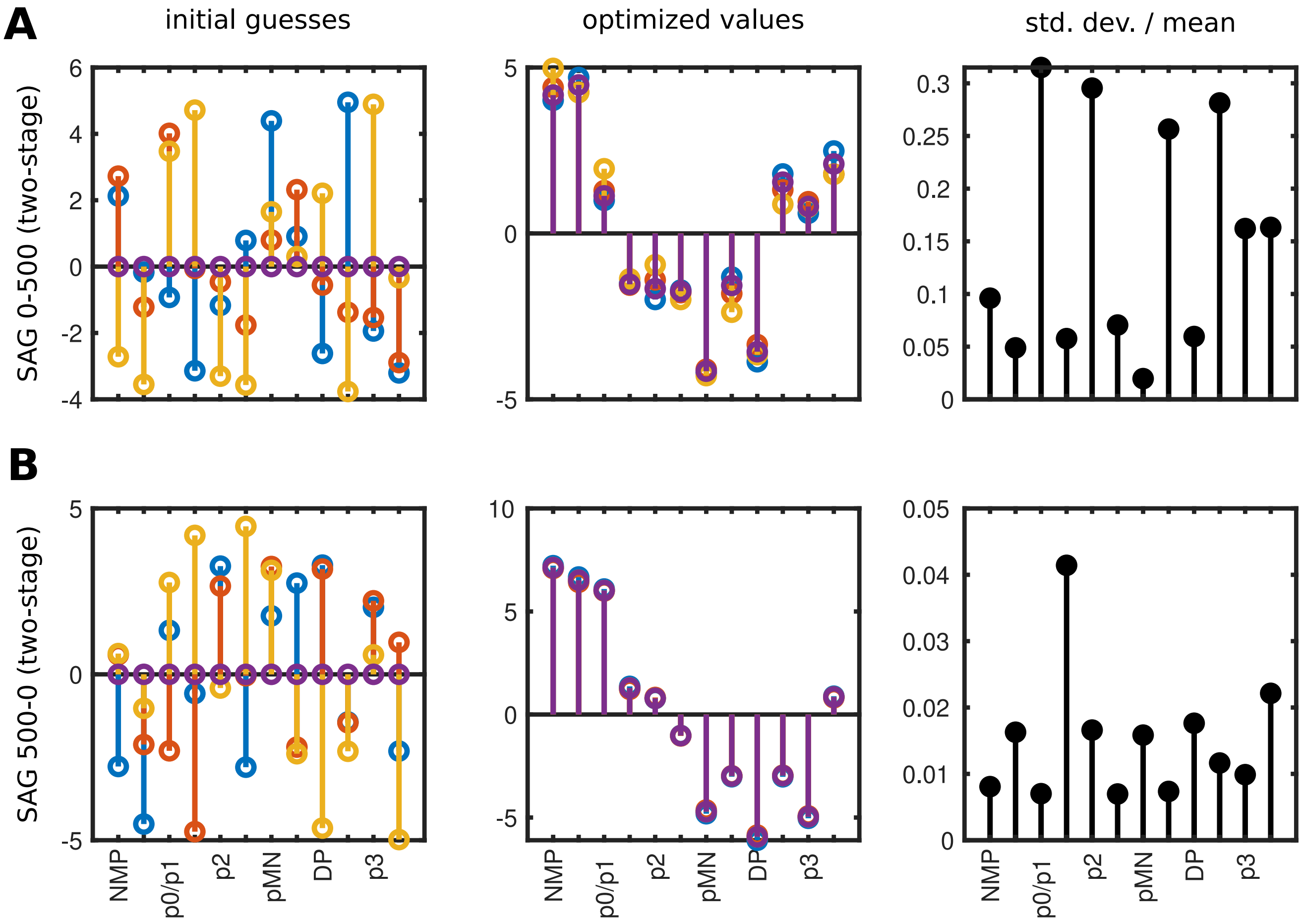

**Figure S21.** Parameter reproducibility for the fits to the SAG 0-500 and 500-0 flow cytometry data. Displayed are the fit heights rescaled by the product of the metric tensor and diffusion coefficient $h/(g\,D)$. Left column shows the initial guesses (three random initial conditions and a homogeneous initial condition with all heights equal to zero). Note that the initial guesses for $g$ and $D$ are always 1. Middle column shows optimized values. Right column shows the standard deviation of the optimized $h_i/(g\,D)$ divided by $\max\left[\langle h_i/(g\,D)\rangle, \max\left[\langle h_j/(g\,D)\rangle\right]/5\right]$, i.e. standard deviations are rescaled by a minimum of 20% of the maximum mean height. This rescaling reduces irrelevant variability for heights with small absolute values. (A) Results for SAG 0-500. Optimized $g = 2.75$, $D = 1.41$. (B) Results for SAG 500-0. Optimized $g = 2.72$, $D = 1.21$.

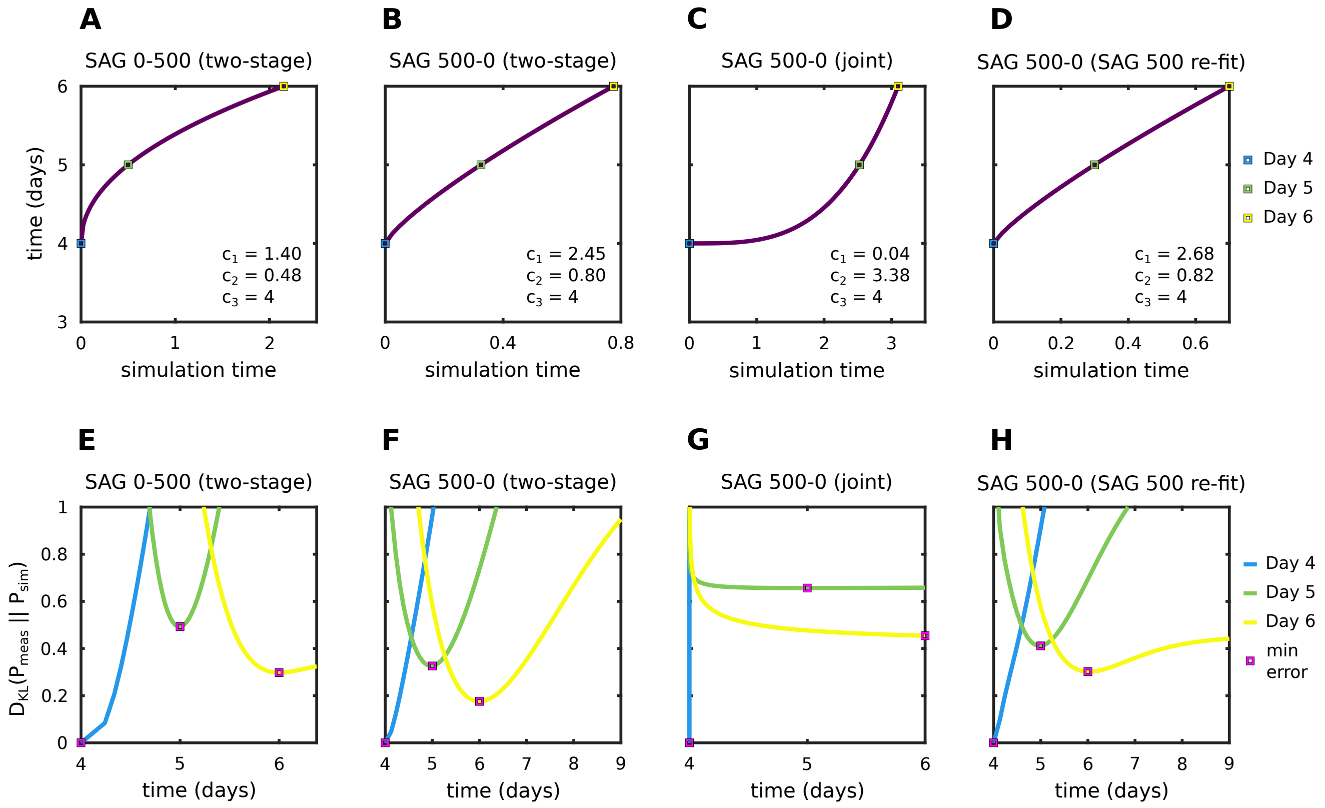

**Figure S22.** Time scales fit to the SAG 0-500 and 500-0 flow cytometry data (see Appendix S3.8). Top row shows the actual power law that maps simulation time to experimental time ($c_1\, t^{c_2} + c_3$). Bottom row shows the K-L divergence $D_{KL}(P_{meas}||P_{sim})$ for each of the measured probability distributions on Days 4 to 6. The magenta squares show the time of minimum error for each data point, which, in this case, is exactly equal to the associated fit time. Notably the parameters found in (D) are slightly greater than, but very close to the constant SAG 500 nM time scale parameters (Fig. S17D).

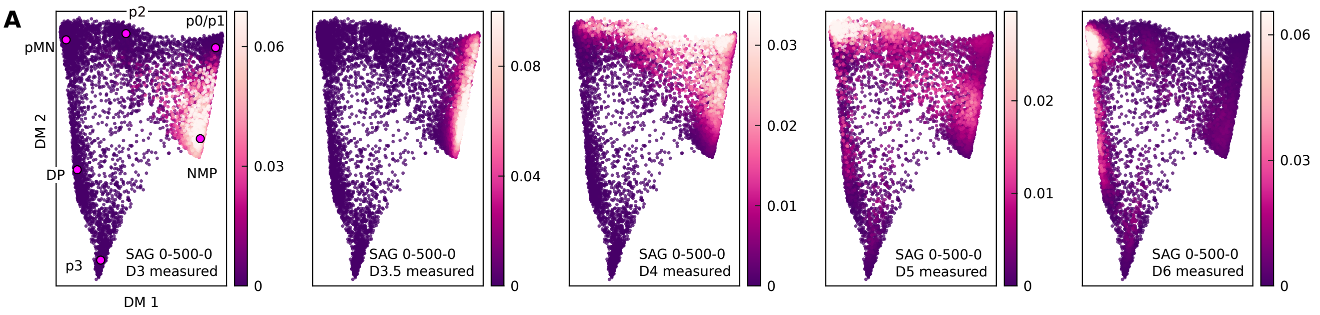

**Figure S23.** Measured probability density time course for the pulsed SAG 0-500-0 flow cytometry data. Crucially, notice that there is virtually zero probability localized near the p0/p1 basin on Day 6.

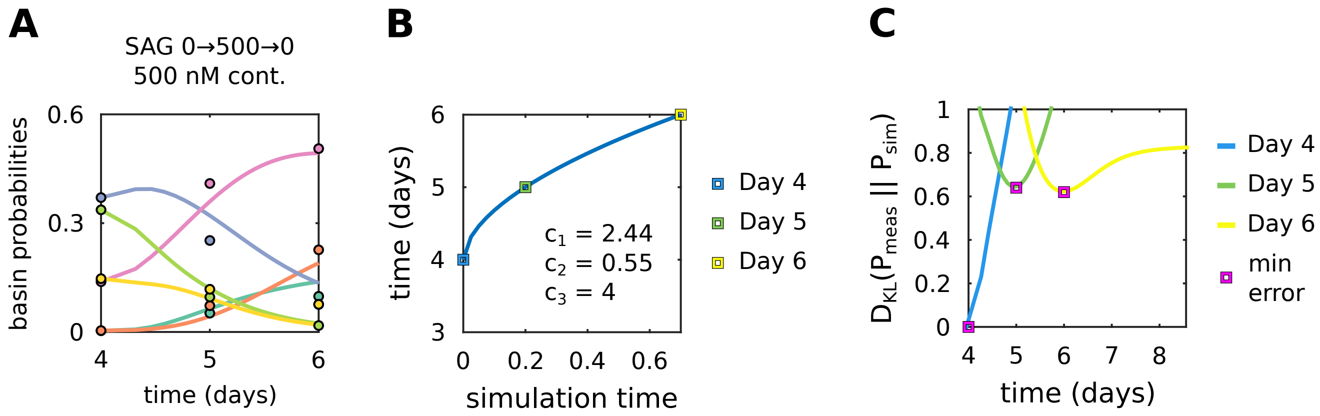

**Figure S24.** The constant SAG 500 nM roughly explains dynamics following withdrawal of SAG in the pulsed SAG 0-500-0 experiment. (A) Point set manifold basin probabilities over time for the pulsed SAG 0-500-0 data beginning on Day 4. Filled circles show measured values and solid lines show the simulated basin probabilities. Simulated values are generated by simulating data in the constant SAG 500 nM landscape using the measured probabilities on Day 4 (Fig. S23, middle panel) as an initial condition and fitting a new time scale. (B) Time scales fit to the pulsed SAG 0-500-0 data (see Appendix S3.8). Notably, the parameters found in nearly identical to the constant SAG 500 nM time scale parameters. (C) K-L divergence $D_{KL}(P_{meas}||P_{sim})$ for each of the measured probability distributions on Days 4 to 6. The magenta squares show the time of minimum error for each data point, which, in this case, is exactly equal to the associated fit time.

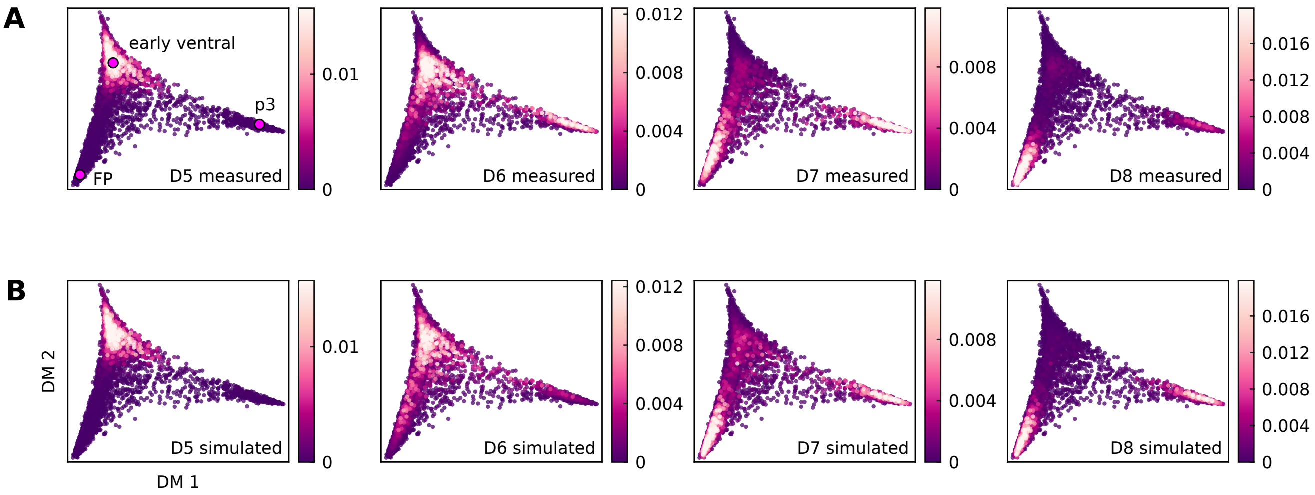

**Figure S25.** Probability density time courses for the SAG 500 nM RNA-seq data. See Fig. 9. Fits use the Day 5 measured probabilities as an initial condition. Errors and entropies are reported in Table S7. (A) Measured data. Notice that the p3 well fills first (Day 6), but mostly empties by Day 8, which indicates a disparity with the notion that this landscape is a single static heteroclinic flip. (B) Simulated probability density time course. The fit produces a static flip-like landscape with the well at p3 being slightly deeper than the well at FP (see Table S8). This compromise to the seemingly time-variable landscape in the measured dynamics results in a slight underfit of the p3 proportion on Day 6 and a slight underfit of FP on Day 8.

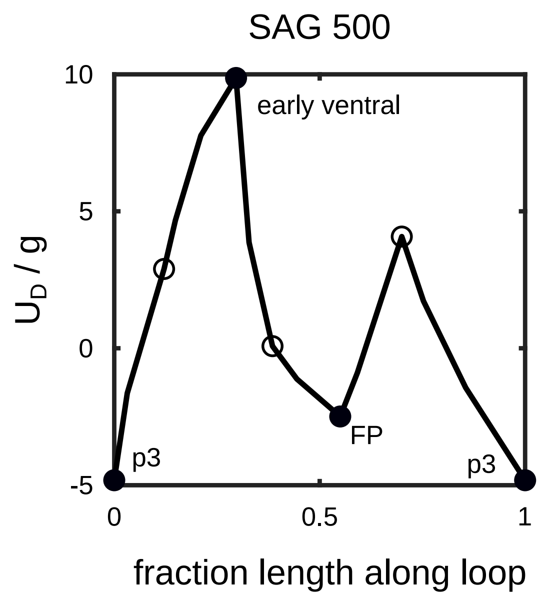

**Figure S26.** Rescaled dynamical potential $U_D/g$ along the loop defined by the base potential unstable manifolds for the RNA-seq data manifold (see Fig. 9B). Filled circles correspond to base potential minima and empty circles correspond to the associated saddles. Optimized $g = 0.52$.

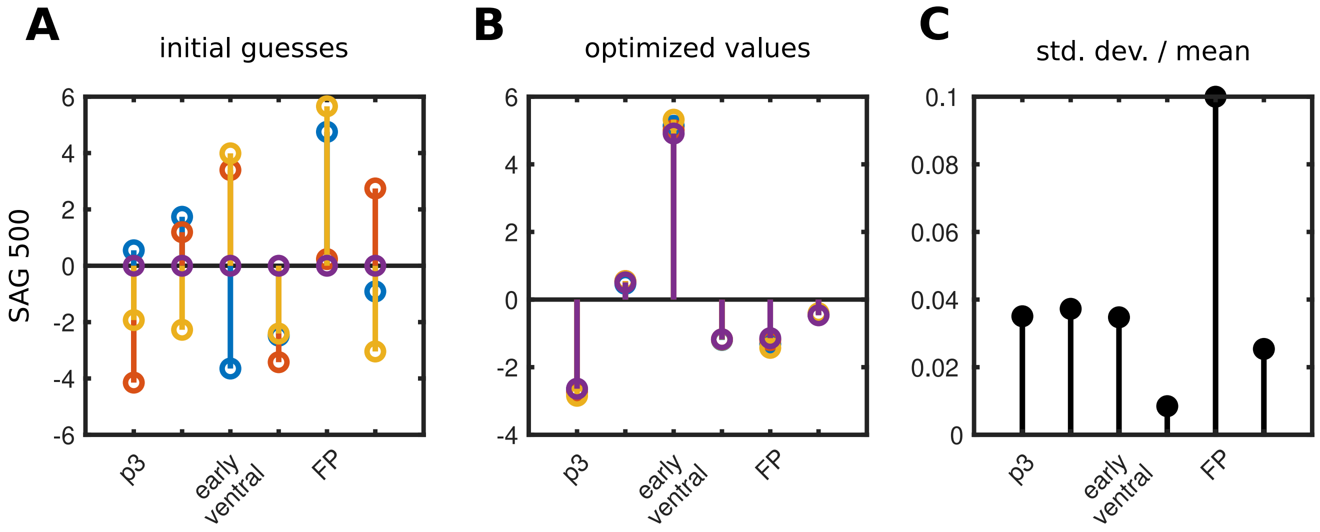

**Figure S27.** Parameter reproducibility for the fits to the SAG 500 nM RNA-seq data. Displayed are the fit heights rescaled by the product of the metric tensor and diffusion coefficient $h/(g\,D)$. Optimized $g = 0.52$, $D = 1.90$. (A) The initial guesses (three random initial conditions and a homogeneous initial condition with all heights equal to zero). Note that the initial guesses for $g$ and $D$ are always 1. (B) Optimized values. (C) Standard deviation of the optimized $h_i/(g\,D)$ divided by $\max\left[\langle h_i/(g\,D)\rangle, \max\left[\langle h_j/(g\,D)\rangle\right]/5\right]$, i.e. standard deviations are rescaled by a minimum of 20% of the maximum mean height. This rescaling reduces irrelevant variability for heights with small absolute values.

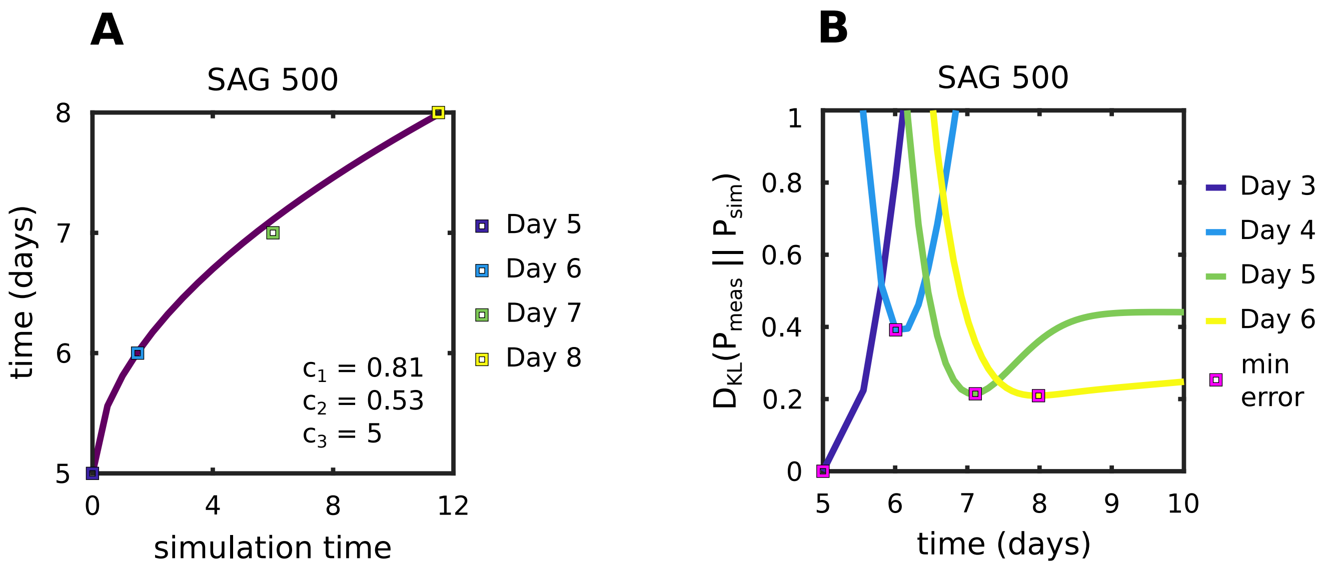

**Figure S28.** Time scales fit to the SAG 500 nM RNA-seq data (see Appendix S3.8). (A) The actual power law that maps simulation time to experimental time ($c_1\, t^{c_2} + c_3$). (B) K-L divergence $D_{KL}(P_{meas}||P_{sim})$ for each of the measured probability distributions on Days 5 to 8. The magenta squares show the time of minimum error for each data point, which is not necessarily equal to the associated fit time since our power law only has three parameters.

| Parameters | $T = 0$ | | $T = 1$ | | $T = 2$ | | $T = 3$ | | $T = 4$ | |
|---|---|---|---|---|---|---|---|---|---|---|
| | $H_{meas}$ | $D_{KL}$ | $H_{meas}$ | $D_{KL}$ | $H_{meas}$ | $D_{KL}$ | $H_{meas}$ | $D_{KL}$ | $H_{meas}$ | $D_{KL}$ |
| $(a = -1.6,\ b = -0.4)$ | 10.909 | — | 11.919 | 0.009 | 12.494 | 0.008 | 12.325 | 0.006 | 12.129 | 0.005 |
| $(a = -1.4,\ b = 0.5)$ | 10.907 | — | 11.891 | 0.005 | 12.501 | 0.007 | 12.371 | 0.009 | 12.176 | 0.007 |
| $(a = -1.1,\ b = -0.1)$ | 10.842 | — | 11.637 | 0.011 | 12.464 | 0.009 | 12.514 | 0.009 | 12.398 | 0.010 |

**Table S1.** Error of optimized fits for the synthetic data example discussed in Algorithm Applied To Simulated Data. $H_{meas}$ denotes the entropy of the measured distributions $P_{meas}$. $D_{KL}$ denotes the (non-symmetric) Kullback-Leibler divergence between $P_{meas}$ and $P_{sim}$, i.e. $D_{KL}(P_{meas}||P_{sim})$. All values are reported in bits. Time points are in arbitrary units. Since the measured distributions at $T = 0$ are used as initial conditions, the associated $D_{KL} = 0$ bits is omitted.

| Parameters | Point | Case | $\lambda_1$ | $\lambda_2$ | Potential | Position $(x, y)$ |
|---|---|---|---|---|---|---|
| $a = -1.60,\ b = -0.40$ | (+,–) | $U_{\text{eff}}$ | -3.91 | -13.26 | -1.11 | (0.86, -0.74) |
| | | $U_{\text{true}}$ | -4.05 | -13.64 | -1.31 | (0.86, -0.87) |
| | X | $U_{\text{eff}}$ | 3.95 | -9.47 | -0.86 | (0.79, -0.19) |
| | | $U_{\text{true}}$ | 2.68 | -8.66 | -1.00 | (0.72, -0.14) |
| | (+,+) | $U_{\text{eff}}$ | -6.24 | -13.23 | -2.07 | (0.83, 0.88) |
| | | $U_{\text{true}}$ | -6.17 | -15.52 | -2.07 | (0.90, 1.00) |
| $a = -1.40,\ b = 0.50$ | (–,0) | $U_{\text{eff}}$ | -1.08 | -6.85 | 0.74 | (-0.57, -0.21) |
| | | $U_{\text{true}}$ | 0.98 | -3.13 | 0.34 | (-0.65, -0.18) |
| | X | $U_{\text{eff}}$ | 0.03 | -6.43 | 0.74 | (-0.51, -0.22) |
| | | $U_{\text{true}}$ | 0.98 | -3.13 | 0.34 | (-0.65, -0.18) |
| | (+,–) | $U_{\text{eff}}$ | -7.58 | -13.09 | -1.99 | (0.83, -0.85) |
| | | $U_{\text{true}}$ | -6.19 | -15.30 | -1.99 | (0.89, -1.01) |
| | X | $U_{\text{eff}}$ | 2.29 | -9.51 | -0.57 | (0.74, 0.16) |
| | | $U_{\text{true}}$ | 2.45 | -8.21 | -0.84 | (0.70, 0.19) |
| | (+,+) | $U_{\text{eff}}$ | -1.43 | -13.10 | -0.64 | (0.83, 0.70) |
| | | $U_{\text{true}}$ | -3.50 | -12.81 | -1.06 | (0.84, 0.83) |
| $a = -1.10,\ b = -0.10$ | (–,0) | $U_{\text{eff}}$ | -3.98 | -6.68 | 0.05 | (-0.74, -0.03) |
| | | $U_{\text{true}}$ | -3.50 | -3.93 | 0.10 | (-0.97, 0.03) |
| | X | $U_{\text{eff}}$ | 3.62 | -4.37 | 0.23 | (-0.41, 0.11) |
| | | $U_{\text{true}}$ | 2.36 | -1.79 | 0.24 | (-0.43, 0.06) |
| | (+,+) | $U_{\text{eff}}$ | -4.55 | -11.86 | -1.34 | (0.79, 0.84) |
| | | $U_{\text{true}}$ | -4.96 | -13.68 | -1.34 | (0.84, 0.93) |
| | X | $U_{\text{eff}}$ | 3.62 | -7.31 | -0.67 | (0.70, -0.09) |
| | | $U_{\text{true}}$ | 2.61 | -7.09 | -0.68 | (0.66, -0.04) |
| | (+,–) | $U_{\text{eff}}$ | -5.13 | -12.76 | -1.17 | (0.84, -0.84) |
| | | $U_{\text{true}}$ | -4.44 | -13.15 | -1.15 | (0.83, -0.90) |

**Table S2.** Hessian eigenvalues ($\lambda_1$, $\lambda_2$), potentials, and fixed-point positions for the synthetic data example discussed in Algorithm Applied To Simulated Data. The moving least-squares method for computing the eigensystem is discussed in Appendix S3.10. See also Fig. 4B. Saddle points are labeled as $X$. The symbol $(+,+)$ denotes the potential minimum near $\sim(1,1)$, $(+,-)$ denotes the potential minimum near $\sim(1,-1)$, and $(-,0)$ denotes the potential minimum near $\sim(-1,0)$. Notice that all potential minima correctly correspond to stable attractors and all saddle points are index-1. We compare the effective potential values, eigenvalues, and inferred fixed point positions to the true values derived from the analytic potential underlying the synthetic example (see Appendix S3.1).

| SAG | Fit Type | Day 3 | | Day 4 | | Day 5 | | Day 6 | |
|---|---|---|---|---|---|---|---|---|---|
| | | $H_{meas}$ | $D_{KL}$ | $H_{meas}$ | $D_{KL}$ | $H_{meas}$ | $D_{KL}$ | $H_{meas}$ | $D_{KL}$ |
| 0 | stand alone | 11.05 | — | 11.39 | 0.10 | 10.91 | 0.14 | 11.12 | — |
| | stand alone, no saddles | | — | | 0.12 | | 0.11 | | — |
| | joint | | — | | 0.30 | | 0.45 | | — |
| 10 | stand alone | 11.05 | — | 11.57 | 0.27 | 11.59 | 0.14 | 11.66 | — |
| | interpolated | | — | | 0.29 | | 0.14 | | — |
| | interpolated, no saddles | | — | | 0.38 | | 0.13 | | — |
| 100 | stand alone | 11.05 | — | 11.79 | 0.33 | 11.90 | 0.23 | 11.88 | 0.36 |
| | interpolated | | — | | 0.32 | | 0.22 | | 0.39 |
| | interpolated, no saddles | | — | | 0.43 | | 0.24 | | 0.39 |
| 500 | stand alone | 11.05 | — | 12.06 | 0.30 | 12.15 | 0.27 | 11.93 | 0.22 |
| | interpolated | | — | | 0.30 | | 0.29 | | 0.20 |
| | interpolated, no saddles | | — | | 0.42 | | 0.32 | | 0.20 |

**Table S3.** Error of optimized fits for the constant SAG flow cytometry data. $H_{meas}$ denotes the entropy of the measured distributions $P_{meas}$. $D_{KL}$ denotes the (non-symmetric) Kullback-Leibler divergence between $P_{meas}$ and $P_{sim}$, i.e. $D_{KL}(P_{meas}||P_{sim})$. All values are reported in bits. Since the measured distributions at Day 3 are used as initial conditions, the associated $D_{KL} = 0$ bits is omitted. $D_{KL}$ are also omitted on Day 6 for SAG 0 nM and SAG 10 nM since aging effects preclude a fit to that time point. The "joint" entry for SAG 0 nM refers to a shared landscape fit simultaneously to SAG 0 nM data (Days 3 to 5) and SAG 500-0 data (Days 4 to 6) (see Table S5)

| SAG | Point | $\lambda_1$ | $\lambda_2$ | $\lambda_3$ | $\lambda_4$ | $\lambda_5$ | $U_{\text{eff}}$ |
|---|---|---|---|---|---|---|---|
| 0 | p0/p1 | -18.02 | -21.69 | -23.32 | -25.39 | -32.81 | 4.38 |
| 10 | p0/p1 | -4.22 | -6.72 | -9.79 | -12.24 | -14.03 | 3.66 |
| | X | 1.11 | -8.76 | -9.67 | -12.24 | -14.68 | 4.01 |
| | p2 | -7.02 | -11.84 | -12.90 | -15.27 | -16.28 | 2.61 |
| 100 | pMN | -13.67 | -21.95 | -27.27 | -30.41 | -35.22 | 8.30 |
| | X | 6.92 | -20.25 | -26.34 | -31.43 | -34.80 | 18.33 |
| | p3 | -8.38 | -16.60 | -20.71 | -26.74 | -30.27 | 15.80 |
| 500 | pMN | -5.66 | -8.66 | -9.35 | -10.88 | -13.26 | 3.34 |
| | X | 5.31 | -1.17 | -7.56 | -9.25 | -12.60 | 5.92 |
| | p3 | -6.41 | -7.45 | -8.01 | -10.03 | -12.23 | 4.52 |

**Table S4.** Hessian eigenvalues ($\lambda_1$ to $\lambda_5$) and effective potential ($U_{\text{eff}}$) at fixed points for the constant SAG flow cytometry data. The moving least-squares method for computing the eigensystem is discussed in Appendix S3.10. See also Fig. 6A. Saddle points are labeled as $X$. Notice that all potential minima correctly correspond to stable attractors and all saddle points are index-1.

| SAG | Fit Type | Day 4 | | Day 5 | | Day 6 | |
|---|---|---|---|---|---|---|---|
| | | $H_{\text{meas}}$ | $D_{KL}$ | $H_{\text{meas}}$ | $D_{KL}$ | $H_{\text{meas}}$ | $D_{KL}$ |
| 0–500 | two-stage | 11.39 | — | 11.64 | 0.49 | 11.67 | 0.30 |
| | static | | — | | 1.22 | | 0.73 |
| 500–0 | two-stage | 12.06 | — | 12.24 | 0.33 | 12.12 | 0.18 |
| | static | | — | | 2.20 | | 5.74 |
| | joint | | — | | 0.66 | | 0.45 |
| | SAG 500 re-fit | | — | | 0.41 | | 0.30 |

**Table S5.** Error of optimized fits for the SAG 0-500 and 500-0 flow cytometry data. $H_{meas}$ denotes the entropy of the measured distributions $P_{meas}$. $D_{KL}$ denotes the (non-symmetric) Kullback-Leibler divergence between $P_{meas}$ and $P_{sim}$, i.e. $D_{KL}(P_{meas}||P_{sim})$. All values are reported in bits. Since the measured distributions at Day 4 are used as initial conditions, the associated $D_{KL} = 0$ bits is omitted. The "joint" entry for SAG 0 nM refers to a shared landscape fit simultaneously to SAG 0 nM data (Days 3 to 5) and SAG 500-0 data (Days 4 to 6) (see Table S3)

| SAG | Point | $\lambda_1$ | $\lambda_2$ | $\lambda_3$ | $\lambda_4$ | $\lambda_5$ | $U_{\text{eff}}$ |
|---|---|---|---|---|---|---|---|
| 0-500 | pMN | -18.12 | -34.08 | -40.97 | -42.67 | -47.42 | 8.78 |
| 500-0 | pMN | -18.20 | -32.69 | -37.05 | -42.14 | -45.91 | 10.61 |
| | X | 27.66 | -18.55 | -21.42 | -35.13 | -37.47 | 15.24 |
| | DP | -19.72 | -28.02 | -29.72 | -35.48 | -42.52 | 8.57 |
| | X | 29.18 | -11.32 | -19.09 | -28.10 | -38.39 | 16.23 |
| | p3 | -22.78 | -25.50 | -33.78 | -35.92 | -42.47 | 11.13 |

**Table S6.** Hessian eigenvalues ($\lambda_1$ to $\lambda_5$) and effective potential ($U_{\text{eff}}$) at fixed points for the SAG 0-500 and 500-0 flow cytometry data. The moving least-squares method for computing the eigensystem is discussed in Appendix S3.10. See also Fig. 7A and D. Saddle points are labeled as $X$. Notice that all potential minima correctly correspond to stable attractors and all saddle points are index-1.

| Condition | Day 5 | | Day 6 | | Day 7 | | Day 8 | |
|---|---|---|---|---|---|---|---|---|
| | $H_{\text{meas}}$ | $D_{KL}$ | $H_{\text{meas}}$ | $D_{KL}$ | $H_{\text{meas}}$ | $D_{KL}$ | $H_{\text{meas}}$ | $D_{KL}$ |
| SAG 500 | 10.203 | — | 10.886 | 0.39 | 10.951 | 0.21 | 10.519 | 0.21 |

**Table S7.** Error of optimized fits for the constant SAG 500 nM RNA-seq data. $H_{meas}$ denotes the entropy of the measured distribution $P_{meas}$. $D_{KL}$ denotes the (non-symmetric) Kullback-Leibler divergence between $P_{meas}$ and $P_{sim}$, i.e. $D_{KL}(P_{meas}||P_{sim})$. All values are reported in bits. Since the measured distribution at Day 5 is used as an initial condition, the associated $D_{KL} = 0$ bits is omitted.

| Condition | Point | $\lambda_1$ | $\lambda_2$ | $\lambda_3$ | $\lambda_4$ | $\lambda_5$ | $\lambda_6$ | $\lambda_7$ | $\lambda_8$ | ··· |
|---|---|---|---|---|---|---|---|---|---|---|
| SAG 500 | FP | -0.13 | -0.32 | -0.38 | -0.42 | -0.46 | -0.50 | -0.53 | -0.56 | |
| | X | 0.87 | -0.21 | -0.30 | -0.31 | -0.38 | -0.40 | -0.44 | -0.50 | |
| | p3 | 0.91 | -0.10 | -0.18 | -0.23 | -0.27 | -0.33 | -0.41 | -0.44 | |

| | $\lambda_9$ | $\lambda_{10}$ | $\lambda_{11}$ | $\lambda_{12}$ | $\lambda_{13}$ | $\lambda_{14}$ | $\lambda_{15}$ | $\lambda_{16}$ | $\lambda_{17}$ | $U_{\text{eff}}$ |
|---|---|---|---|---|---|---|---|---|---|---|
| FP | -0.60 | -0.64 | -0.68 | -0.73 | -0.85 | -0.96 | -1.12 | -1.26 | -3.89 | 3.63 |
| X | -0.51 | -0.56 | -0.57 | -0.65 | -0.73 | -0.74 | -0.82 | -0.90 | -2.87 | 7.31 |
| p3 | -0.48 | -0.55 | -0.55 | -0.66 | -0.69 | -0.78 | -0.84 | -0.90 | -2.03 | 2.83 |

**Table S8.** Hessian eigenvalues ($\lambda_1$ to $\lambda_{17}$) and effective potential ($U_{\text{eff}}$) at fixed points for the SAG 500 nM RNA-seq data. The moving least-squares method for computing the eigensystem is discussed in Appendix S3.10. See also Fig. 9C. Saddle points are labeled as $X$. The potential minima at FP is a stable attractor and the saddle point is index-1. The minimum at p3 has a single positive eigenvalue, however, this point is the absolute minimum of $U_{\text{eff}}$ and is surely an attractor. We note that despite this point being the absolute minimum of the dynamical potential, it has fewer points associated to it in $\hat{\mathcal{M}}$ simply due to variability in sampling on different experimental days. This artifact highlights some of the limitations of moving least-squares methods around sparse points in high dimensions.

**SI Movie 1.** The discrete dynamical manifold subsampled from all experimental conditions in the flow cytometry data discussed in Landscape model for neural tube patterning (see Fig. 5), projected onto the three lowest diffusion map dimensions. Colors denote the basins of attraction for each base density local maxima. It is visually clear that the manifold is roughly two-dimensional.

**SI Movie 2.** The discrete dynamical manifold for the SAG 500 nM RNA seq data (see Fig. 9), projected onto the three lowest diffusion map dimensions. Colors denote the basins of attraction for each base density local maxima.

**SI Movie 3.** Dynamical diffusion map embedding of the discrete manifold for the SAG 500 nM RNA seq data (see Fig. 9F and G). Colors denote the basins of attraction for each base density local maxima.
